# Fixation Before Dissociation Using a Deep Eutectic Solvent Preserves *In Vivo* States and Phospho-Signaling in Single-Cell Sequencing

**DOI:** 10.1101/2023.02.13.528370

**Authors:** Seth D. Fortmann, Blake F. Frey, Vidya Sagar Hanumanthu, Shanrun Liu, Andrew Goldsborough, Kameron V. Kilchrist, P. Brent Ferrell, Casey T. Weaver, Maria B. Grant, Robert S. Welner

## Abstract

Single-cell RNA-sequencing (scRNA-seq) presents an opportunity to deconstruct cellular networks but is limited by the loss of biological information, including *in vivo* cellular states and phospho-signaling. Herein, we present fixation before dissociation using a deep eutectic solvent (DES), which preserves multiple domains of *in vivo* biological data, including morphology, RNA, proteins, and post-translational modifications. In scRNA-seq of viable versus DES bone marrow, dissociation induced global stress responses, immune and stromal cell activation, and loss of highly sensitive cell populations, which were prevented with DES. Further, we introduce a validated and flexible method for performing intracellular CITE-seq in DES-fixed cells. Leveraging this approach during Th17 T cell stimulation allowed the simultaneous quantification of transcriptomes and four phosphorylated proteins, leading to the identification of a hyperactivated state in p-ERK/p-FOS double positive cells, which we experimentally validated. We anticipate that DES-based fixatives will allow the accurate reconstruction of *in vivo* cellular networks and uncover cooperativity amongst intracellular pathways.

## INTRODUCTION

Biological systems are composed of cellular networks that rely on the continuous, massive exchange of information to maintain homeostasis. As such, achieving a complete understanding of tissues in health and disease will require deconstructing these cellular networks, and this will necessitate the simultaneous collection of multiple domains of biological data at high resolution and in a massively parallel fashion. Of equal necessity is that the captured data accurately reflects that being exchanged *in vivo*. This remains a fundamental obstacle in studying procured tissues because information flow and adaptation continue until cellular termination, thereby perpetually deviating further from the *in vivo* state. To this end, single-cell RNA-sequencing (scRNA-seq) presents an opportunity to achieve the resolution and scale required for deconstructing cellular networks, but the loss of biological data, including that of the *in vivo* state as well as entire information domains like intracellular signaling, remains a core challenge.

Fundamental to this loss of *in vivo* information in scRNA-seq is the need to dissociate solid tissues into a single-cell suspension. At present, this is performed on viable specimens through enzymatic and/or mechanical approaches. While the harshness and duration are tissue-specific, dissociation, by definition, involves the complete destruction of the extracellular matrix and all cell-to-cell connections. Such a drastic perturbation would be expected to alter cellular states, but the effects of dissociation on single-cell technologies are challenging to know. Nonetheless, one approach is to compare single-cell (scRNA-seq) to single-nucleus RNA-sequencing (snRNA-seq) from the same tissue type, where snRNA-seq eliminates the need for viable dissociation but at the cost of retaining only nuclear information. While such comparisons have important limitations, the results frequently reveal upregulation of stress and activation signatures and partial-to-entire losses of sensitive cell types. In the case of disease states, these shifts may be exacerbated by preexisting pathology, making it challenging to differentiate artifacts from true disease processes (Wu et al., 2019). For example, in brain specimens from patients with epilepsy, snRNA-seq failed to capture the microglial activation signature that was observed in dissociated versions of the same samples with scRNA-seq (Thrupp et al., 2020). In the retina, single-nuclei recovered entire neural populations and supporting cell types that were effectively absent with tissue dissociation (Santiago et al., 2022). Epithelial cells are especially sensitive to dissociation and are often underrepresented in single-cell data. In the kidney, dissociation induced prominent stress artifacts in tubular epithelial cells, which were absent with snRNA-seq (Wu et al., 2019), and in the colon and liver, dissociation altered epithelial gene expression and skewed cell recovery (Oh et al., 2022). Beyond transcription, remarkably little is known about how tissue dissociation influences the activity of intracellular pathways, including phospho-signaling, which are key determinants of cell phenotype.

Fixation of tissues prior to dissociation is an attractive alternative to preserve the *in vivo* state. The most widely used fixatives are formaldehydes and alcohols, which act through protein cross-linking and dehydration, respectively. Fixation prior to dissociation has been achieved using diluted paraformaldehyde, which was sufficient to prevent activation artifacts in muscle stem cells (Machado et al., 2017). However, formalin causes cross-linking of RNAs, which necessitates a harsh cross-link reversal step that is widely known to skew RNA-sequencing metrics (Jones et al., 2019). Conversely, alcohol-based fixation is non-crosslinking and generally results in higher-quality RNA, but sequencing is biased by inherent transcript characteristics, such as length and G-C content (Wang et al., 2021). For these reasons, novel biological fixatives are needed that allow dissociation post-fixation without biomolecule degradation or transcript skewing.

Deep eutectic solvents (DES) are a novel class of compounds with unique chemical properties that are favorable for next-generation fixatives. DES’s are composed of two room-temperature solids, one hydrogen bond donor, such as urea, and one hydrogen bond acceptor, such as choline chloride, which when mixed in particular stoichiometric ratios, cause a characteristically large melting point depression and the subsequent formation of a viscous room-temperature liquid (Abbott et al., 2003; Hansen et al., 2021). A subset of DES’s, including the commercially available product vivoPHIX™ (RNAssist, Cambridge UK), rapidly penetrate and fix bacterial, plant, and animal cells, likely through prolific intermolecular hydrogen bonding that displaces water molecules in the hydration shells of biomolecules (Goldsborough and Bates, 2019). Importantly, DES fixation reportedly allows the long-term stabilization of biomolecules, while weakening cell-to-cell interactions thereby facilitating dissociation post-fixation (Goldsborough and Bates, 2021). Thus, DES-based compounds are an attractive alternative to traditional biological fixatives and have the potential to preserve *in vivo* biology.

Herein, we demonstrate that fixation before dissociation using a DES (vivoPHIX™) preserves *in vivo* transcription and phosphorylation in single-cell technologies. We show that DES retains complex cellular morphologies with ultrasonic dissociation, such as neurons of the human retina. We demonstrate that DES allows the long-term stabilization of biobanked tissues and results in high-quality scRNA-seq data that lack activation artifacts observed in paired viable samples. Lastly, we show that DES preserves post-translational modifications and provides sufficient membrane permeabilization for probing intracellular epitopes. We demonstrate a validated and flexible approach to intracellular CITE-seq (cellular indexing of transcriptomes and epitopes by sequencing) using DES and apply it to study four phospho-targets during T cell activation, revealing specific pathways associated with cytokine production and hyperactivated p-FOS/p-ERK double positive cells. We anticipate that the use of DES-based fixatives in single-cell technologies will improve the reconstruction of *in vivo* cellular networks and allow the parallel integration of transcriptomes and intracellular signaling pathways in single cells.

## RESULTS

### DES-Based Fixation Preserves RNA Quality and Quantity

We developed a novel approach, using a DES-based fixative (vivoPHIX™), to preserve *in vivo* cellular states with minimum loss of biological information in dissociated tissues (**Fig. 1A**). In overview, cells or whole tissues are immersion fixed in DES for ≥2 hours at room temperature, followed either by immediate downstream processing or long-term storage at -80°C. For tissues, dissociation post-fixation is achieved using an ultrasonication water bath. The cell suspension is then briefly treated with 10% glacial acetic acid (AcOH; diluted in DES), which inactivates cellular RNAses upon resuspension in aqueous buffer (Lopez-Alonso et al., 2010). The 10% AcOH/DES cell slurry is then transferred to an aqueous solution via an intermediate wash step with excess 3x saturated sodium chloride (SSC), followed by resuspension in 1x SSC for all downstream processing. The cell suspension can then be used for a wide variety of standard analytical techniques.

**Figure 1.**
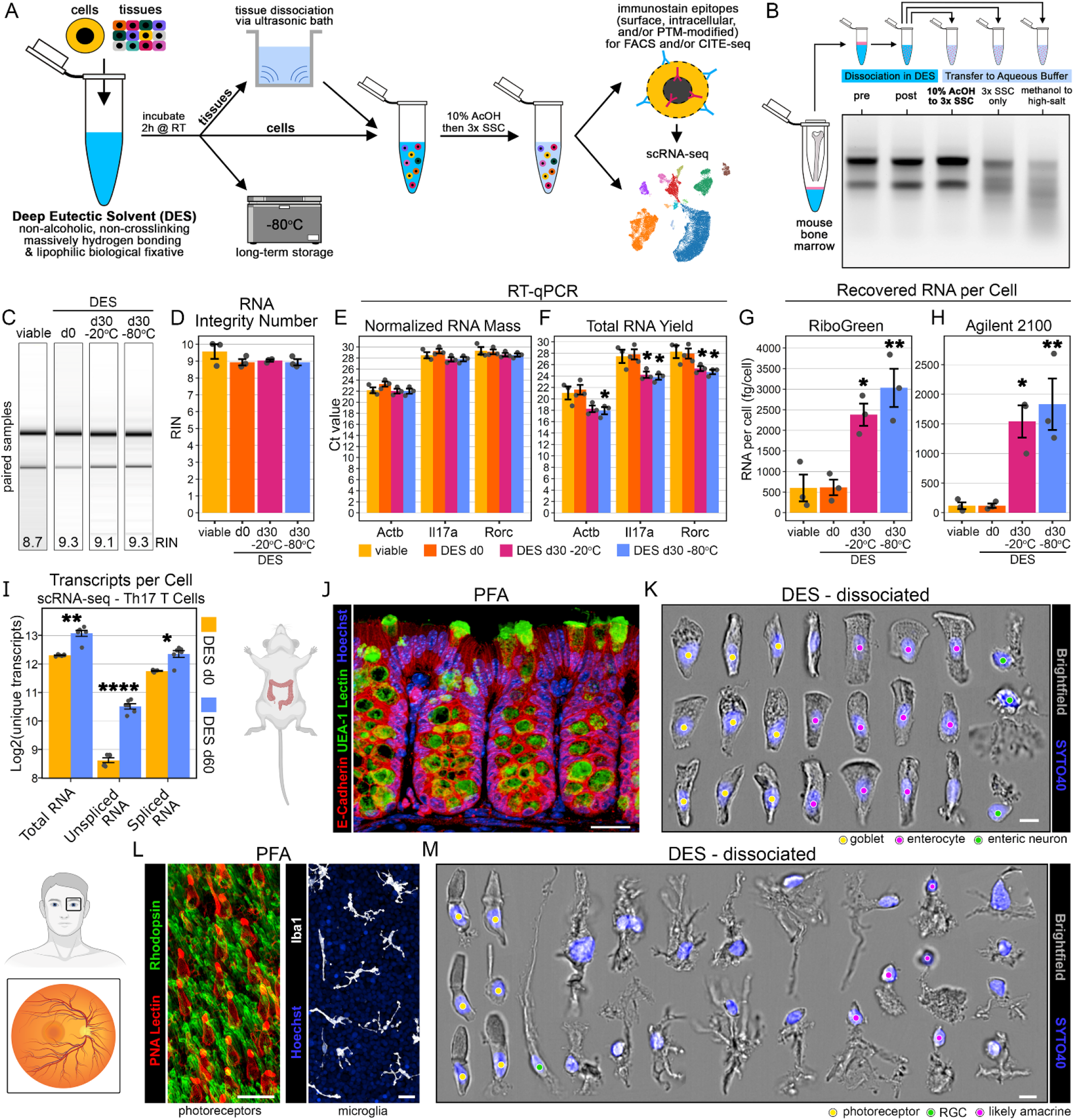
DES fixation allows long-term preservation of RNA and *in vivo* cell morphology after ultrasonic dissociation. (**A**) Schematic overview of fixation before dissociation using DES. h, hours. RT, room temperature. SSC, saturated sodium chloride. AcOH, acetic acid. PTM, post-translational modifications. FACS, fluorescence-activated cell sorting. (**B**) Gel electrophoresis for RNA integrity from mouse bone marrow before and after ultrasonic dissociation and transfer to aqueous buffer using three different methods: (1) 10% AcOH in DES for 5 minutes at RT followed by excess 3x SSC, (2) excess 3x SSC alone, or (3) methanol followed by high salt buffer [4M ammonium sulfate]. The same bone marrow sample was used for each step. (**C**) Automated gel electrophoresis for RNA integrity from primary mouse T cells. The same culture is shown across each timepoint/condition. (**D**) RNA integrity number (RIN) from primary mouse T cells. (**E-F**) RT-qPCR from 100,000 FACS-sorted primary mouse T cells using two forms of normalization: (**H**) constant RNA mass per sample and (**G**) total RNA yield per sample. (**G-H**) RNA recovery per cell from 100,000 FACS-sorted primary mouse T cells using two methods: (**G**) RiboGreen fluorescence assay and (**H**) Agilent 2100. (**I**) Total transcripts per cell from scRNA-seq on paired Th17 T cell cultures from either the day of fixation (d0) or samples stored for 60 days in DES at -80°C (d60). Mean measurements per culture/condition combination (n=6; condition=stimulated or unstimulated). (**J**) Confocal microscopy of PFA fixed mouse colon. Scale bar=25μm. (**K**) Collage of imaging flow cytometry photos from DES fixed and ultrasonically dissociated mouse colon. Scale bar=7μm. Yellow circles=goblet cells, magenta circles=enterocytes, green circles=enteric neurons. (**L**) Confocal microscopy of PFA fixed human retina whole mount showing photoreceptors (left) and microglia (right). Scale bars=25μm. (**M**) Collage of imaging flow cytometry photos from DES fixed and ultrasonically dissociated human retina. Scale bar=7μm. Yellow circles=photoreceptors, green circle=retinal ganglion cells (RGC), magenta circles=likely amacrine cells. (**L**) and (**M**) are from the same non-diseased donor. (**E**,**F**,**I**) Two-way ANOVA with Bonferroni correction (control group=viable); * p < 0.05; ** p ≤ 0.01; **** p ≤ 0.0001. (**D**,**G**,**H**) One-way ANOVA with Bonferroni correction (control group=viable); * p ≤ 0.05; ** p ≤ 0.01. (**D-H**) n=3 independent cultures.

First, we optimized the preservation of mouse bone marrow in DES, which involved centrifuging whole bone marrow directly into the fixative (**Fig. 1B**). Using gel electrophoresis to visualize RNA degradation, we compared pre- and post-dissociation of identical bone marrow samples and found that ultrasonication had no qualitative effect on RNA integrity (**Fig. 1B**). We then optimized the transition from DES to aqueous solution in fixed, dissociated bone marrow, which is required for compatibility with standard downstream analytical techniques. Similar to other fixatives like methanol (Chen et al., 2018), the transition from DES to aqueous buffer is a critical and highly sensitive step where RNAses can become reactivated, leading to acute degradation. We tested 3 aqueous transition strategies in paired bone marrow samples: (1) treatment with 10% AcOH followed by excess 3x SSC, (2) excess 3x SSC alone, as previously reported for methanol-fixed cells (Chen et al., 2018), and (3) methanol treatment followed by resuspension in a high-salt buffer akin to diluted RNAlater, as recently reported for methanol-fixed cells (Amit et al., 2020; Katzenelenbogen et al., 2020). With the 10% AcOH to 3x SSC approach, gel electrophoresis revealed intact and undegraded RNA, whereas 3x SSC alone resulted in moderate RNA degradation and methanol to high-salt resulted in severe RNA degradation (**Fig.1B**).

Next, using the AcOH transition strategy, we quantitatively assessed the ability of DES to preserve RNA quality and yield in acutely fixed and long-term stored samples. To this end, we utilized a murine primary culture approach of Th17 T cells (Harrington et al., 2005), which allowed for a homogenous and tightly controlled primary cell population. The cultures were split into viable and DES fixed groups, and aliquots of the DES fixed cells were stored at -20°C or -80°C for 1 month to determine the effects of long-term storage. It should be noted that, even at -80°C, DES does not freeze. Gel electrophoresis for visualization of RNA degradation (**Fig. 1C**) and RNA Integrity Number (RIN) analysis (**Fig. 1D**) confirmed the ability of DES to preserve RNA quality. We performed RT-qPCR using 100K fluorescence-activated cell sorting (FACS)-sorted cells per sample and found no difference in Ct values between groups when normalized by input RNA mass per sample (**Fig. 1E**), consistent with preserved RNA quality. Unexpectedly, normalization by total RNA yield per sample returned significantly decreased Ct values in long-term stored samples (**Fig. 1F**), suggesting increased RNA recovery per cell. We measured the RNA quantity per cell using two independent methods and determined that DES d0 samples had essentially identical recovery to that of paired viable samples (**Fig. 1G-H**). However, we again found that samples stored for 30d in DES at -20°C or -80°C resulted in significantly increased RNA yields (**Fig. 1G-H**). We reasoned that this increase in RNA recovery may be due to continued fixation that facilitates transcript release from RNA binding proteins and/or promotes more complete cell lysis. scRNA-seq data of DES fixed Th17 cells from paired d0 versus d60 samples were downsampled to an even number of unique reads per cell and analyzed for total transcripts, which confirmed increased quantities in long-term stored samples (**Fig. 1I**; **Supp. Fig. 1)**. Interestingly, when using splicing to distinguish transcript recovery, we found that unspliced RNA was disproportionately increased in DES d60 samples compared to spliced RNA (fold-change d60/d0 [mean±SD]; unspliced: 3.74±0.56; spliced: 1.54±0.27; **Fig. 1I)**. Because the majority of RNA binding proteins and unspliced RNA are located within the nucleus (Van Nostrand et al., 2020), this data supports that longer durations of DES exposure either facilitate the disruption of RNA-protein interactions and/or promote more complete nuclear lysis, leading to increased recovery of nascent RNA.

### Ultrasonic Dissociation of DES-Fixed Tissues Preserves *in vivo* Cell Morphology

Current methods for tissue dissociation result in changes to cellular structure and loss of morphological information; this is especially evident in sensitive cell types with complex morphologies, like neurons and epithelial cells (Fadl et al., 2020). Therefore, methods aimed at preserving *in vivo* biology in dissociated tissues should also retain *in vivo* morphology. To this end, we tested the ability of DES to preserve morphological information in dissociated cells by comparing confocal microscopy of PFA-fixed tissues to imaging flow cytometry of their DES-dissociated counterparts. We chose two divergent tissue types, both of which are known to pose challenges in scRNA-seq (Oh et al., 2022; Santiago et al., 2022), mouse colon epithelium (**Fig. 1J-K**) and human retina (**Fig. 1L-M**).

We first explored the ability of DES to preserve epithelial morphology, a broad class of directionally polarized cells. To this end, we used mouse colon and compared PFA-fixed sections to DES-dissociated large intestine. In PFA-fixed sections, E-cadherin staining distinguished the colonic epithelium, which lined the crypts and the interface of the intestinal lumen, and UEA-1 lectin identified mucin-producing goblet cells, with characteristic narrow bases and cup-shaped apical bodies (**Fig. 1J**). Imaging flow cytometry of DES-dissociated colon revealed near identical morphologies, and subtypes of colonic epithelium were readily identifiable (**Fig. 1K**). We observed many columnar cells, consistent with enterocytes, which varied from rectangular to funnel-shaped and often had brush boarder-like structures at the apical surface (**Fig. 1K**, magenta circles). Goblet cells were also easily identifiable with narrow bases, cup-like apical structures, and high granularity (**Fig. 1K**, yellow circles). Remarkably, we also observed whole enteric neurons with dendritic ramifications of various sizes and orientations (**Fig. 1K**, green circles).

Given that DES preserved the morphology of enteric neurons, we next assessed human retina, which as an extension of the central nervous system, is composed of a variety of neural cell types with unique, complex morphologies. Compared to confocal microscopy of PFA-fixed retinal wholemounts (**Fig. 1L**), imaging flow cytometry of DES-dissociated retina from the same donor revealed remarkably preserved cell morphologies (**Fig. 1M**). Photoreceptors (**Fig. 1L**, left), in particular cones, retained intact somata and inner segments, and in some cases, even synaptic termini and outer segments were preserved (**Fig. 1M**, yellow circles). Cells with long axonal projections, akin to retinal ganglion cells, were observed (**Fig. 1M**, green circle), as well as neurons that resembled amacrine cells with somata connected to bistratified or monostratified neurites terminating in large dendritic arbors (**Fig. 1M**, magenta circles). In addition, we recovered a variety of other neurons of unknown type that retained many intact dendritic processes with various arrays of branching patterns and ramifications (**Fig. 1M**). Together, these data spanning divergent tissues across mice and humans demonstrate the ability of DES to preserve the *in vivo* morphology of a variety of delicate cell types.

### scRNA-seq of DES Tissue Recovers Similar Cell Populations and Cluster Frequencies Without Transcript Bias

Next, we assessed the ability of DES to preserve *in vivo* transcription by performing a head-to-head scRNA-seq comparison of paired viable versus DES bone marrow. Bone marrow represents a favorable test case because it is a heterogeneous tissue with several cell types that pose distinct challenges in scRNA-seq. In viable bone marrow, the majority of cells were CD45+ immune cells, followed closely by Ter119+ erythrocytes, while CD45-Ter119-stromal-enriched cells accounted for the smallest population (**Fig. 2A**). In DES bone marrow, similar proportions of the same populations were observed, except more stromal-enriched cells were recovered, and Ter119 needed to be replaced with Spectrinβ, another erythrocyte-specific marker (**Supp. Fig. 2**), due to loss of antigenicity from DES fixation (**Fig. 2B**). In viable bone marrow, live cells were gated using viability dye (**Fig. 2A**; **Supp. Fig. 2**), while in DES-fixed bone marrow, intact DNA+ cells were gated using SYTO40, a DNA dye with a narrow excitation/emission in the violet spectrum (**Fig. 2B**; **Supp. Fig. 2**).

**Figure 2.**
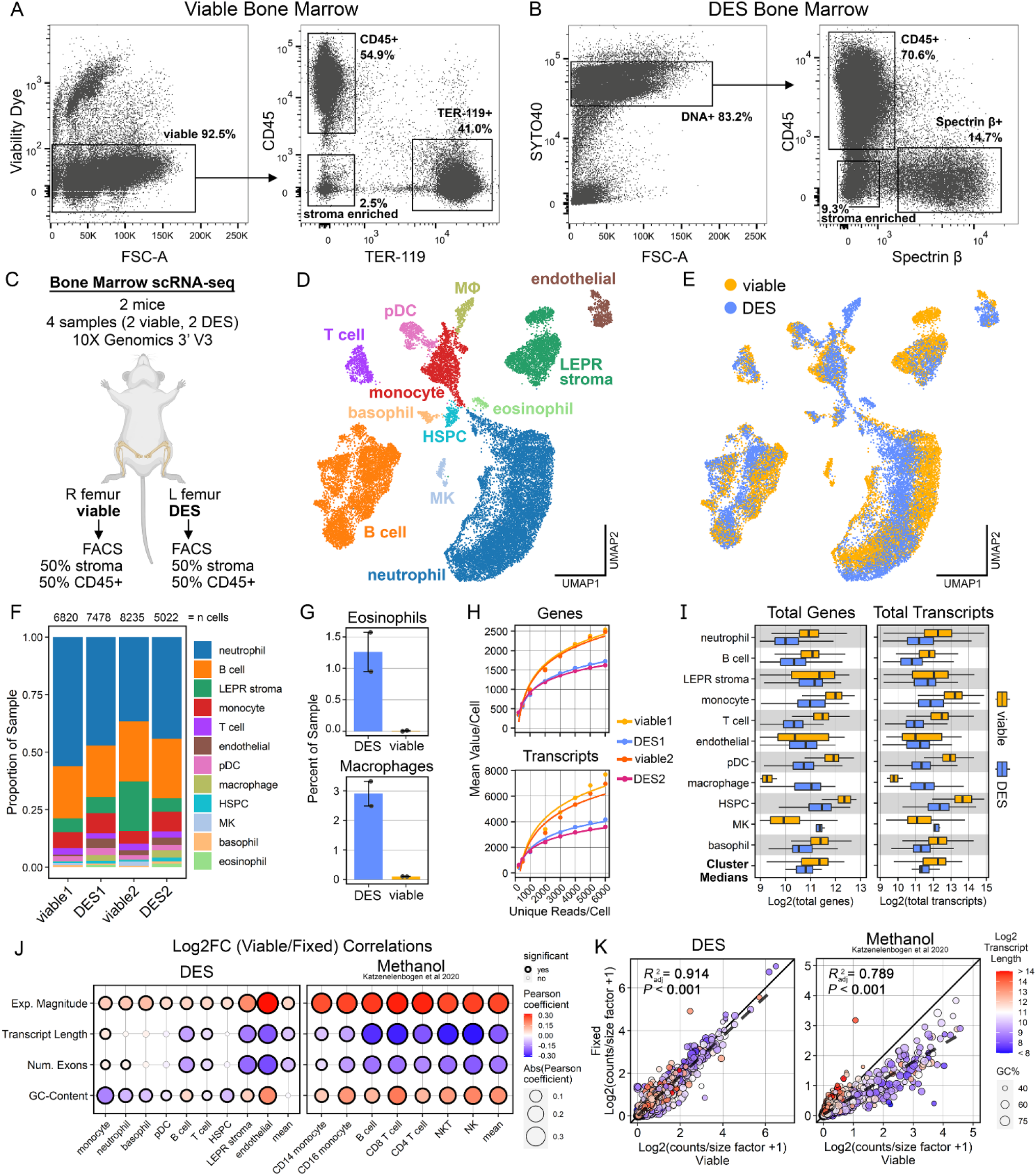
scRNA-seq of DES-fixed mouse bone marrow recovers similar cell populations and cluster frequencies without transcript bias. (**A-B**) Flow cytometry gating from (**A**) viable and (**B**) DES-fixed mouse bone marrow. (**C**) Schematic overview of scRNA-seq design. (**D-E**) Uniform manifold approximation and projection (UMAP) of scRNA-seq data from 27,555 mouse bone marrow cells, annotated by (**D**) clusters or (**E**) sample type. MK, megakaryocyte. HSPC, hematopoietic stem and progenitor cell. pDC, plasmacytoid dendritic cell. MΦ, macrophages. (**F**) Cluster proportions of each sample. (**G**) Percent of total sample for eosinophils (top) and macrophages (bottom) in DES versus viable. (**H**) Stepwise downsampling of scRNA-seq showing fitted curves per individual replicate for total genes (top) and total transcripts (bottom). (**I**) Per cluster quantifications of total genes (left) and total transcripts (right) per cell. Cluster medians refers to plotting the medians of each cluster for DES versus viable. (**J**) Per cluster Pearson correlation analyses comparing gene-wise Log2 Fold Changes (Log2FC) of viable versus fixed (viable/fixed) expression to G-C Content, Number of exons, transcript length, and expression magnitude (with respect to viable samples). Left plot is for DES fixation in mouse bone marrow and right plot is for methanol fixation in human PBMCs from Katzenelenbogen *et al* 2020. Mean refers to the overall average of all cluster means with respect to gene-wise expression. Bonferroni was used for p-value correction. (**K**) Gene-wise linear regressions of size factor normalized expression for the overall sample mean comparing viable versus fixed. Left plot is for DES fixation in mouse bone marrow and right plot is for methanol fixation in human PBMCs from Katzenelenbogen *et al* 2020. Overall sample mean refers to the average of all cluster means with respect to gene-wise expression, as done in (**J**).

For our bone marrow scRNA-seq, we used four samples from two mice. From each animal, one femur was used for viable bone marrow and the other for DES after 2 hours of fixation; half of each sample was FACS sorted for immune cells (CD45+ Ter119-or Spectrinβ-), while the other half was sorted for stromal-enriched cells (CD45-Ter119-or Spectrinβ-; **Fig. 2C**). To control for differences in sequencing depth, we downsampled each of the 4 biological replicates to a common ratio of unique reads per cell. After quality control filtering, removal of contaminating erythrocytes, and dimensionality reduction, 12 main clusters were identified from 27,555 total cells, with the largest cluster being neutrophils, followed by B cells, LEPR+ stroma, monocytes, and T cells (**Fig. 2D**; **Supp. Fig. 3**). The proportions of these predominant populations were similar between viable and DES samples (**Fig. 2E-F**). However, rarer populations showed greater heterogeneity; for example, macrophages and eosinophils, which are known to be poorly represented in bone marrow scRNA-seq (Diny et al., 2022; Millard et al., 2021), were nearly exclusive to DES samples (**Fig. 2G**). DES samples also had higher recovery of endothelial cells and hematopoietic stem and progenitor cells (HSPCs), whereas megakaryocytes (MKs) were more frequent in viable bone marrow (**Fig. 2E-F**).

**Figure 3.**
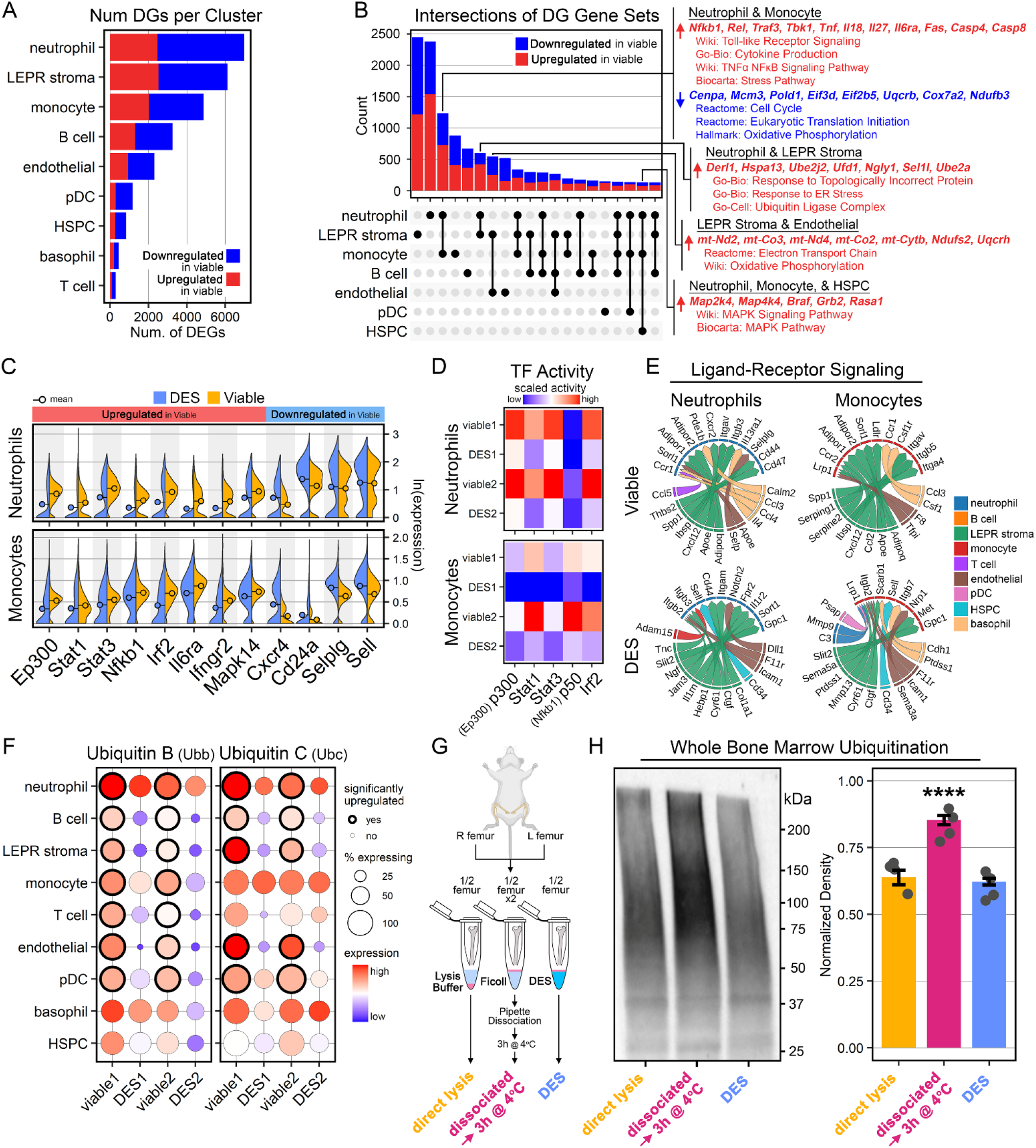
DES prevents immune cell activation and global stress response induced by tissue dissociation. (**A**) Total number of DGs per cluster for viable versus DES. Eosinophil, MK, and macrophage not tested because <100 cells per group. (**B**) Upset plot showing intersections of cluster-wise DGs. Selected DGs and enriched pathways for four gene sets are shown (right). Red text=upregulated. Blue text=downregulated. (**C**) Expression of selected upregulated and downregulated DGS in neutrophils (top) and monocytes (bottom) from the neutrophil—monocyte gene set in (**B**). Each split violin represents a paired biological sample from the same mouse. (**D**) SCENIC transcription factor (TF) activity for neutrophils (top) and monocytes (bottom) showing the 5 TFs from (**C**). Viable1 and DES1 are paired samples from one mouse, as are viable2 and DES2. (**E**) Circosplots of top 15 ligand-receptor pairs from NicheNet analysis for neutrophils (left) and monocytes (right), comparing viable (top row) versus DES (bottom row). Ligands are represented on bottom half of circosplots and receptors (from either neutrophils or monocytes) are on the top half. (**F**) Mean expression of ubiquitin genes, Ubb (left) and Ubc (right), per cluster for each individual biological replicate. Statistical significance, determined by cluster-wise differential gene expression analyses, is represented with thickened black outline. (**G**) Schematic of experimental design for (**H**). (**H**) Immunoblot for ubiquitin (left) and quantification per group (right) from paired mice (n=4). Normalized to total protein stain. One-way ANOVA with Dunnett’s post-hoc test (control group=Direct Lysis); **** p ≤ 0.0001.

To determine RNA recovery in viable versus DES scRNA-seq, we compared the total number of genes and transcripts per cell using two methods: stepwise downsampling of each biological replicate and total recovery on a per cluster basis. Downsampling analysis showed that, globally, DES samples tended to recover fewer genes and transcripts per cell (**Fig. 2H**). Similarly, on a per cluster basis, most hematopoietic cells tended to recover more genes and transcripts in viable samples, with the exception of macrophages and MKs, which had greater recover with DES fixation (**Fig. 2I**). For non-hematopoietic cells, LEPR+ stroma had similar RNA counts whereas endothelial cells had greater recovery of genes and transcripts with DES fixation (**Fig. 2I**). Comparing the average of cluster medians, genes per cell in DES versus viable (mean ± SD) were 1,858±574 versus 2,584±1,407 respectively, and transcripts/cell were 2,996±1,057 versus 5,318±3,547 respectively (**Fig. 2I**).

Methanol, the current gold-standard fixative in scRNA-seq, is known to artificially skew RNA recovery based on inherent transcript characteristics, like nucleotide length and G-C content (Wang et al., 2021). To determine whether differences in gene expression between DES and viable samples were related to transcript bias, we calculated gene-wise Log2 fold-changes (Log2FC; viable/DES) for each cluster and performed Pearson correlation analyses with G-C content, number of exons, transcript length, and the expression magnitude of each gene (**Fig. 2J**). For comparison to methanol fixation, we performed an identical analysis using similar data from a recent scRNA-seq report (Katzenelenbogen *et al*., 2020) that applied methanol fixation to human peripheral blood mononuclear cells (PBMCs) versus paired viable samples (**Supp. Fig. 4**). In methanol fixation, all clusters as well as the overall cluster mean showed strong, significant correlations with each of the four metrics, with G-C content and expression magnitude showing positive associations and transcript length and the number of exons displaying inverse associations (**Fig. 2J**). Conversely, with DES fixation, only expression magnitude had a consistent relationship across all clusters and the cluster mean, with weaker, albeit still significant, positive correlations to Log2FC (**Fig. 2J**). G-C content, transcript length, and number of exons in DES fixation showed mostly weak and inconsistent associations, with some clusters having non-significant correlations and others showing opposite directions of association (**Fig. 2J**). Lastly, for each cluster as well as the overall average, we performed linear regressions by comparing gene expression in fixed versus viable samples from DES and methanol fixation, respectively. Amongst clusters, gene expression in DES samples showed overall greater correlations compared to methanol, with 67% of clusters having adjusted R^2^ > 0.9 versus 0% with methanol (**Supp. Fig. 5**). Similarly, for the overall average, gene expression in fixed versus viable samples had a stronger correlation with DES compared to methanol (adjusted R^2^ 0.914 versus 0.789, respectively; **Fig. 2K**). As expected, genes with higher expression in methanol samples tended to have longer transcripts and lower G-C contents, whereas in DES samples, these same gene characteristics appeared stochastically distributed (**Fig. 2K**). Thus, unlike other fixation methods, DES lacks transcript bias.

**Figure 4.**
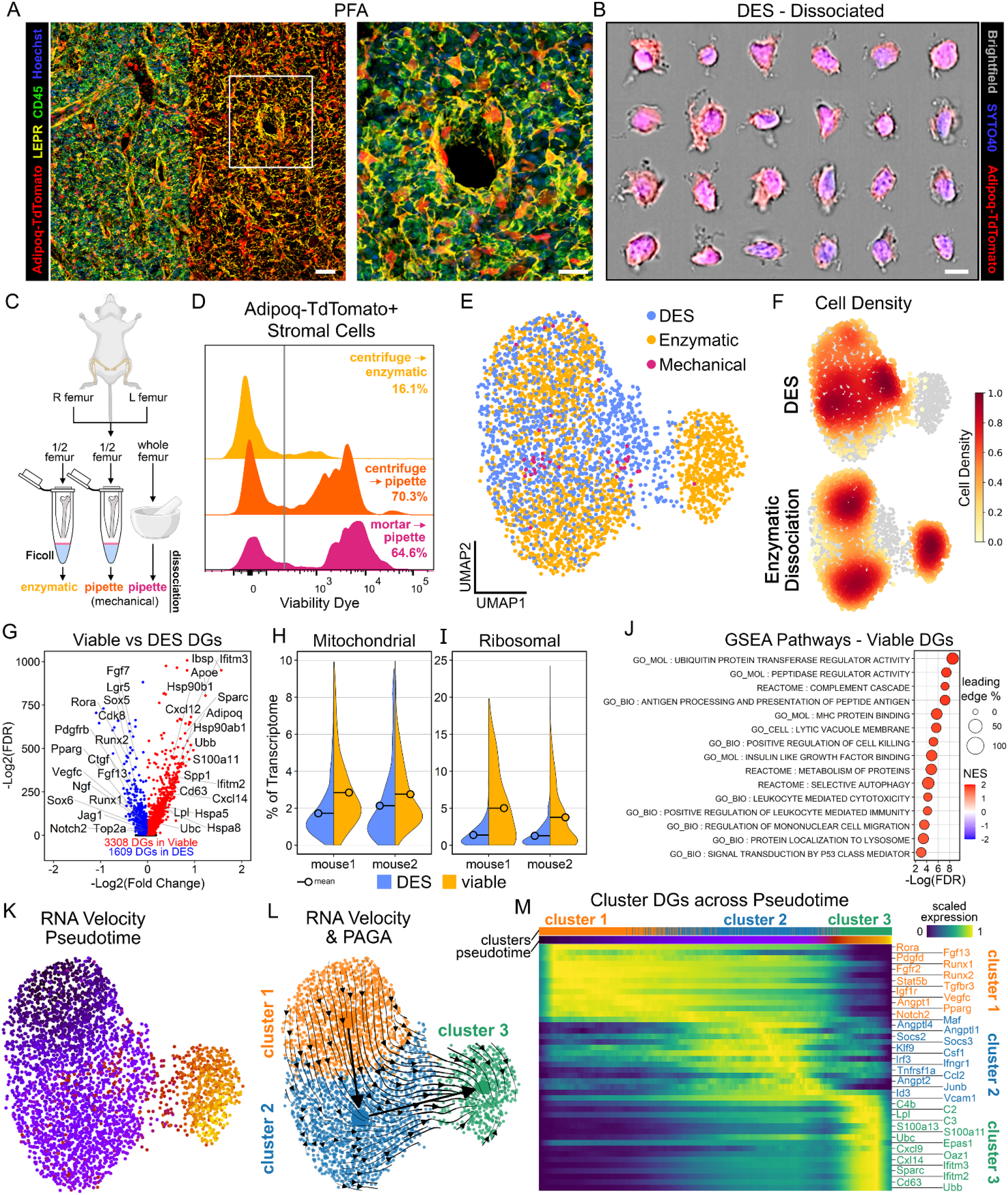
DES fixation protects bone marrow stroma from dissociation-induced injury and artificial activation. **A**) Confocal microscopy of a PFA-fixed femoral section for Adipoq-TdTomato transgenic mice. Left scale bar=40μm. Right scale bar=20μm. (**B**) Collage of imaging flow cytometry photos of TdTomato+ stroma from DES fixed and ultrasonically dissociated mouse bone marrow. Scale bar=7μm. (**C**) Schematic of experimental design for (**D**). (**D**) Flow cytometry histogram plots of viability dye from TdTomato+ bone marrow stroma using 3 different dissociation methods on one paired sample. Gray line indicates viability gate. (**E**) UMAP of combined scRNA-seq data from earlier optimization dataset (DES versus mechanical dissociation; n=2 samples from n=1 mouse) and **Fig. 2C** dataset (DES versus enzymatic dissociation; n=4 samples from n=2 mice). 3,876 total LEPR stroma cells annotated according to DES or viable dissociation method. (**F**) LEPR stroma density with respect to UMAP space. Top row is DES, bottom row is enzymatic dissociation. (**G**) Volcano plot of differential gene expression analysis comparing viable stroma versus DES. (**H**,**I**) Violin plots showing the percentage of the total transcriptome derived from either (**H**) mitochondrial or (**I**) ribosomal reads for viable versus DES stroma. (**J**) Selected top GSEA pathways from viable versus DES stroma. Mitochondrial and ribosomal pathways are not shown. (**K**) UMAP of bone marrow stroma scRNA-seq showing RNA velocity pseudotime. (**L**) UMAP of bone marrow stroma scRNA-seq showing RNA velocity and partition-based graph abstraction (PAGA). Annotation is of stroma sub-clustering showing 3 discrete clusters. (**M**) Smoothened heatmap showing pseudotime activation trajectory of stroma. Cluster specific DGs were calculated using viable stroma.

**Figure 5.**
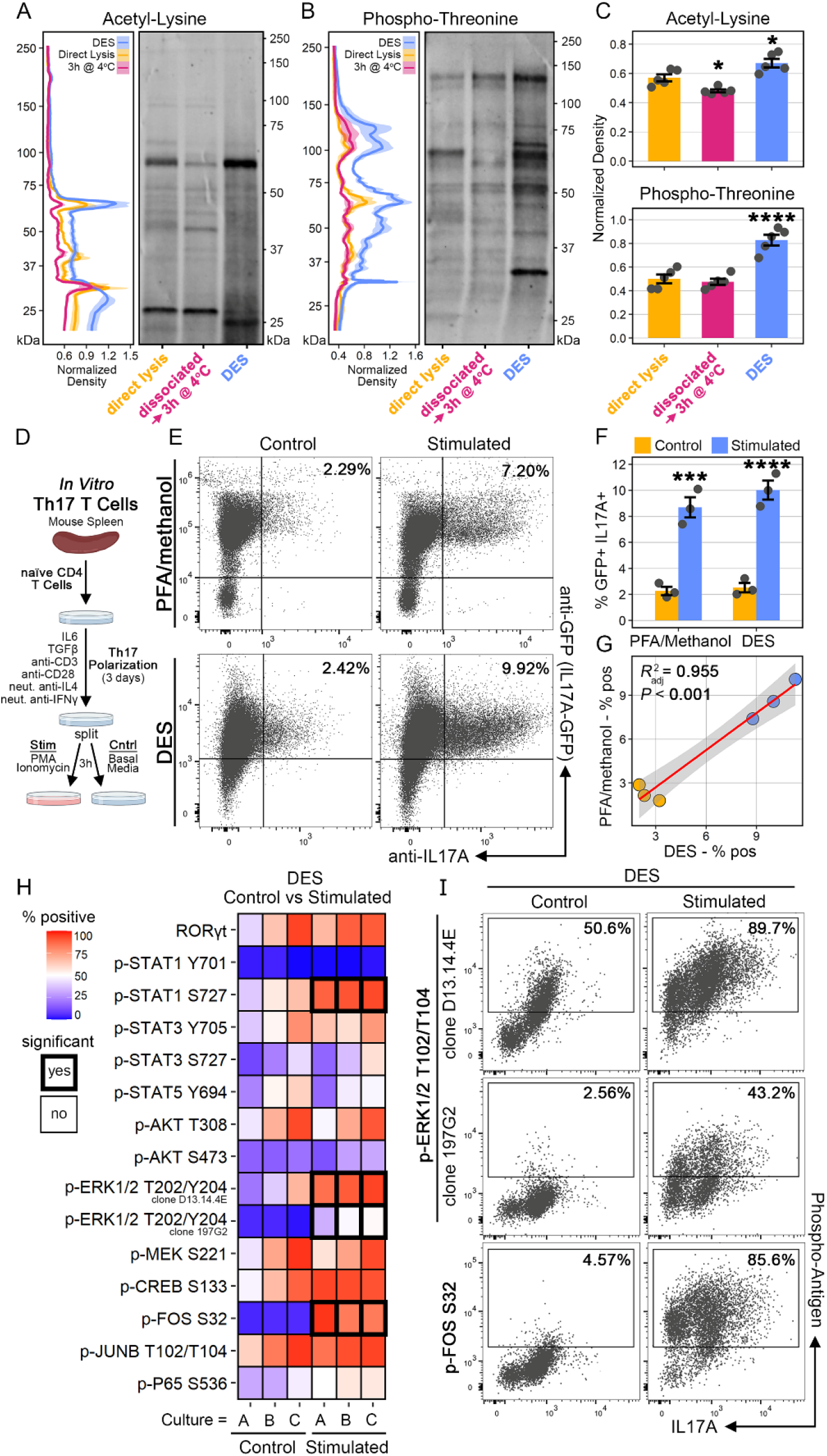
DES preserves post-translational modifications and provides sufficient permeabilization for intracellular immunostaining. (**A-B**) Immunoblots of bone marrow from paired mice for (**A**) acetyl-lysine and (**B**) phospho-threonine using identical experimental setup to **Fig. 3G**. Error bars represent SEM from n=5 mice. (**C**) Immunoblot quantifications of (top) acetyl-lysine and (bottom) phospho-threonine (n=5). One-way ANOVA with Dunn’s post-hoc test (control group=direct lysis). * p < 0.05; **** p ≤ 0.0001. (**D**) Schematic of experimental setup for (**E-G**). (**E**) Representative flow cytometry plots of stimulated and unstimulated (control) primary Th17 T cells from Il17a-GFP transgenic mice. Top row is standard PFA/methanol. Bottom row is DES. The same number of events are shown in all 4 plots. (**F**) Quantification of GFP+/IL17a-protein+ cells with PFA/methanol versus DES (n=3). One-way ANOVA with Dunn’s post-hoc test. *** p ≤ 0.001; **** p ≤ 0.0001. (**G**) Linear regression comparing GFP+/IL17a-protein+ cells with PFA/methanol versus DES. (**H**) Heatmap representation of flow cytometry data using stimulated versus control Th17 T cells showing percent positivity for 14 different targets. Identical experimental design to (**D**) except non-transgenic mice were used and no PFA/methanol groups were included. Isotype controls were used for gating and all cells were fixed with DES. Two-way ANOVA with Holm-Sidak post-hoc test. (**I**) Representative flow cytometry plots from (**H**) showing 2 different antibody clones to phospho-ERK1/2 T204/Y204 (top and middle) and phospho-FOS S32 (bottom). The same number of events are shown in all 6 plots.

### DES Prevents Artificial Immune Cell Activation and Global Stress Responses Caused by Tissue Dissociation

To determine whether DES fixation preserves *in vivo* states, we analyzed changes in gene expression patterns across fixed versus viable samples in our bone marrow scRNA-seq dataset. Across all clusters, neutrophils, LEPR+ stroma, monocytes, B cells, and endothelium had the greatest number of differentially expressed genes (DGs), with a relatively even distribution with respect to the direction of change (**Fig. 3A**). To determine if particular DG gene sets were unique to one cell population or shared with others, we analyzed the intersections of upregulated and downregulated DGs across clusters to identify shared patterns. The top 2 gene sets that accounted for the highest number of DGs were unique to LEPR+ stroma (2,450 DGs) and neutrophils (2,378 DGs), respectively (**Fig. 3B**). The third highest gene set (1,236 DGs) was shared between neutrophils and monocytes (**Fig. 3B**) and pathway enrichment analysis revealed upregulation of stress pathways (Fas, Casp4, Casp8) and inflammatory signaling including cytokine production (Tnf, Il18, Il27) and NF-Kb activation (Nfkb1, Rel, Traf3, Tbk1). Downregulated pathways in the neutrophil-monocyte gene set included cell cycle (Cenpa, Mcm3, Pold1) and oxidative phosphorylation (Uqcrb, Cox7a2, Ndufb3). The next largest shared gene set (602 DGs) was between neutrophils and LEPR+ stroma and was enriched with an upregulation of genes related to endoplasmic reticulum stress and ubiquitination (**Fig. 3B**). Of additional note was a shared upregulation of mitochondrial genes in LEPR+ stroma and endothelium and an upregulation of mitogen-activated protein kinases (MAPKs) signaling in neutrophils, monocytes, and HSPCs (**Fig. 3B**).

Given the predominant gene expression shifts in neutrophils and monocytes, we explored these two populations further. Several key genes involved in immune cell activation were upregulated with respect to viable samples in neutrophils and monocytes (**Fig. 3C**). These included pioneering factors like p300 (Ep300), pro-inflammatory transcription factors (Stat1, Stat3, Nfkb1, Irf2), activation pathways like p38 MAPK (Mapk14), and pro-inflammatory cytokine receptors like Il6ra and Ifngr2 (**Fig. 3C**). Conversely, neutrophils and monocytes shared a downregulation, relative to DES, of genes coding for adhesion proteins/receptors involved in bone marrow retention, including Cxcr4, Cd24a, Selectin P ligand (Selplg), and Selectin L (Sell; **Fig. 3C**). Using SCENIC (Aibar et al., 2017) to infer transcription factor (TF) activity by assessing enrichment of TF motifs in the regulatory regions of expressed genes, we again found increases in TF activity for p300, STAT1, STAT3, p50, and IRF2 (**Fig. 3D**). Finally, using NicheNet (Browaeys et al., 2020) to determine ligand-receptor activity in neutrophils and monocytes for viable versus DES samples, we identified several key shifts in intercellular signaling, many of which aligned with our previous findings (**Fig. 3E**). Of note, amongst the top 15 ligands signaling to receptors on viable neutrophils were Ccl3/4/5—Ccr1 and Il4—Il13ra1 compared to Dll1—Notch2, F11r/Icam1—Itgb2, and Cd34—Sell in DES neutrophils (**Fig. 3E**). For the same analysis in monocytes, top 15 ligands in viable samples included Ccl3—Ccr1, Csf1—Csf1r, and Ccl2/Cxcl12—Ccr2 compared to Icam1—Itgb2 and Cd34—Sell in DES monocytes (**Fig. 3E**).

In addition to immune cell activation, markers of cellular stress were increased in viable samples. Among others, these included markers of endoplasmic reticulum stress (**Fig. 3B**) as well as robust upregulation of ubiquitin, an essential component of protein degradation. Across the majority of clusters, both viable samples showed a consistent increase in genes encoding the two key subunits of ubiquitin, Ubiquitin B (Ubb) and Ubiquitin C (Ubc; **Fig. 3F**). To validate whether ubiquitination was indeed increased at the protein level following tissue dissociation, we isolated femurs from individual mice, cut the bones in half, and briefly centrifuged the bone marrow into 3 separate solutions: (1) DES, (2) lysis buffer, or (3) Ficoll (**Fig. 3G**). Bone marrow from the Ficoll group was immediately dissociated via pipetting in FACS buffer and then stored on ice for 3 hours (h) before being washed and lysed in lysis buffer, thus simulating our scRNA-seq experiment. Immunoblotting for ubiquitin revealed a strong, consistent, and significant increase in total protein ubiquitination in the dissociated samples (**Fig. 3H**). Importantly, this increase in ubiquitination was absent from both the direct lysis group as well as the DES fixed group, confirming that DES prevents cellular stress responses that occur only post-dissociation.

### Preservation of Sensitive Cell Types using DES Fixation

Most tissues contain one or more highly sensitive cell populations that are especially prone to dissociation-induced injury, and thus, are poorly represented in scRNA-seq. In the bone marrow, one such population is the LEPR+ stroma, which act as key regulators of hematopoiesis through the production of critical niche factors, like Cxcl12 and Kitlg (Morrison and Scadden, 2014). One potential reason why LEPR+ stroma are especially sensitive to dissociation is their delicate morphology. Using Adipoq-TdTomato reporter mice and leptin receptor (LEPR) staining in PFA-fixed sections, bone marrow stroma were visible as a physical network of ramified cells with numerous cellular processes that directly contacted hematopoietic and non-hematopoietic cells (**Fig. 4A**). Fixation before dissociation using DES, followed by imaging flow cytometry, revealed similar appearing TdTomato+ cells with ramified morphologies of varying degrees (**Fig. 4B**), suggesting that DES preserves, at least, the *in vivo* morphology of bone marrow stroma.

Given the delicate structure of LEPR+ stroma, we hypothesized that enzymatic dissociation, as opposed to mechanical, would reduce shearing and thereby result in increased cell recovery. To this end, we isolated femurs from individual Adipoq-TdTomato mice and quantified the viability of the TdTomato+ stroma using three different dissociation methods (**Fig. 4C**). One femur was crushed with a mortar and pestle, and the isolated bone marrow was further dissociated into a single-cell suspension via pipetting (**Fig. 4C**). The remaining femur was cut in half and the bone marrow was isolated by rapid centrifugation into Ficoll, followed by either brief enzymatic dissociation or mechanical dissociation in FACS buffer (**Fig. 4C**). Flow cytometry for viability staining definitively showed that enzymatic dissociation improved the recovery of viable stroma, with both methods of mechanical dissociation resulting in roughly 65-70% loss in viability whereas ∼85% of stroma were viable using the enzymatic method (**Fig. 4D**).

For the two viable samples in our bone marrow scRNA-seq dataset, the fraction used for stroma FACS sorting was derived from enzymatically dissociated bone marrow, whereas the CD45+ fraction was isolated with mechanical dissociation (**Fig. 2C**; supplemental methods). To compare the transcriptomes of viable versus DES stroma, we combined the stromal cells from the aforementioned dataset with those from an earlier optimization study where stromal cells were isolated from a single animal using either DES or the mortar-based method (**Supp. Fig. 6A-C**). UMAP dimensionality reduction revealed one main population of stroma connected to a smaller subpopulation, which was derived almost exclusively from the viable enzymatic samples (**Fig. 4E**), similar to the original global clustering (**Fig. 2D-E**). Few stromal cells were isolated from the mechanically dissociated sample (**Fig. 4E**), consistent with our flow cytometry viability findings (**Fig. 4D**). Comparing cell densities within UMAP space revealed major shifts between DES and viable stroma (**Fig. 4F**), consistent with a potential differentiation trajectory.

**Figure 6.**
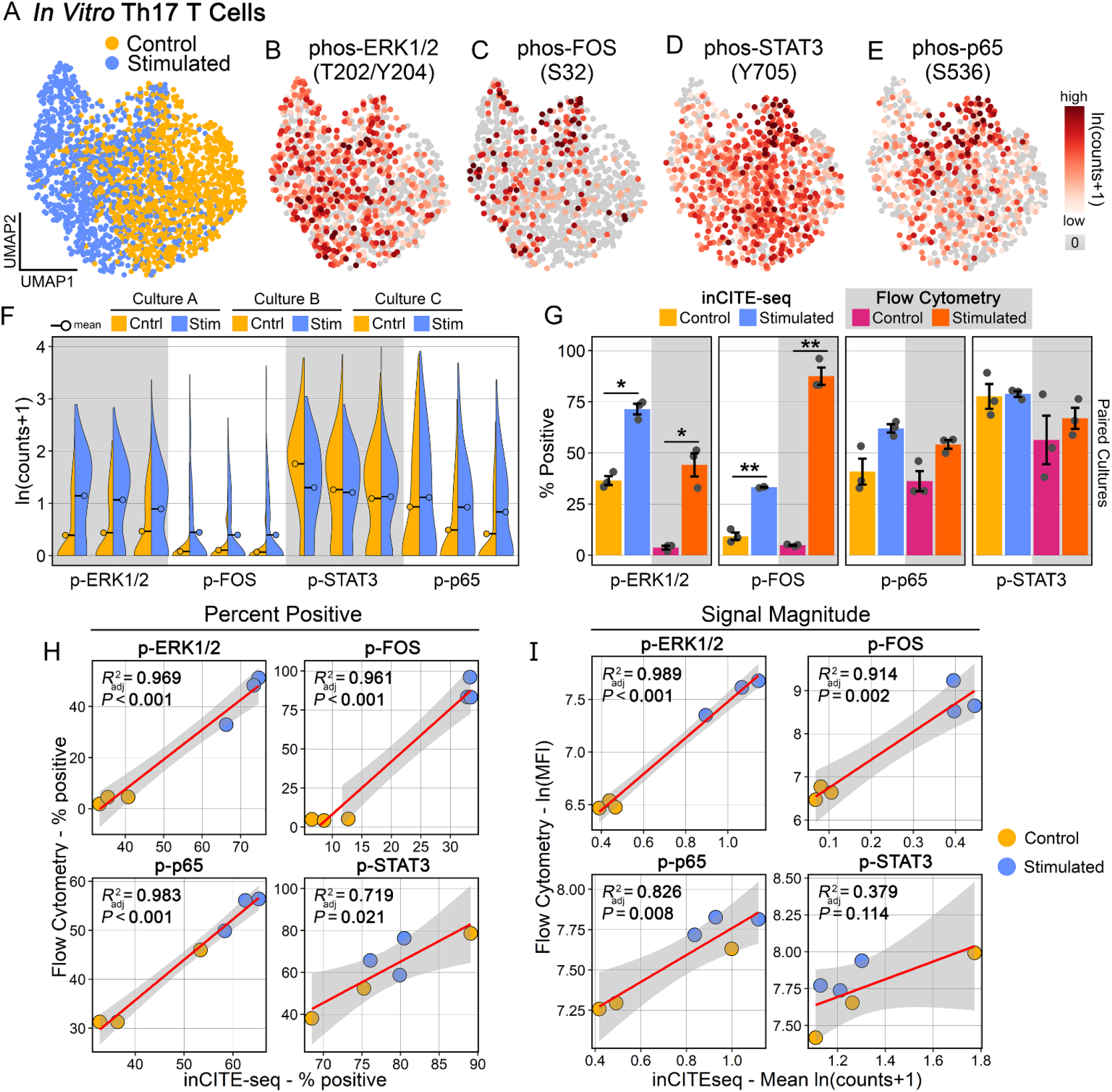
Validation of intracellular CITE-seq for four phospho-targets during Th17 stimulation. (**A**) UMAP of scRNA-seq data from 2,230 cells annotated according to stimulation conditions. (**B-E**) UMAP plots of log normalized counts from inCITE-seq for: (**B**) phospho-ERK1/2 (T204/Y204), (**C**) phospho-FOS (S32), (**D**) phospho-STAT3 (Y705), and (**E**) phospho-p65 (S536). (**F**) Comparisons of log normalized counts showing stimulated versus control for each of the 4 phospho-targets. Split violins represent paired cultures. (**G**) Comparisons of percent positivity for the 4 phospho-targets using inCITE-seq versus flow cytometry from the same cultures. Positive gating for inCITE-seq was determined as any cell with >0 phospho-target counts after isotype subtraction and natural log (ln) +1 scaling. One way ANOVA with Sidak’s multiple comparisons test. * p < 0.05; ** p ≤ 0.01. (**H**) Linear regressions for the 4 phospho-targets comparing percent positivity (**G**) from inCITE-seq versus flow cytometry. (**I**) Linear regressions for the 4 phospho-targets comparing signaling magnitude from inCITE-seq (ln(counts-isotype +1)) versus flow cytometry (mean fluorescent intensity). MFI, mean fluorescent intensity.

Next, we performed a differential gene expression analysis comparing viable versus DES stroma. In total, 5,490 DGs were identified, including 3,020 upregulated and 2,470 downregulated DGs (**Fig. 4G**). Among the top upregulated DGs in viable stroma were numerous mitochondrial and ribosomal genes. We compared the mitochondrial and ribosomal percentages of the total transcriptome, standard quality control metrics in scRNA-seq, and confirmed that viable stroma had increased representation of both (**Fig. 4H-I**). Beyond mitochondrial and ribosomal genes, several stress and activation markers were upregulated in viable stroma, including heat-shock proteins (Hsp90ab1, Hsp90b1, Hspa8, Hspa5), ubiquitin (Ubb, Ubc), interferon-inducible genes (Ifitm3, Ifitm2), and cytokines like Cxcl14 (**Fig. 4G**). Conversely, downregulated DGs in viable stroma included several homeostatic markers (Pdgfrb, Runx2, Notch2, Pparg), cell cycle genes (Top2a, Cdk8, Mki67), and a variety of growth factors (Vegfc, Fgf13, Ctgf, Ngf, Fgf7; **Fig. 4G**). Excluding mitochondrial and ribosomal gene sets, top pathways from gene set enrichment analysis (GSEA) included several related to ubiquitination, antigen presentation, immune-mediated cytotoxicity, and apoptosis (**Fig. 4J**).

Lastly, we performed an RNA velocity analysis of the bone marrow stroma. Pseudotime reconstruction revealed a terminal state within the small subpopulation that was specific to the viable samples (**Fig. 4K**). Overlaying RNA velocity and partition-based graph abstraction (PAGA) with subclustering revealed 3 distinct clusters that were connected by a differentiation trajectory that terminated with the aforementioned subpopulation (**Fig. 4L**). Using only the viable stroma, we performed differentiational gene expression analyses comparing each of the 3 clusters (**Supp. Fig. 6D**) and then plotted the DGs along pseudotime. DGs that were specific to the initial cell state (cluster 1) included several of the downregulated DGs found in the viable stroma (**Fig. 4M**). Similarly, the DGs that were specific to the intermediate (cluster 2) and terminal (cluster 3) states were many of the same upregulated DGs found in viable stroma (**Fig. 4M**). Thus, we have identified an artifactual activation signature in LEPR stroma caused by tissue dissociation, suggesting that fixation before dissociation using DES better preserves the *in vivo* phenotype of sensitive cell types.

### DES Retains Post-Translational Modifications and Allows Intracellular Staining

We next explored whether DES is capable of preserving *in vivo* post-translational modifications (PTMs), like phosphorylation and acetylation. These dynamic processes are key indicators of active intracellular signaling but are generally unstable and sensitive to perturbations (Baker et al., 2005). To this end, we performed total protein immunoblotting for acetyl-lysine (**Fig. 5A**) and phospho-threonine (**Fig. 5B**) from paired whole bone marrow using the same experimental design described previously (**Fig. 4G**). For acetylation, a mild but significant decrease was observed following tissue dissociation compared to direct lysis in non-fixed samples (**Fig. 5A,C**). DES preserved more acetylation across the proteome (**Fig. 5A**), resulting in a slight but significant increase in total acetylation compared to non-fixed direct lysis samples (**Fig. 5C**). For phosphorylation, tissue dissociation caused loss at particular kDa’s across the proteome (**Fig. 5B**), but this was not significant for overall total phosphorylation compared to direct lysis samples (**Fig. 5C**). Conversely, DES provided robust preservation of phosphorylation across the proteome (**Fig. 5B**), resulting in a highly significant increase in total recovered phosphorylation compared to direct lysis samples, which contained phosphatase inhibitors in the lysis buffer (**Fig. 5C**). These data indicate that DES is capable of preserving another level of *in vivo* biological information, active intracellular signaling, and does so to a degree superior to that of current gold-standard phosphatase inhibitors.

An empirical observation of DES fixation is that it provides sufficient membrane permeabilization for intracellular staining. In scRNA-seq, this resulted in moderate increases in ambient RNA, mostly from contaminating erythrocytes, which was sufficiently removed with machine learning software that models the ambient profile of empty droplets (Sheng et al., 2022). More intriguingly, DES permeabilization presents an opportunity to quantify intracellular signaling in scRNA-seq, thereby potentially allowing the parallel integration of *in vivo* transcription and protein signaling in single-cells. To interrogate whether DES could be used to study the phosphorylation of particular protein species in single-cells, we first assessed the reliability of intracellular staining with DES compared to current gold-standard methods. To do this, we used *in vitro* differentiated mouse Th17 T cells, which were cultured in Th17 polarizing conditions followed by a secondary stimulation in half the cells (**Fig. 5D**). Using Il17a-GFP transgenic mice, we compared flow cytometry quantifications of double positive GFP+/anti-IL17A+ cells in identical cultures under stimulated versus control conditions using either DES or standard PFA/methanol (**Fig. 5E**; **Supp. Fig. 7A-C**). Both methods resulted in similar recovery of double positive cells (**Fig. 5E**), which were highly significant compared to paired control cultures (**Fig. 5F**). Importantly, linear regression comparing the double positive percentage of each culture for DES versus PFA/methanol revealed a robust and highly significant correlation (R^2^=0.955; **Fig. 5G**), demonstrating that intracellular staining with DES is reproducible and reliable compared to standard methods.

With this validation, we next assessed the ability of DES to preserve the phosphorylation of individual protein species in single cells using flow cytometry. We again utilized *in vitro* differentiated Th17 cultures and compared the levels of 14 different intracellular targets, including 13 distinct phospho-species, between stimulated and unstimulated cells (**Fig. 5H**; **Supp. Fig. 7D**). Results between paired cultures and replicates were highly consistent and revealed increased phosphorylation in several species, including significant increases in both antibody clones to p-ERK1/2 (T202/Y204), p-FOS (S32), and non-canonical p-STAT1 (S727), but not canonical p-STAT1 (Y701), which was inhibited by neutralizing anti-IFNγ within the cultures (**Fig. 5H**)(Harrington *et al*., 2005).

Other phospho-species that showed consistent increasing trends with stimulation but were not significant after correction for multiple comparisons included p-p65 (S536), p-CREB (S133), and non-canonical p-STAT3 (S727) (**Fig. 5H**). Given that phorbol myristate acetate (PMA), a component of our secondary stimulation (PMA/ionomycin), drives phosphorylation of ERK1/2 via protein kinase C (Niedel et al., 1983), we included 2 different clones to p-ERK1/2 as positive controls, both of which showed significant increases (**Fig. 5I**). The single phospho-species with the largest increase between stimulated and control cultures was p-FOS (S32), which in total increased from 4.57% to 85.6% (**Fig. 5I**). Together, these data demonstrate that DES preserves the 3 main PTMs found on proteins, ubiquitination, acetylation, and phosphorylation, and provides sufficient membrane permeabilization for the quantification of individual phospho-species in single-cells using immunostaining.

### Quantification of Phospho-Signaling with inCITE-seq in DES-Fixed Cells

A key domain of biological information that is lost with current scRNA-seq methods is intracellular protein signaling and in particular, phospho-signaling, which regulates the majority of pathways in biology. Given that DES preserves PTMs and allows immunostaining of intracellular epitopes, we next sought to develop an optimized, scalable, and modular approach to inCITE-seq in DES fixed samples (**Supp. Fig. 8A**). To this end, we used a photo-crosslinking small peptide (oligo-IgG BP) that binds with high affinity and specificity to the Fc region of immunoglobulins and conjugated it with a modified TotalSeq™-B oligonucleotide (oYo-link^®^). Following incubation in the presence of UV light, single micrograms of antibody can be covalently conjugated with site-specificity and high efficiency (**Supp. Fig. 8B**).

Optimization studies suggested that high levels of background staining resulted from unbound residual IgG BP as well as off-target interactions between the ssDNA oligonucleotide and endogenous intracellular molecules. To circumvent these issues, we used magnetic beads containing human IgG Fc fragments to remove unbound oligo-IgG BP (**Supp. Fig. 8C**) and E. coli ssDNA BP to block oligonucleotide interactions (**Supp. Fig. 8A**), as was recently demonstrated (Chen et al., 2022).

As a test case for inCITE-seq in DES fixed cells, we used the same three Th17 cultures under stimulated and control conditions from (**Fig. 5H-I**) and stained four phospho-targets, p-ERK1/2 (T202/Y204), p-FOS (S32), p-STAT3 (Y704), and p-p65 (S536) as well as an isotype control antibody. To minimize technical variation, we controlled for the number of cells during immunostaining, used the same antibody cocktails, and hash-tagged the culture replicates into one sample for scRNA-seq. After quality control filtering and dimensionality reduction 2,230 total cells were recovered, with a clear separation between control and stimulated cells (**Fig. 6A**). For processing of inCITE-seq, isotype control counts were subtracted from each target followed by Log+1 transformation. Staining distributions for each of the 4 phospho-species, p-ERK1/2 (**Fig. 6B**), p-FOS (**Fig. 6C**), p-STAT3 (**Fig. 6D**), and p-p65 (**Fig. 6E**) revealed distinct patterns, some of which had overlapping features with other phospho-targets.

Comparing stimulated and control conditions in each of the 3 paired cultures revealed a consistent increase in phospho levels for p-ERK1/2, p-FOS, and p-p65, but not p-STAT3 (**Fig. 6F**), mirroring our previous findings for the same targets in the same cultures using flow cytometry (**Fig. 5H**). Of note, p-STAT3 was maintained at a high level independent of PMA/ionomycin stimulation, consistent with IL6-pSTAT3 signaling from the culture media, an essential component of Th17 maintenance (Harbour et al., 2020). We next determined the percentage of positive staining for inCITE-seq in each of the culture/conditions for the four phospho-targets (defined as any cell with >0 counts for a target after isotype correction) and compared this to the percent positive as determined by flow cytometry for the same targets in the same cultures. In agreement, stimulated cultures showed significant increases for p-ERK1/2 and p-FOS with both flow cytometry and inCITE-seq, while neither p-STAT3 nor p-p65 showed significant shifts with either method (**Fig. 6G**). Finally, using linear regressions to compare percent positivity for paired culture/conditions between inCITE-seq and flow cytometry revealed robust and significant positive correlations for all 4 targets (p-ERK1/2 R^2^=0.969; p-FOS R^2^=0.961; p-p65 R^2^=0.983; p-STAT3 R^2^=0.719; **Fig. 6H**).

Similarly, linear regressions for signal magnitude between inCITE-seq and flow cytometry, using log counts and mean fluorescent intensity (MFI), respectively, showed strong positive correlations, with the exception of p-STAT3, which was not significant (p-ERK1/2 R^2^=0.989; p-FOS R^2^=0.914; p-p65 R^2^=0.826; p-STAT3 R^2^=0.379; **Fig. 6I**).

These data provide definitive validation that phosphorylated proteins can be reliably quantified with inCITE-seq following DES fixation.

### Parallel Integration of Transcription and Phospho-Signaling during Th17 Stimulation Reveals Hyperactivated p-ERK1/2 p-FOS Double Positive Cells

Our secondary stimulation using PMA/ionomycin functions via phosphorylation of ERK1/2 and calcium influx, respectively (Feske et al., 2001; Niedel et al., 1983). Therefore, as a positive control, we expected that DGs from phosphorylated versus unphosphorylated ERK1/2 would be contained within the DG gene set from stimulated versus unstimulated cells. In agreement, all 497 DGs identified in p-ERK1/2+ cells overlapped with DGs from the stimulated gene set. Among p-ERK1/2+ cells, top upregulated DGs included several key genes involved in Th17 activation including Il17a, Il17f, Tnf, Il2, Cd40lg, Tnfrsf4, and CD69, among others (**Fig. 7A; Supp. Fig. 9A**) (Hirota et al., 2007). Downregulated genes were enriched for mitochondrial transcripts as well as markers of resting T cells including Tcf7, Ccr7, and Sell (**Fig. 7A**) (Szabo et al., 2019). Il2 was the most highly upregulated gene in p-ERK1/2+ cells (**Fig. 7A**), consistent with its central role in early T cell activation (Krönke et al., 1985), and Pearson correlation analysis demonstrated a positive linear relationship between p-ERK1/2 levels and Il2 gene expression (**Supp. Fig. 9B**).

**Figure 7.**
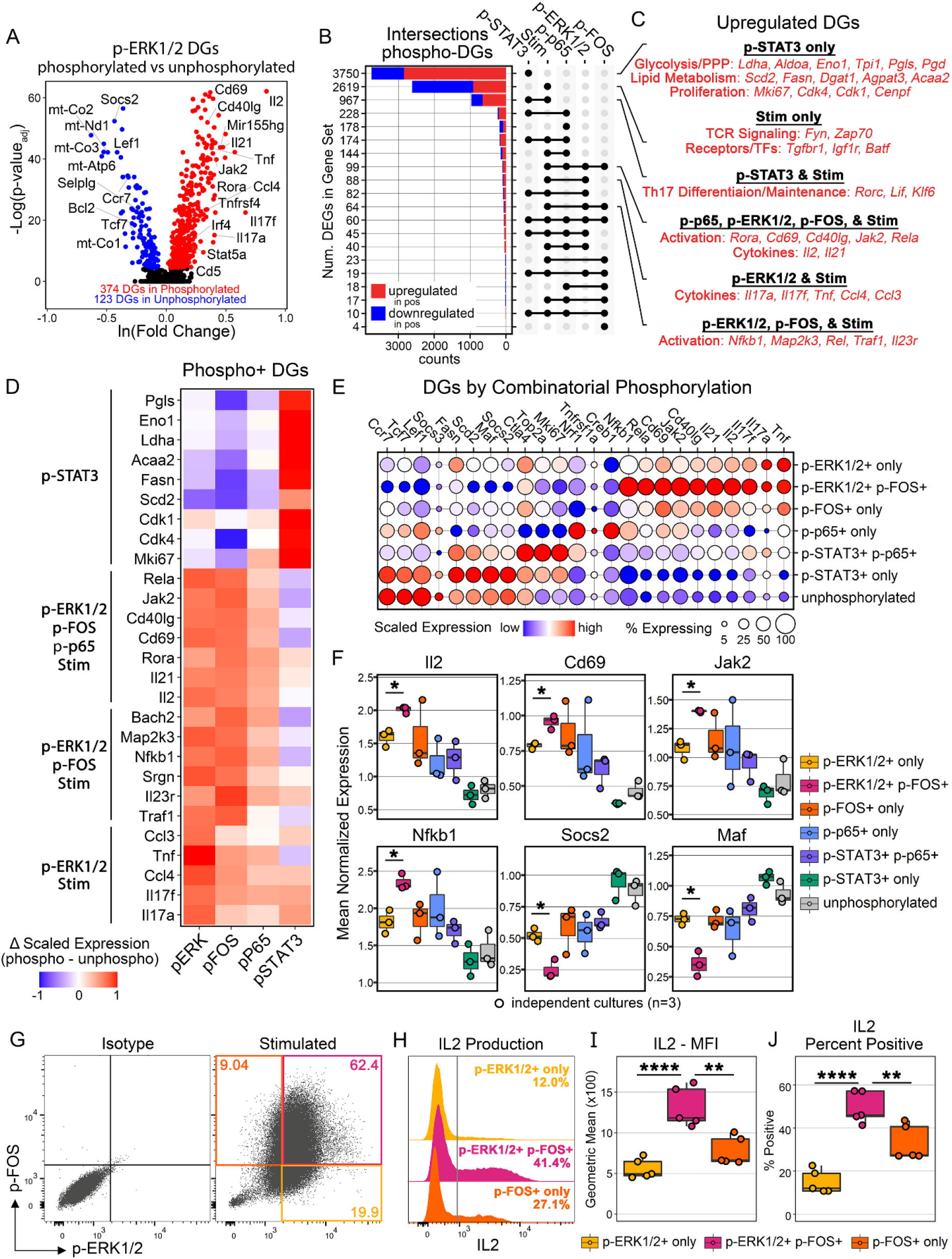
Parallel integration of transcriptomes and phospho-signaling in Th17 stimulation identifies hyperactivated pFOS+/pERK1/2+ double-positive cells. (**A**) Volcano plot for p-ERK1/2 DGs (phosphorylated p-ERK1/2 versus unphosphorylated p-ERK1/2). (**B**) Upset plot showing intersections of DG gene sets from each of the 4 phospho-targets and stimulation. Stimulation refers to stimulated versus unstimulated cells. p-ERK1/2=497 DGs. p-FOS=318 DGs. p-p65=1,013 DGs. p-STAT3=5,339 DGs. Stimulation=4,454 DGs. Stim, stimulation. (**C**) Selected upregulated DGs from 6 different gene sets. (**D**) Heatmap of selected DGs from (**C**) showing the change in scaled expression between phosphorylated and unphosphorylated cells with respect to each of the 4 phospho-targets. (**E**) Combinatorial phosphorylation analysis showing selected top DGs from each combination of phospho-targets. inCITE-seq was performed with 2 panels of antibodies and combinations are restricted to the 2 panels. Panel 1=p-STAT3 and p-p65. Panel 2=p-ERK1/2 and p-FOS. (**F**) Selected combinatorial phosphorylation DGs from (**E**) showing culture-wise mean expression. One-way ANOVA with FDR correction. * p < 0.05. (**G-J**) Flow cytometry from Th17 validation cohort (n=5 independent cultures). (**G**) Representative p-FOS and p-ERK1/2 staining in PFA/methanol fixed cells showing gating of 3 populations: p-FOS+ p-ERK1/2-, p-FOS+ p-ERK1/2+, and p-FOS-p-ERK1/2+. (**H**) Histograms showing IL2 staining in the same 3 populations from (**G**). Gray line indicates isotype gating for IL2 staining. (**I**) IL2 mean fluorescent intensity (MFI) and (**J**) percent positive staining across the validation cohort. One-way ANOVA with Tukey’s post-hoc test. ** p ≤ 0.01; **** p ≤ 0.0001.

Next, to determine if specific phosphorylation events were associated with discrete gene sets during Th17 activation, such as cytokine production, we computed DGs for each of the four phospho-targets (**Supp. Fig. 9C-F**) and compared the intersections of these DGs with those from stimulated versus unstimulated cells. In total, 22 unique gene sets were identified. The largest gene set was specific to p-STAT3 (**Fig. 7B**), with the most upregulated DG being Lgals1, a recently described direct target of STAT3 (Sharanek et al., 2021). Consistent with prior reports (Berod et al., 2014; Shi et al., 2011), additional upregulated DGs in the p-STAT3 gene-set included proliferation markers (Mki67, Cdk4, Cdk1, Cenpf), lipid metabolism mediators (Scd2, Fasn, Dgat1, Agpat3, Gpat4, Acaa2, Acot7, Fabp5), and glycolysis and pentose phosphate pathway genes (Ldha, Aldoa, Eno1, Tpi1, Pgls, Pgd) (**Fig. 7C-D**). The second largest gene set was specific to stimulation only (**Fig. 7B**) and included several ribosomal genes as well as TCR signaling proteins (Fyn, Zap70), cytokine/growth factor receptors (Tgfbr1, Igf1r), and the transcription factor Batf (**Fig. 7C-D**). The largest shared gene set was between p-STAT3 and stimulation (**Fig. 7B**) and included key genes involved in Th17 differentiation and maintenance such as Rorc, Lif, and Klf6 (Ciofani et al., 2012; Ivanov et al., 2006) (**Fig. 7C-D**). Finally, the gene set shared between p-ERK1/2 and stimulation contained several prominent cytokines like Il17a, Il17f, Tnf, Ccl3, and Ccl4 (**Fig. 7C-D**), whereas the gene set shared between p-p65, p-ERK1/2, p-FOS, and stimulation contained mostly markers of T cell activation such as Cd69, Cd40lg, Il2, Jak2, and Rela (**Fig. 7C-D**) (Hirota et al., 2007).

We next examined whether specific combinations of phosphorylation events were associated with unique transcriptional states during Th17 activation. To this end, we annotated cells as either unphosphorylated, singly phosphorylated, or doubly phosphorylated for each of the two panels of antibodies used in our inCITE-seq experiment (panel 1: p-STAT3 and p-p65; panel 2: p-ERK1/2 and p-FOS). We computed DGs for each phospho combination and compared their gene expression across combinatorial groups, revealing distinct patterns in transcription (**Fig. 7E**).

Compared to p-ERK1/2 only or p-FOS only, p-ERK/p-FOS double positive cells showed the highest expression of Th17 activation genes, including Il17f, Il2, Il21, Cd40lg, Cd69, Mir155hg, Jak2, Rela, and Nfkb1 (**Fig. 7E**) (Hirota et al., 2007). Similarly, the highest expression of proliferation genes, like Mki67 and Top2a, were seen in p-STAT3/p65 double positive cells (**Fig. 7E**). p-STAT3 only cells were uniquely enriched in lipid metabolism genes (**Fig. 7E**). Unphosphorylated cells expressed the highest levels of resting markers, like Ccr7, Tcf7, and Lef1 (**Fig. 7E**) (Szabo et al., 2019). Comparing mean expression for each phospho-combination across the three culture replicates revealed a significant increase in several of the aforementioned activation markers in p-ERK/p-FOS double positive cells compared to p-ERK1/2 alone, including Il2, Cd69, Jak2, and Nfkb1 (**Fig. 7F**). Further, p-ERK/p-FOS double positive cells showed significant decreases in levels of suppressive mediators, like Maf and Socs2, compared to p-ERK1/2 alone (**Fig. 7F**) (Aschenbrenner et al., 2018; Kaufmann et al., 2019). These data suggest that p-ERK/p-FOS double positive cells possess a unique gene expression program associated with hyperactivation in Th17 T cells.

Finally, to experimentally validate that p-ERK/p-FOS double positive Th17 cells display heightened activation, we used flow cytometry to examine IL2 production with respect to p-ERK1/2 and p-FOS staining. In a validation cohort of stimulated cells processed with standard PFA/methanol fixation, we gated on p-ERK1/2- p-FOS+, p-ERK1/2+ p-FOS+, and p-ERK1/2+ p-FOS-cells to compare IL2 staining (**Figure 7G**). In agreement with our inCITE-seq findings, p-ERK/p-FOS double positive cells had the greatest IL2 production amongst all three groups, and both IL2 mean fluorescent intensity (MFI) (**Figure 7H-I**) and IL2 percent positivity (**Figure 7J**; **Supp. Fig. 10B**) showed highly significant increases in double positive cells compared to either phospho-species alone. Thus, using inCITE-seq, we have identified distinct transcriptional signatures derived solely from patterns in phospho-signaling. This included a discrete and experimentally validated cell state, based on combinatorial phosphorylation, associated with heightened IL2 production during Th17 stimulation. In total, these data provide a first-of-its-kind demonstration that droplet-based scRNA-seq can be reliably integrated with phospho-signaling using DES fixation. We confirm that this previously hidden layer of biological information in single-cell sequencing is instructive of cell phenotype.

## DISCUSSION

Single-cell sequencing presents the opportunity to deconstruct cellular networks across health and disease, but key limitations remain, namely loss of biological data, which includes entire information levels like phospho-signaling, as well as loss of *in vivo* cell states caused by tissue dissociation and biological decay. Herein, we present the concept of fixation before dissociation using a next generation preservative that retains *in vivo* biology and allows the parallel quantification of transcription and intracellular signaling in single cells. We demonstrate that DES preserves multiple levels of biological information, including morphology, RNA, proteins, and post-translational modifications. In scRNA-seq, two challenging cell populations were effectively absent with viable dissociation but preserved with DES: macrophages, recently shown to fragment during bone marrow dissociation (Millard et al., 2021), and eosinophils, cells with uniquely high RNase expression posing historical challenges to sequencing (Diny et al., 2022). Comparing viable to DES bone marrow revealed several detrimental effects of dissociation, including immune cell polarization, global stress responses, and artificial activation states in highly sensitive cells. Further, we show that DES provided robust stabilization of phosphorylation, both *in vivo* and *in vitro*, and allowed sufficient permeabilization for intracellular immunostaining. Finally, we validated an optimized and flexible method for performing intracellular CITE-seq in DES fixed cells, which permitted the parallel quantification of transcriptomes and phosphorylated signaling pathways in single-cell sequencing. Applying this method to four key phosphorylated targets during Th17 stimulation revealed a novel, hyperactivated subpopulation, containing both p-ERK1/2 and p-FOS, which had the overall highest expression of activation genes, like Il2, Il17f, Il21, Cd69, and Cd40lg, and the lowest expression of suppressive markers like Socs2 and Maf. Finally, we experimentally validated that p-ERK1/2+ p-FOS+ cells demonstrate heightened activation by showing that this subset had the greatest IL2 production during Th17 stimulation. We anticipate that the use of DES-based fixatives in single-cell technologies will broadly improve the modeling of cellular networks by preserving *in vivo* biology and allowing the parallel integration of transcription and intracellular signaling.

DES fixation coupled with inCITE-seq has the potential to transform our understanding of intracellular signaling by illuminating cooperativity amongst pathways with high resolution and in a massively parallel fashion. To date, many ligand-mediated signaling pathways have been mapped, such that we know a given receptor’s intracellular signaling domains, the peptide signaling motifs that make up those domains, and the signaling proteins that are recruited by a given motif (Daniels et al., 2022; Pawson and Nash, 2003). However, the current understanding of how intracellular signaling shapes cell phenotype is fragmented due to the reliance on reductionist approaches that focus on individual pathways (Hyduke and Palsson, 2010). While these experiments are necessary to understand the discrete components of signaling networks, the generalizability of such results is challenged by the nature of intracellular signaling, which is context-dependent (i.e. the signaling machinery present within a given cell), cooperative in nature (i.e. the interactions between discrete signaling cascades), and highly modular (i.e. the recombining of a fixed set of signaling modalities). For example, there are at least 1,894 literature-supported ligand-receptor pairs in humans (Ramilowski et al., 2015), with >10,000 more predicted (Browaeys et al., 2020). Yet, across eukaryotes, only 178 ligand-binding signaling motifs have been recognized (Kumar et al., 2021), many of which show redundancy in binding partners and thereby the signaling cascades that are modulated. Such convergence onto a comparatively small diversity of signaling machinery suggests that the summation of intracellular signaling may be more generalizable than the individual components, with respect to the emergence of cell phenotype. Using inCITE-seq with DES fixation, we anticipate that entire signaling cascades will be able to be quantified in parallel, from phosphorylated receptors to downstream signaling targets and differential sites of phosphorylation. In the context of neural networks, simultaneously integrating such intracellular information with other single-cell modalities, like transcription (scRNA-seq) and chromatin accessibility (scATAC-seq), presents an opportunity to unravel how cooperativity in signaling networks shapes cell phenotypes in health and disease.

It should be noted that our group is not the first to perform intracellular CITE-seq (Chen et al., 2022; Chung et al., 2021; Mimitou et al., 2021). Moreover, one other article has reported inCITE-seq for phosphorylated proteins using droplet-based single-cell (Rivello et al., 2021). However, validation against gold-standard methods is lacking, and the technique requires harsh fixation using a cross-linking agent, which resulted in poor RNA recovery in cell lines (<1,000 median genes/cell) and unusable RNA in primary cells (<100 median genes/cell). Fixation before dissociation using DES has shown the ability to retain *in vivo* phosphorylation, while simultaneously preserving RNA integrity.

Lastly, DES allows the long-term preservation of biological specimens, which we anticipate will be useful for establishing biobanks of human tissues and other precious samples. For example, we demonstrated at least equivalent RNA quality and quantity in samples stored in DES for 1 month. Moreover, the cells used for inCITE-seq were stored at -80°C for 2 months, during which aliquots were used for multiple experiments. This allowed us to first screen several phospho-species in our Th17 T cells, leading to the identification of unexpected candidate targets like p-FOS, which were then explored in further detail with inCITE-seq using the same samples, ultimately leading to the identification of p-FOS/p-ERK hyperactivated cells. Thus, DES allows the long-term preservation of RNA, proteins, and post-translational modifications and can be used to improve the efficiency of precious biological samples.

### Limitations

While the conclusions herein are rigorous and validated, some important limitations of our study should be noted. Due to constraints from sequencing costs, we chose a single tissue type for our head-to-head comparison of viable versus DES scRNA-seq. Bone marrow represented a favorable test case because it contains hematopoietic and nonhematopoietic cells, including several that have posed prior challenges to sequencing. Nonetheless, we have successfully used fixation before dissociation with DES in scRNA-seq from other tissues, which are outside of the scope of this report. Another limitation is that our inCITE-seq experiment had a relatively low number of recovered cells, which reduced the statistical power of some of our analyses. We hashed 12 inCITE-seq samples into one 10X Genomics well as an effort to control for technical variables. Lastly, we recovered slightly less RNA in DES fixed bone marrow with scRNA-seq but equal or more RNA using standard benchtop methods. This may be due, in part, to the fact that the DES samples had a relatively short duration of fixation (2 hours), which was necessary given our comparison to paired viable bone marrow. We have shown that long-term storage in DES at -80°C resulted in improved recovery of nascent RNA. Therefore, it is likely that a longer duration of fixation would have increased transcript recovery, but this would have required performing scRNA-seq on separate days for paired samples, thereby introducing additional technical covariates.

### STAR★METHODS

Detailed methods are provided in the online supplement and include the following:

- KEY RESOURCES TABLE
- RESOURCE AVAILABILITY
  - Lead Contact
  - Materials Availability
  - Data and Code Availability
- EXPERIMENTAL MODELS AND SUBJECT DETAILS
  - Mice
  - Th17 T Cell Culture and Stimulation
  - Human Retina
- METHOD DETAILS
  - DES Fixation
  - Dissociation of DES-Fixed Tissues
  - Transfer from DES to Aqueous Buffer,
  - Blocking, and Immunostaining
  - Bone Marrow RNA Electrophoresis
  - RT-qPCR, RNA Integrity, and RNA Quantity in Th17 T cells
  - Imaging Flow Cytometry
  - Confocal Immunofluorescence
  - Bone Marrow Single-Cell RNA-Sequencing
  - Immunoblotting for Bone Marrow Post-
  - Translational Modifications
  - Flow Cytometry for LEPR+ Stroma Viability
  - Th17 Flow Cytometry
  - Oligonucleotide-Antibody Conjugation for Intracellular CITE-Sequencing
  - Th17 T Cell Single-Cell RNA-Sequencing
- QUANTIFICATION AND STATISTICAL ANALYSES
  - Bone Marrow Single-Cell RNA-Sequencing
  - Pearson Correlation Analyses
  - Differential Gene Expression Analyses
  - Intersection Analyses of Differentially Expressed Genes
  - Pathway Enrichment Analyses
  - Transcription Factor Activity Analyses
  - Ligand-Receptor Analyses
  - RNA Velocity and Pseudotime Analyses
  - Analyses of Th17 T Cell intracellular CITE-Sequencing
  - Th17 Single-Cell RNA-Sequencing: d0 versus d60
  - Analyses of Flow Cytometry Data
  - Statistics and Data Visualizations
- SUPPLEMENTAL FIGURES

## ACKNOWLEDGMENTS

The authors would like to acknowledge the UAB Core Grant for Vision Research, P30-EY003039, the Carl G. and Pauline Buck Trust, and the UAB Flow Cytometry & Single Cell Core Facility, supported by the Center for AIDS Research, AI027767, and The O’Neal Comprehensive Cancer Center, CA013148. S.D.F. is supported by NEI, NIH F30-EY033198. B.F.F. is supported by NIDDK, NIH F30-DK127865. P.B.F. is supported by the Mark Foundation for Cancer Research. C.T.W. is supported by NIAID, NIH R01-AI164772, and U01-AI163063. M.B.G. is supported by NEI, NIH R01-EY033620, R01-EY032753, R01-EY025383, R01-EY028858, and R01-EY012601. R.S.W. is supported by NHLBI, NIH R01-HL150078 and the Mark Foundation for Cancer Research.

## AUTHOR CONTRIBUTIONS

SDF: conceptualization, data curation, formal analysis, investigation, methodology, validation, visualization, writing—original draft, writing— review & editing

BFF, VSH, SL: investigation, writing—review & editing

AG, KVK: resources, writing—review & editing

CTW, PBF: supervision, resources, writing— review & editing

RSW, MBG: supervision, funding acquisition, writing—review & editing

## DECLARATION OF INTERESTS

AG is a founder of RNAssist Limited, UK and holds multiple patents for DES including vivoPHIX™ (EP2961268B1, EP3430903B1) and other national equivalents. KVK is an employee at Kodiak Sciences Inc. All other authors declare no competing interests.

