## Supplement - Methods and Supplemental Figures for "Fixation Before Dissociation Using a Deep Eutectic Solvent Preserves *In Vivo* States and Phospho-Signaling in Single-Cell Sequencing"

### STAR★METHODS

### KEY RESOURCES TABLE

| REAGENT or RESOURCE | SOURCE | IDENTIFIER |
|---|---|---|
| | Antibodies | |
| PE anti-CD45 (30F11) | BioLegend | cat: 103105 |
| PE anti-CD45R (B220) | BioLegend | cat: 103207 |
| APC anti-Ter119 (Ter-119) | BioLegend | cat: 116211 |
| AF647 anti-Spectrinβ (B-1) | Santa Cruz | cat: sc-374309 AF647 |
| APC anti-IL17A (eBio17B7) | ThermoFisher | cat: 25-7177-81 |
| PE/Cy7 anti-IL17A (eBio17B7) | ThermoFisher | cat: 25-7177-82 |
| PE anti-IL2 (JES6-5H4) | BioLegend | cat: 503807 |
| PE anti-RORγT (AFKJS-9) | ThermoFisher | cat: 12-6988-82 |
| anti-phos-STAT1 Y701 (D4A7) | Cell Signaling | cat: 7649 |
| anti-phos-STAT1 S727 (D3B7) | Cell Signaling | cat: 8826 |
| anti-phos-STAT3 Y705 (4P/STAT3) | BD Biosciences | cat: 612357 |
| anti-phos-STAT3 S727 (D4X3C) | Cell Signaling | cat: 34911 |
| anti-phos-STAT5 Y694 (D47E7) | Cell Signaling | cat: 4322 |
| anti-phos-AKT T308 (C31E5E) | Cell Signaling | cat: 2965 |
| anti-phos-AKT S473 (D9E) | Cell Signaling | cat: 4060 |
| anti-phos-ERK1/2 T202/Y204 (D13.14.4E) | Cell Signaling | cat: 4370 |
| anti-phos-ERK1/2 T202/Y204 (197G2) | Cell Signaling | cat: 4377 |
| anti-phos-MEK1/2 S221 (166F8) | Cell Signaling | cat: 2338 |
| anti-phos-CREB S133 (87G3) | Cell Signaling | cat: 9198 |
| anti-phos-FOS S32 (D82C12) | Cell Signaling | cat: 5348 |
| anti-phos-JUNB T102/T104 (D3C6) | Cell Signaling | cat: 8053 |
| anti-phos-p65 S536 (93H1) | Cell Signaling | cat: 3033 |
| AF647 anti-phos-FOS S32 (D82C12) | Cell Signaling | cat: 8677 |
| anti-GFP (polyclonal) | Abcam | cat: ab13970 |
| anti-Ubiquitin (E4I2J) | Cell Signaling | cat: 43124 |
| anti-Phospho Threonine (42H4) | Cell Signaling | cat: 9386 |
| anti-Acetylated Lysine (Ac-K-103) | Cell Signaling | cat: 9681 |
| anti-E-cadherin (24E10) | Cell Signaling | cat: 3195 |
| AF647 anti-rhodopsin (RET-P1) | Santa Cruz | cat: sc-57433 AF647 |
| AF647 anti-Iba1 (EPR16588) | Abcam | cat: ab225261 |
| FITC anti-CD45 (30F11) | BioLegend | cat: 103107 |
| Biotinylated anti-LEPR (polyclonal) | R&D Systems | cat: BAF497 |
| AF+488 anti-rabbit (polyclonal) | ThermoFisher | cat: A32790 |
| AF+647 anti-rabbit (polyclonal) | ThermoFisher | cat: A32795 |
| AF+488 anti-chicken (polyclonal) | ThermoFisher | cat: A32931 |
| AF+800 anti-rabbit (polyclonal) | ThermoFisher | cat: A32808 |
| AF+800 anti-mouse (polyclonal) | ThermoFisher | cat: A32789 |

| | | |
|---|---|---|
| APC rat isotype (G155-178) | BD Biosciences | cat: 550882 |
| PE rat isotype (MPC-11) | BioLegend | cat: 400312 |
| AF647 rabbit isotype (DA1E) | Cell Signaling | cat: 2985 |
| Rabbit isotype (DA1E) | Cell Signaling | cat: 3900 |
| Mouse isotype (G3A1) | Cell Signaling | cat: 5415 |
| True-Stain Monocyte Blocker™ | BioLegend | cat: 426102 |
| Mouse TruStain FcX™ anti-CD16/32 (93) | BioLegend | cat: 101319 |
| Mouse TruStain FcX™ PLUS anti-CD16/32 (S17011E) | BioLegend | cat: 156603 |
| Human TruStain FcX™ | BioLegend | cat: 422301 |
| Neutralizing anti-IL4 (11B11) | BioLegend | cat: 504121 |
| Neutralizing anti-IFNγ (XMG1.2) | BioLegend | cat: 505833 |
| anti-CD28 (37.51) | BioLegend | cat: 102115 |
| anti-CD3 (145-2C11) | ThermoFisher | cat: 16-0031-82 |
| TotalSeq™-B0301 anti-mouse Hashtag 1 | BioLegend | cat: 155831 |
| TotalSeq™-B0302 anti-mouse Hashtag 2 | BioLegend | cat: 155833 |
| TotalSeq™-B0303 anti-mouse Hashtag 3 | BioLegend | cat: 155835 |
| TotalSeq™-B0304 anti-mouse Hashtag 4 | BioLegend | cat: 155837 |
| TotalSeq™-B0305 anti-mouse Hashtag 5 | BioLegend | cat: 155839 |
| TotalSeq™-B0306 anti-mouse Hashtag 6 | BioLegend | cat: 155841 |
| TotalSeq™-B0307 anti-mouse Hashtag 7 | BioLegend | cat: 155843 |
| TotalSeq™-B0308 anti-mouse Hashtag 8 | BioLegend | cat: 155845 |
| TotalSeq™-B0309 anti-mouse Hashtag 9 | BioLegend | cat: 155847 |
| **Dyes and Lectins** | | |
| SYTO40 | ThermoFisher | cat: S11351 |
| LIVE/DEAD™ Fixable Near-IR Dead Cell Stain | ThermoFisher | cat: L34975 |
| Hoechst 33342 Solution | BD Biosciences | cat: 561908 |
| Ulex Europaeus Agglutinin I (UEA I), Fluorescein | Vector Labs | cat: FL-1061-2 |
| Peanut Agglutinin (PNA), Biotinylated | Vector Labs | cat: B-1075-5 |
| AF647 Streptavidin | ThermoFisher | cat: S32357 |
| **Chemicals, Peptides, and Recombinant Proteins** | | |
| vivoPHIX™ | Rapid Labs Ltd | cat: RD-VIVO-50 |
| Glacial acetic acid | Fisher Scientific | cat: A38S-500 |
| 20x saturated sodium chloride | Invitrogen | cat: AM9770 |
| Saturated Ammonium Sulfate Solution | Thermo Scientific | cat: 45216 |
| 1x PBS | Corning | cat: 21-040-CV |
| ACK lysis buffer | ThermoFisher | cat: A1049201 |
| CD4+ T Cell Isolation Kit, mouse | Miltenyi | cat: 130-104-454 |
| IMDM media | ThermoFisher | cat: 12440046 |
| Recombinant Mouse IL-6 Protein | R&D Systems | cat: 406-ML-025/CF |
| Recombinant Human TGF-beta 1 Protein | R&D Systems | cat: 240-B-010/CF |
| Phorbol 12-myristate 13-acetate | Millipore Sigma | cat: P8139-1MG |
| Ionomycin calcium salt | Millipore Sigma | cat: I3909-1ML |

| | | |
|---|---|---|
| Fixation/Permeabilization Solution Kit with BD GolgiStop™ | BD Biosciences | cat: 554715 |
| SsoAdvanced Universal SYBR Green Supermix | Bio-Rad | cat: 1725270 |
| RNeasy Plus Mini Kit | Qiagen | cat: 74134 |
| Direct-zol RNA Miniprep | Zymo Research | cat: R2051 |
| Quant-it™ RiboGreen RNA Assay Kit and RiboGreen RNA Reagent, RediPlate™ 96 RiboGreen™ RNA Quantitation Kit | ThermoFisher | cat: R11490 |
| Trident Universal Protein Blocking Reagent (animal serum free) | GeneTex | cat: GTX30963 |
| ProLong™ Gold Antifade Mountant | ThermoFisher | cat: P36930 |
| 1 mL, Open-Top Thickwall Polycarbonate Tube, 8 x 51mm | Beckman Coulter | cat: 355657 |
| Whatman® Nuclepore™ Track-Etched Membranes | Millipore Sigma | cat: WHA10417406 |
| Heparin sodium salt from porcine intestinal mucosa | Millipore Sigma | cat: H3393-100KU |
| RNasin® Plus Ribonuclease Inhibitor | Promega | cat: N2615 |
| Single-Stranded DNA Binding Protein | Promega | cat: M3011 |
| Halt™ Phosphatase Inhibitor Cocktail | ThermoFisher | cat: 78426 |
| Ethylenediamine Tetraacetate Acid (EDTA) | Fisher Scientific | cat: BP2482100 |
| HyClone HyPure Water, Molecular Biology Grade | Cytivia | cat: SH30538.LS |
| Tissue-Tek® O.C.T. Compound | Sakura | cat: 4583 |
| Ficoll® Paque Plus | Millipore Sigma | cat: GE17-1440-02 |
| ULAB Test Tubes with Black Screw Caps, Vol.10ml, Dia.16x100mm with Marking Area, Borosilicate Glass Material | Amazon | cat: B07S77FF5H |
| Collagenase D | Millipore Sigma | cat: 11088866001 |
| Dispase® II (neutral protease, grade II) | Millipore Sigma | cat: 4942078001 |
| Corning® 500 mL MEM (Minimum Essential Medium) | Corning | cat: 10-010-CV |
| TrypLE™ Express Enzyme (1X), no phenol red | ThermoFisher | cat: 12604021 |
| MEM Non-essential Amino Acid Solution (100×) | Millipore Sigma | cat: M7145-100ML |
| GlutaMAX™ Supplement | ThermoFisher | cat: 35050061 |
| Sodium Pyruvate (100 mM) | ThermoFisher | cat: 11360070 |
| DNase I recombinant, RNase-free | Millipore Sigma | cat: 4716728001 |
| RIPA Buffer | Millipore Sigma | cat: R0278-500ML |
| 4X Protein Sample Loading Buffer | Li-Cor | cat: 928-40004 |
| 10x TBS | Bio-Rad | cat: 1706435 |
| Pierce™ Western Blot Signal Enhancer | ThermoFisher | cat: 21050 |
| Intercept® (TBS) Blocking Buffer | Li-Cor | cat: 927-60001 |
| Criterion TGX Stain-Free Precast Gels | Bio-Rad | cat: 5678034 |
| Intercept® T20 (TBS) Antibody Diluent | Li-Cor | cat: 927-65001 |
| Pierce™ BCA Protein Assay Kit | ThermoFisher | cat: 23225 |

| | | |
|---|---|---|
| Trans-Blot Turbo RTA Midi 0.2 µm Nitrocellulose Transfer Kit | Bio-Rad | cat: 1704271 |
| oYo-Link® Oligo Custom | AlphaThera | cat: AT1002-25ss |
| Invitrogen™ Dynabeads™ MyOne™ Streptavidin T1) | ThermoFisher | cat: 65601 |
| Biotin-SP (long spacer) ChromPure Human IgG, Fc fragment | Jackson ImmunoResearch | cat: 009-060-008 |
| NEBuffer™ 4 | New England BioLabs | cat: B7004S |
| **Animal Models** | | |
| Mice: C57BL/6J | Jackson Laboratory | cat: 000664 |
| Mice: Adipoq-TdTomato | Jackson Laboratory | cat: 010803 x cat: 007909 |
| Mice: Il17a-GFP | Esplugues et al., 2011 | |
| **qPCR Oligonucleotides** | | |
| Actb – forward primer (5'-3') | Millipore Sigma | GGCTGTATTCCCCTCCATCG |
| Actb – reverse primer (5'-3') | Millipore Sigma | CCAGTTGGTAACAATGCCATGT |
| Il17a – forward primer (5'-3') | Millipore Sigma | CAGACTACCTCAACCGTTCCAC |
| Il17a – reverse primer (5'-3') | Millipore Sigma | TCCAGCTTTCCCTCCGCATTGA |
| Rorc – forward primer (5'-3') | Millipore Sigma | GTGGAGTTTGCCAAGCGGCTTT |
| Rorc – reverse primer (5'-3') | Millipore Sigma | CCTGCACATTCTGACTAGGACG |
| **Software** | | |
| CellRanger - CLI | 10X Genomics | https://www.10xgenomics.com |
| Seqtk - CLI | GitHub | https://github.com/lh3/seqtk |
| velocyto - CLI | GitHub | https://github.com/velocyto-team/velocyto.py |
| Scanpy - python | GitHub | https://github.com/scverse/scanpy |
| scAR - python | GitHub | https://github.com/Novartis/scar |
| scVI-tools - python | GitHub | https://github.com/scverse/scvi-tools |
| Solo - python | GitHub | https://github.com/calico/solo |
| GSEApy - python | GitHub | https://github.com/zqfang/GSEApy |
| pySCENIC - python | GitHub | https://github.com/aertslab/pySCENIC |
| scVelo - python | GitHub | https://github.com/theislab/scvelo |
| velovi - python | GitHub | https://github.com/YosefLab/velovi |
| SCRAN - R | GitHub | https://github.com/MarioniLab/scran |
| Correlation - R | GitHub | https://github.com/easystats/correlation |
| GenomicFeatures - R | GitHub | https://github.com/Bioconductor/GenomicFeatures |
| Seurat - R | GitHub | https://github.com/satijalab/seurat |
| MAST - R | GitHub | https://github.com/RGLab/MAST |
| ggupset - R | GitHub | https://github.com/const-ae/ggupset |
| NicheNet - R | GitHub | https://github.com/saeyslab/nichenetr |
| rstatix - R | GitHub | https://github.com/kassambara/rstatix |
| ggplot2 - R | GitHub | https://github.com/tidyverse/ggplot2 |
| FloJo | BD Biosciences | https://www.flowjo.com |
| Prism | GraphPad | https://www.graphpad.com |

### RESOURCE AVAILABILITY

#### Lead Contact

#### Materials Availability

All reagents are commercially available.

#### Data and Code Availability

GEO accession numbers for raw sequencing data and code for reproducing our analysis will be made available at the time of publication. Raw sequencing data from Katzenelenbogen *et al.*, 2020 is publicly available (GEO GSE150877).

### EXPERIMENTAL MODELS AND SUBJECT DETAILS

#### Mice

Wild-type (WT) mice (8-12 weeks, males, C57BL/6J) were purchased from Jackson Laboratory (Bar Harbor, ME). Adipoq-Cre mice (Jackson Laboratory; stock no. 010803) were crossed with Ai9 mice (Rosa-CAG-LSL-tdTomato-WPRE; Jackson Laboratory; stock no. 007909) to create homozygous Adipoq-TdTomato bone marrow stroma reporter mice, as previously described (Wolock et al., 2019). Il17a-eGFP mice (IL17A-IRES-eGFP) were described previously (Esplugues et al., 2011). All mice were housed at the University of Alabama at Birmingham (UAB). Mice were provided with food and water *ad libitum* and housed under a 12-hour light-dark cycle. All experimental procedures were approved by the UAB Institutional Animal Care and Use Committee (IACUC).

#### Th17 T Cell Culture and Stimulation

Naïve CD4 T cells were isolated as previously described (Harbour et al., 2020). Briefly, mice were sacrificed, and spleens were dissected and mashed over a 70µm filter in complete IMDM media (IMDM media containing 10% FBS, 100 IU/ml penicillin, 100 µg/ml streptomycin, 1mM sodium pyruvate, 1x non-essential amino acids, 50µM β-mercaptoethanol, 1mM HEPES buffer, and 2mM glutamine). Red blood cells were removed using ACK lysis buffer, and naïve CD4 T cells were purified using the MACS Miltenyi negative CD4 T cell selection kit. To generate Th17 cells, naïve CD4 T cells were cultured for 72h at 37°C in complete IMDM media containing plate-bound anti-CD3 (10µg/ml; clone 145-2C11), anti-CD28 (1µg/ml; clone 37.51), IL-6 (20ng/ml), recombinant human TGF-β (2.5ng/ml), neutralizing anti-IFN-γ (10µg/ml; clone XMG1.2), and neutralizing anti-IL-4 (10µg/ml; clone 11B11). After 72h, Th17 cells were restimulated with phorbol myristate acetate (PMA; 50ng/ml), ionomycin (750ng/ml) and BD GolgiStop™ for 3 hours at 37°C.

#### Human Retina

Whole eye globes were obtained from Advancing Sight Network (Birmingham, AL). The donor was an 87-year-old male without a history of ocular disease. *Ex vivo* fundus and optical coherence tomography imaging was unremarkable (data not shown). Death to preservation time was 5.2 hours. Portions of the peripheral retina were stored in either DES at -80°C or fixed overnight at 4°C in 4% paraformaldehyde (PFA).

### METHOD DETAILS

#### DES Fixation

Deep eutectic solvent (DES) fixation was performed using vivoPHIX™ (Rapid Labs Ltd., UK).

For mouse bone marrow fixation, a clean transverse cut was made using a razor blade in the mid-diaphysis of the femur shaft. The bone marrow was then centrifuged directly into DES, similar to a recent report describing the preservation of bone marrow with RNAlater™ (Pedersen et al., 2019). Specifically, the very end of a 0.6mL microcentrifuge tube was cutoff using a razor blade and placed inside a 1.5mL microcentrifuge tube that contained ~100µL of DES. One half of the femur was then placed in the 0.6mL tube, with the exposed bone marrow pointing downward toward the DES (**Fig. 1B**). The two stacked tubes were then centrifuged at 5,700G for 30 seconds. The upper chamber containing the 0.6mL tube and the empty femur shaft was removed, and an additional ~200µL of DES was added on top. A clean 0.1-10µL pipette tip was then used to stir the bone marrow into the DES. The bone marrow-DES mixture was then incubated at room temperature for 2 hours before proceeding with ultrasonic dissociation.

For Th17 T cell fixation, culture media containing cells was transferred to 5mL FACS tubes and centrifuged at 4°C for 5 minutes at 300G. The supernatant was quickly decanted, and the cell pellet was loosened with light vortexing. ~300µL of DES was added directly onto the cells in FACS tubes, followed immediately by vortexing to mix the cells into the DES. The DES-Th17 mixture was then incubated at room temperature for 2 hours, after which aliquots were moved to 1.5mL microcentrifuge tubes and placed at -80°C for long-term storage.

For human retina fixation, after dissection and removal of the anterior chamber, a 10mm corneal trephine was used to isolate a punch biopsy of the peripheral retina. Excess water from the retina punch was gently wicked away, and the retina was then submerged in ~300µL of DES. After 2-3 hours at room temperature, the DES-retina mixture was moved to -80°C for long-term storage.

For mouse colon fixation, the transverse colon was acutely dissected, and feces were removed by washing the luminal contents out with 1x phosphate buffered saline (PBS). The lumen was then filleted open with a single cut and a ~10mm long portion was obtained. Excess water was gently wicked away, and the colon was then submerged in ~500µL of DES. After 2-3 hours at room temperature, the DES-colon mixture was moved to 4°C for short-term storage prior to ultrasonic dissociation.

#### Dissociation of DES-Fixed Tissues

Dissociation was performed in a 28/40khz dual-frequency ultrasonication water bath (Vevor, Walmart; UPC 650971639459). DES-fixed solid tissues (i.e. human retina and mouse colon) were removed from the DES, padded dry using a Kimwipe, and then minced into small (~1mm) pieces in ~100µL of fresh DES using fine dissecting scissors. For bone marrow, any remaining large bone marrow plugs were minced directly in the original DES using fine dissecting scissors. The DES was then diluted to 90% DES/10% $H_2O$, either by adding a small volume of nuclease-free water containing 8mM EDTA directly to the DES (in the case of mouse bone marrow), or by premixing DES with nuclease-free water containing 8mM EDTA and then transferring the minced tissue directly into it (in the case of mouse colon and human retina). The final volume of the tissue plus diluted DES was ~200-300µL. The mixture was transferred to 1mL thick-wall polycarbonate ultracentrifuge tubes, and the open ends were covered using pieces of transparent adhesive film. The tissue plus diluted DES was vortexed at max speed for ~15-30 seconds to ensure complete mixing before being suspended in the ultrasonication water bath, which was prechilled to 4°C using ice. The samples were sonicated for 10 minutes (5 minutes for retina), with pauses every 2-3 minutes for a brief (~10 seconds) vortexing at max speed. Dissociation can be monitored visually through increases in turbidity. Typically, some small clumps of undissociated tissue will remain after 10 minutes of sonication and will be filtered out upon transfer to the aqueous buffer.

### Transfer from DES to Aqueous Buffer, Blocking, and Immunostaining

For DES-fixed cells and dissociated solid tissues, transfer from DES to aqueous buffer occurred following treatment with 10% glacial acetic acid (AcOH), which we found was necessary to prevent RNase reactivation. If the samples were stored at -20°C or -80°C, the specimens were first allowed to acclimate to room temperature for 10-15 minutes. First, dissociated samples/cells were transferred to 5mL FACS tubes, and the volume of each sample was recorded. An equal volume of 20% AcOH/80% DES was then premixed, using vigorous vortexing to ensure complete mixing. Equal volumes of the 20% AcOH/80% DES mixture were then added to the FACS tubes containing the DES-fixed samples, and the tubes were vortexed at max speed for ~10-15 seconds to ensure complete mixing. The samples were then incubated at room temperature for 5 minutes, and then ~4mL of ice-cold rehydration buffer (3X saturated sodium chloride (SSC) containing 8mM EDTA) was added to the samples. The FACS tubes were capped and inverted several times to ensure the complete dissolving of DES. The samples were then filtered through 40-70µm strainers into new FACS tubes, which were immediately centrifuged at 300G for 5 minutes in a centrifuge prechilled to 4°C. Following centrifugation, the supernatant was decanted, and any remaining large beads of solution along the inner walls of the FACS tubes were removed. The samples were then resuspended in staining buffer (~200-500µL depending on the experiment) and incubated on ice for ~15-30 minutes to allow the blocking of non-specific epitopes. Staining buffer contained 1% BSA, 100U/mL heparin sodium salt, 4mM EDTA, 5% True-Stain Monocyte Blocker™, 5% TruStain FcX™ PLUS, 5% TruStain FcX™, and 1x RNasin® Plus in 1x SSC. Following blocking, primary antibodies diluted in staining buffer were added to the FACS tubes, and samples were stained on ice for 30 minutes. Following staining, samples were diluted with 4mL of ice-cold wash buffer (1% BSA, 4mM EDTA in 1x SSC) and centrifuged at 300G for 5 minutes at 4°C to remove unbound antibodies. For flow cytometry, samples were resuspended in wash buffer, whereas for scRNA-seq, samples were resuspended in ice-cold 1x SSC containing 1x RNasin® Plus and 0.04% BSA. Note that for phospho-staining, 1x Halt™ Phosphatase Inhibitor Cocktail was added to the rehydration buffer, and 2x Halt™ Phosphatase Inhibitor Cocktail was added to the staining buffer. Wash buffer and final resuspension buffers did not contain phosphatase inhibitors; we found that primary antibodies protected phospho-epitopes from dephosphorylation, as has been previously reported (Krutzik and Nolan, 2003). Also note that RNasin® Plus was only included in the staining buffer for experiments involving sequencing or RNA analyses. Lastly, for inCITE-seq, in addition to 2x Halt™ Phosphatase Inhibitor Cocktail, 1mg/mL of a 30-mer blocking oligonucleotide was added to the staining buffer. The 30-mer oligonucleotide (5'-CTAGACTGATTACGTACGTAAGATCGCTAC-3') was designed to lack complementarity to any known endogenous mouse or human DNA sequences and contained a dideoxycytosine at the 3' end to prevent extension by polymerases.

### Bone Marrow RNA Electrophoresis

We optimized the transition from DES to aqueous buffer using RNA electrophoresis on ultrasonically dissociated bone marrow samples. We tested 3 aqueous transfer strategies: (1) treatment with 10% AcOH followed by excess 3x SSC, (2) excess 3x SSC only, as previously reported for methanol-fixed cells (Chen et al., 2018), and (3) methanol treatment followed by resuspension in a high-salt buffer akin to diluted RNAlater (4M ammonium sulfate), as recently reported for methanol-fixed cells (Amit *et al.*, 2020; Katzenelenbogen *et al.*, 2020). The same bone marrow DES samples were split 3 ways to test each of the 3 methods in parallel. For the AcOH method, samples were treated with 10% AcOH for 5 minutes at room temperature, followed by washing with 4mL of ice-cold 3x SSC containing 8mM EDTA, as described above. After centrifugation, the solution was decanted, the cell pellet was loosened with light vortexing, and cells were lysed using Qiagen Buffer RLT, according to the manufacturer's instructions. For the 3x SSX only method, the protocol was identical to that described using the AcOH method but without the 10% AcOH treatment step. For the methanol/high-salt buffer method, we adapted a recently described protocol used in methanol fixed cells (Amit *et al.*, 2020; Katzenelenbogen *et al.*, 2020). Briefly, in a 1.5mL microcentrifuge tube containing DES-dissociated bone marrow, 4 volumes of ice-cold methanol were added dropwise while gently vortexing the sample to ensure mixing. The mixture was then centrifuged at 300G for 3 minutes at 4°C. The methanol was removed and 500µL of ice-cold 3x SSC was gently added without disturbing the cell pellet. The 3x SSC was removed, and the cells were resuspended in 100µL of ice-cold high-salt buffer (3.4M ammonium sulfate, 50mM EDTA, 1x RNasin® Plus in nuclease-free water, pH 5.2) and incubated on ice for 10 minutes. The cell mixture in high-salt buffer was then transferred to a FACS tube, 3mL of ice-cold 3x SSC was added, and the sample was centrifuged at 300G for 5 minutes at 4°C. After centrifugation, the solution was decanted, the cell pellet was loosened with light vortexing, and cells were lysed using Qiagen Buffer RLT, according to the manufacturer's instructions. RNA purification and DNase treatment were performed using the Qiagen RNeasy Plus Mini kit according to the manufacturer's instructions. The quality of purified RNA was then visualized with electrophoresis using a 1% bleach gel, prestained with ethidium bromide, as previously described (Aranda et al., 2012).

**RT-qPCR, RNA Integrity, and RNA Quantity in Th17 T Cells**

Th17 T cells were differentiated in culture for 3 days as described above. Th17 cells used for RT-qPCR, RNA integrity number analysis, and RNA yield experiments did not receive secondary stimulation with PMA/ionomycin. DES fixation was performed as described above, and aliquots of unfixed viable cells were used for comparison. After 2 hours of DES fixation at room temperature, aliquots of DES fixed cultures were stored at -20°C or -80°C for 1 month. DES fixed cells were treated with 10% AcOH and transferred to aqueous buffer, as described above. DES fixed cells were stained with 0.01mM SYTO40 in staining buffer on ice for 15 minutes. Viable cells were stained with viability dye on ice for 15 minutes. 100,000 DNA+ cells (or 100,000 live cells for viable samples) were then FACS sorted directly into 500uL of Trizol per sample. RNA purification and DNase treatment were performed using the Direct-zol Miniprep kit according to the manufacturer's instructions. cDNA was generated using the Superscript IV VILO master mix kit according to the manufacturer's instructions. qPCR was performed in 20µL reactions using SsoAdvanced Universal SYBR Green Supermix and a Bio-Rad CFX96 thermocycler. RNA input was normalized using two independent methods: normalized RNA mass per sample (determined using the Quant-it™ RiboGreen RNA Assay Kit) or total RNA yield per sample (i.e. absolute recovery from 100,000 cells). Raw Ct values were used for analysis. RIN analysis was performed using an Agilent 2100 per the manufacturer's instructions. RNA quantity analysis was done using 2 methods, Agilent 2100 and a fluorescence-based method using the Quant-it™ RiboGreen RNA Assay Kit. Both were performed according to the manufacturers' instructions.

**Imaging Flow Cytometry**

Imaging flow cytometry was performed using an Amnis ImageStream MKII. Briefly, DES-fixed mouse bone marrow, human retina, and mouse colon were dissociated in DES as described above. Following dissociation, DES-fixed samples were transferred to FACS tubes and an equal volume of 8% PFA was added directly to the DES followed by gentle stirring with a pipette tip. Additional fixation was done to reinforce delicate structures that can be sheared upon transfer to aqueous buffer, especially when multiple rounds of centrifugation are required. The cells were allowed to fix on ice for 10 minutes in 4% PFA, after which 4mL of ice cold 3x SSC was added. The cell suspensions were then filtered using 5µm track etched hydrophilic membranes under light suction. The cells were then gently washed off of the membranes into FACS tubes by placing the membranes directly into ~500µL of staining buffer. 0.01mM SYTO40 was then added to the FACS tubes, and the cells were stained for 15 minutes on ice. After nuclear staining, the samples were again washed and filtered as described above before being resuspended in ~100µL of wash buffer. The cells were then loaded on the Amnis ImageStream and brightfield, side scatter, and DAPI channels were used for capturing images (the PE channel was also used for the mouse bone marrow samples to capture Adipoq-TdTomato staining). IDEAS (version 6.2) software was used for quality control filtering, image optimization, and exporting captured images. Photoshop was used to crop selected images for creating the collages in Figures 1 and 4.

### Confocal Immunofluorescence

Confocal immunofluorescence was performed on PFA-fixed mouse bone marrow, mouse colon, and human retina. For mouse bone marrow, femurs were dissected from a 13-week-old female Adipoq-Cre/Td-Tomato mouse and fixed overnight in 4% PFA at 4°C. The bones were washed in 1x PBS and then decalcified in 50mL of 10% EDTA (in 1x PBS, pH 7.5) at 4°C for 1 week on a rocker. The bones were then padded dry and cryoprotected in sequential sucrose gradients at 4°C (15% sucrose overnight, then 30% sucrose overnight). The bones were flash frozen in O.C.T. compound, and 50µm cryosections were cut. The slides were dried at 37°C for 2 hours and then rehydrated in 1x PBS, followed by blocking/permeabilization with Trident Universal Protein Blocking Reagent and 5% TruStain FcX™ PLUS at room temperature for 1 hour. Biotinylated anti-LEPR (final concentration 0.04 mg/mL) and FITC conjugated anti-CD45 (1:25) were diluted in Trident Universal Protein Blocking Reagent and incubated overnight at 4°C. The slides were washed in excess 1x PBS and then incubated with Streptavidin-Alexa Fluor 647 in 1% BSA and 1% horse serum in 1x PBS at room temperature for 1 hour. The slides were again washed in 1x PBS followed by Hoechst staining for 5 minutes at room temperature. After one final wash in 1x PBS, the slides were mounted with ProLong Gold Antifade mountant, pressed flat for 10 minutes, and then imaged on a Nikon AX confocal.

For the mouse colon, large intestine was dissected from an 8-week-old WT mouse, flushed of luminal contents, and fixed overnight in 4% PFA at 4°C. Samples were then embedded in paraffin and 5µm thick sections were cut. For staining, the slides were first incubated for 2 hours at 60°C. Deparaffinization and rehydration was performed with three sequential 5 minute incubations in xylene, two sequential 5 minute incubations in 100% ethanol, two sequential 5 minute incubations in 95% ethanol, and washed in distilled water in three sequential 5 minute incubations with gentle agitation. Citrate buffer prewarmed to 70°C was used for antigen retrieval and was incubated in a heated steamer for 20 minutes, followed by three washes in distilled water with gentle agitation. Sections were incubated with 1x PBS for 10 minutes and were then blocked with Trident Universal Protein Blocking Reagent and 5% TruStain FcX™ PLUS at room temperature for 1 hour. Slides were stained overnight at 4°C with a rabbit primary antibody to E-Cadherin (1:100) and fluorescein-conjugated UEA-1 Lectin (1:200). The slides were washed in excess 1x PBS and then incubated with anti-rabbit Alexa Fluor plus 647 in 1% BSA and 1% horse serum in 1x PBS at room temperature for 1 hour. The slides were again washed in 1x PBS followed by Hoechst staining for 5 minutes at room temperature. After one final wash in 1x PBS, the slides were mounted with ProLong Gold Antifade mountant, pressed flat for 10 minutes, and then imaged on a Nikon AX confocal.

For the human retina, pieces of peripheral retina were dissected as described above and fixed overnight in 4% PFA. Retinas were then washed with 1x PBS and blocked/permeabilized with Trident Universal Protein Blocking Reagent and 5% anti-human TruStain FcX™ at room temperature for 1 hour. Retina pieces were then stained overnight at 4°C with either biotinylated peanut agglutinin (PNA lectin; 1:100) and Alexa Fluor 647 conjugated anti-rhodopsin (1:50), or Alexa Fluor 647 conjugated anti-Iba1 (1:100). Retina pieces were then washed with 1x PBS followed by Hoechst staining for 5 minutes at room temperature. After one final wash in 1x PBS, the slides were mounted with ProLong Gold Antifade mountant, pressed flat for 10 minutes, and then imaged on a Nikon AX confocal.

### Bone Marrow Single-Cell RNA-Sequencing

Bone marrow single-cell RNA-sequencing (scRNA-seq) was performed on two 8-week-old male WT mice. The mice were euthanized, and femurs were dissected. A clean transverse cut was made using a razor blade in the mid-diaphysis of both femur shafts. The proximal half of one femur and the distal half of the other femur were then used to centrifuge the bone marrow directly into DES, as described above. Additional DES was added to the tubes immediately after centrifugation, a clean pipette tip was used to mix the bone marrow into the DES, and then the samples were incubated at room temperature for 2 hours.

The remaining 2 femur halves were used for viable samples. Using a similar bone marrow centrifugation strategy as DES (i.e. placing a 0.6mL microcentrifuge tube with the end cut off inside of a 1.5mL microcentrifuge tube), bone marrow was briefly centrifuged out of the femur shaft into ~100µL of Ficoll (5,700G for 30 seconds at 4°C). For one of the viable ½ femurs, 1mL of ice-cold FACS buffer (2% FBS, 2mM EDTA in 1x PBS) was immediately added on top of the Ficoll, and the sample was mechanically dissociated via pipetting with a 1000µL pipette until the bone marrow plug was completely disaggregated. The sample was then filtered through a 70µm strainer into a FACS tube and 3mL of additional ice-cold FACS buffer was added on top. For the remaining ½ femur, the end of a 1mL pipette tip was cut off with a razor blade and the bone marrow plug was transferred to a glass flat bottom 5mL conical containing a micro-magnetic stir bar and 1mL of dissociation buffer prewarmed to 37°C (minimum essential media (MEM) with 10% FBS, 1x pyruvate, 1x GlutaMAX, 1x MEM Non-Essential Amino Acids Solution, 25mM HEPES, 1% collagenase D, 0.5% dispase II, 0.5x Tryple Express, and 10U/mL DNase I). Care was taken not to disturb the bone marrow plug during the transfer from Ficoll to dissociation buffer. The glass dissociation vessel was then placed in a 37°C water bath, and the bone marrow was dissociated with gentle magnetic stirring for ~3 minutes or until the bone marrow plug was completely disaggregated. After 3 minutes, the glass dissociation vessel was immediately moved to ice and ~3.5mL of ice-cold FACS buffer was added. The cell suspension was then filtered through a 70µm strainer into a FACS tube. Both viable samples were then centrifuged at 300G for 5 minutes at 4°C in a prechilled centrifuge. The supernatant was decanted, and the cell pellets were resuspended in 0.5mL FACS buffer on ice. The enzymatically dissociated bone marrow was used for FACS sorting the non-hematopoietic stroma enriched fraction (CD45- B220- Ter-119-). Conversely, the mechanically dissociated viable bone marrow was used for FACS sorting CD45+/B220+ Ter-119- hematopoietic cells; we did this to eliminate any potential negative effects from the dissociation enzymes on immune cells. Of note, validation studies prior to our bone marrow scRNA-seq experiment confirmed that rapid centrifugation of bone marrow into Ficoll followed by mechanical dissociation via pipetting was no more harmful to CD45+ Ter-119- hematopoietic cells than other standard methods. Compared to flushing bone marrow out of the femur shaft or crushing bones in a mortar (2 standard methods of bone marrow isolation), centrifugation of bone marrow onto Ficoll resulted in equivalent viability, equivalent RIN, and equivalent RNA yield per cell in 100,000 FACS sorted CD45+ Ter-119- cells (**Supp. Fig. 11**).

After 2 hours of fixation, DES bone marrow was dissociated in 90% DES/10% $H_2O$, as described previously. Following dissociation, equal volumes of 20% AcOH/80% DES were added, and the samples were incubated for 5 minutes at room temperature. After 5 minutes, excess 3x SSC was added and the DES samples were filtered, centrifuged, and resuspended in staining buffer, as described previously. The DES samples were stained on ice for 30 minutes with PE conjugated anti-CD45 (1:50), PE conjugated anti-CD45R (1:50), Alexa Fluor 647 conjugated anti-Spectrinβ (1:25), and SYTO40 (0.01mM). After staining, excess antibody was washed off, and the cells were resuspended in ~500uL of final resuspension buffer (ice-cold 1x SSC containing 1x RNasin® Plus and 0.04% BSA) for FACS sorting. 50,000 CD45+/B220+ Spectrinβ- cells and 50,000 CD45- B220- Spectrinβ- cells per animal were FACS-sorted into the same 1.5mL microcentrifuge tube containing 5µL of final resuspension buffer. These FACS-sorted DES cells were then counted and loaded directly into 10X Genomics 3' V3 wells (1 well per animal).

Viable samples were blocked with 5% True-Stain Monocyte Blocker™, 5% TruStain FcX™ PLUS, and 5% TruStain FcX™ in FACS buffer for 15 minutes on ice. After blocking, the following antibodies and viability dye were added: PE conjugated anti-CD45 (1:50), PE conjugated anti-CD45R (1:50), APC conjugated anti-Ter-119 (1:50), and LIVE/DEAD™ Fixable Near-IR Dead Cell Stain (1:1000). Samples were stained for 30 minutes on ice, washed, and resuspended in 1x PBS with 0.04% BSA for FACS sorting. 50,000 CD45+/B220+ Ter119- cells from the mechanically dissociated fraction and 50,000 CD45- B220- Ter119- cells from the enzymatically dissociated fraction were FACS sorted per animal into the same 1.5mL microcentrifuge tube containing 5µL of 1x PBS with 0.04% BSA. These sorted viable cells were then counted and loaded directly into 10X Genomics 3' V3 wells (1 well per animal).

**Immunoblotting for Bone Marrow Post-Translational Modifications**

As described in **Figure 3G**, we isolated femurs from individual mice (n=4 for ubiquitin experiment; n=5 for phosphorylation/acetylation experiment), cut the bones in half, and briefly centrifuged the bone marrow into 3 separate solutions: (1) DES, (2) lysis buffer, or (3) Ficoll. Bone marrow from the Ficoll group was immediately dissociated, via pipetting in FACS buffer, washed, and then stored on ice for 3 hours before being washed with 1x PBS and lysed in lysis buffer, thereby simulating our scRNA-seq experiment. The same lysis buffer was used for the 3 experimental groups (RIPA buffer with 2x Halt™ Protease and Phosphatase Inhibitor Cocktail). For the DES group, ~200µL of additional DES was added on top following centrifugation. However, for this experiment the cells were not mixed with a pipette but rather left as a thin layer, which was submerged in the DES for 2 hours at room temperature. After incubation, the thin layer of cells was removed from DES using forceps, padded dry with a kimwipe, and chopped into tiny fragments using a razorblade on a petri dish containing ~500µL of lysis buffer. All samples were allowed to incubate on ice for 1 hour in lysis buffer with periodic vortexing. After 1 hour, the samples were sonicated using a probe sonicator (3x10s pulses at 30% power on ice). The samples were then centrifuged at 10,000G for 10 minutes at 4°C, and the supernatant was collected and stored at -80°C. Protein was quantified using the Pierce™ BCA Protein Assay Kit. Protein was linearized in Laemmli buffer at 100°C for 10 minutes, and then 20µg per sample were loaded onto 18 well 10% Criterion™ TGX Stain-Free™ Protein Gels. Following electrophoresis, total protein stains were activated using a Bio-Rad ChemiDoc Imaging System per the manufacturer's instructions and then transferred onto 0.2µm nitrocellulose membranes using the Trans-Blot® Turbo™ Transfer System per the manufacturer's instructions. After transfer, images of total protein stains were obtained. The blots were then treated with Pierce™ SuperSignal™ Western Blot Enhancer per the manufacturer's instructions and then blocked for 1 hour at room temperature using Intercept® (TBS) Blocking Buffer. After washing with 1x TBS, blots were incubated overnight with gentle rocking at 4°C with the following primary antibodies: anti-ubiquitin (1:1000), anti-acetyl lysine (1:1000), and anti-phospho-threonine (1:1000). Antibodies were diluted in primary antibody buffer from the Pierce™ SuperSignal™ Western Blot Enhancer kit. The next day, blots were washed several times with 0.1% Tween-20 in 1x TBS and incubated for 1 hour with secondary antibodies diluted in Intercept® T20 (TBS) Antibody Diluent. Alexa Fluor Plus 800 secondary antibodies were used and diluted at 1:5000. The blots were then washed several times with 0.1% Tween-20 in 1x TBS and then 2 washes in 1x TBS without Tween-20 before being imaged on a Bio-Rad ChemiDoc Imaging System.

**Flow Cytometry for LEPR+ Stroma Viability**

Femurs from individual Adipoq-TdTomato mice were dissected and bone marrow was isolated using 3 different methods (**Fig. 4C**). One femur was crushed with a mortar and pestle, and the isolated bone marrow was further dissociated into a single-cell suspension via pipetting. The remaining femur was cut in half and the bone marrow was isolated by rapid centrifugation into Ficoll, followed by either brief enzymatic dissociation (described in further detail above) or mechanical dissociation in FACS buffer via pipetting (described in further detail above). Samples were then blocked with 5% True-Stain Monocyte Blocker™, 5% TruStain FcX™ PLUS, and 5% TruStain FcX™ in FACS buffer for 15 minutes on ice. Following blocking, the following antibodies and viability dye were added: APC conjugated anti-CD45 (1:50), APC conjugated anti-CD45R (1:50), Brilliant Violet 605 conjugated anti-Ter-119 (1:50), and LIVE/DEAD™ Fixable Near-IR Dead Cell Stain (1:1000). Samples were quantified using a BD FACSymphony™ A5 and the viability of TdTomato+ CD45- CD45R- Ter119- stromal cells were compared.

### Th17 Flow Cytometry

Flow cytometry was performed on Th17 T cells for IL17A protein and GFP from Il17a-IRES-eGFP mice. After 3 days of differentiation, Th17 cultures (n=3) were split and half were given a secondary stimulation with PMA/ionomycin and BD GolgiStop™ for 3 hours, as described above. These control and stimulated cultures were then split a final time, with half using DES for fixation/permeabilization and the other half with PFA/methanol using the BD Cytofix/Cytoperm Kit. Specifically, the cultures were split into 2 equal volumes using FACS tubes and centrifuged at 4°C in undiluted culture medium (300G for 5 minutes). Supernatant was discarded and cell pellets were loosened with light vortexing before adding either ~300µL of DES or ~300µL of ice-cold Cytofix/Cytoperm. DES samples fixed for 2 hours at room temperature while PFA/methanol samples fixed for 30 minutes on ice before being exchanged with BD Perm/Wash™ Buffer. After 2 hours of fixation, DES samples were treated with AcOH, moved through 3x SSC to 1x SSC, and incubated in staining buffer for 15 minutes, as described in detail above. Both DES and PFA/methanol cells were then stained with APC anti-IL17A (1:100) and unconjugated chicken anti-GFP (1:200) for 30 minutes on ice. Cells were washed with their respective wash buffers and then stained with Alexa Fluor Plus 488 anti-chicken secondary antibody (1:750) and SYTO40 (0.01mM) for 30 minutes on ice. Samples were washed a final time and then resuspended in either FACS buffer for PFA/methanol or 1% BSA, 4mM EDTA in 1x SSC for DES cells. For both fixation types, APC conjugated rat isotype controls were included as well as controls without primary antibody for the AF+488 secondary stain (**Supp. Fig. 7**). Samples were quantified using a BD FACSymphony™ A5.

Flow cytometry for phospho-epitopes was performed on DES-fixed Th17 T cells (n=3 cultures) that were either left in basal culture medium (unstimulated) or treated with PMA/ionomycin for 3 hours (stimulated), as described previously. After 2 hours of fixation in DES at room temperature, samples were stored at -80°C for several days. Samples were allowed to warm to room temperature for ~15 minutes after which aliquots of each were moved to FACS tubes. Samples were treated with AcOH and moved through 3x SSC to staining buffer, as described previously but with the addition of 1x Halt™ Phosphatase Inhibitor Cocktail in the rehydration buffer and 2x Halt™ Phosphatase Inhibitor Cocktail in the staining buffer. After blocking for 15 minutes, samples from each culture and condition were then counted and diluted to the same concentration of cells/µL. Samples were then split 15 ways in equal volumes and stained with 1 of 13 different unconjugated anti-phospho-epitope antibodies (**Fig. 5H**), 12 of which were rabbit clones and the remaining was a mouse clone (See resources table; each used at 1:25 dilution). Isotype controls for both rabbit and mouse were included. Cells were washed with wash buffer (1% BSA, 4mM EDTA in 1x SSC) and then stained with Alexa Fluor Plus 647 anti-rabbit or anti-mouse secondary antibodies (1:750) and SYTO40 (0.01mM) for 30 minutes on ice. Samples were washed a final time and then resuspended in wash buffer. In addition, an aliquot of each culture was stained with PE conjugated anti-RORγt and corresponding isotype controls were included. Samples were quantified using a BD FACSymphony™ A5.

Flow cytometry for IL2 production was performed on PFA/methanol fixed Th17 T cells (n=5 cultures) that were treated with PMA/ionomycin for 3 hours, as described previously. After 3 hours, cultures were moved to FACS tubes and centrifuged at 4°C in undiluted culture medium (300G for 5 minutes). Supernatant was discarded and cell pellets were loosened with light vortexing before adding ~300µL of ice-cold Cytofix/Cytoperm. Samples were incubated on ice for 1 hour before being exchanged with BD Perm/Wash™ Buffer containing 1x Halt™ Phosphatase Inhibitor Cocktail and 0.1% Triton-X100 for nuclear permeabilization. After 30 minutes of permeabilization on ice, samples were exchanged with BD Perm/Wash™ Buffer containing only 1x Halt™ Phosphatase Inhibitor Cocktail and then stained with unconjugated rabbit anti-p-ERK1/2 (1:33 dilution) for 30 minutes on ice. After washing off the staining buffer, samples were resuspended in BD Perm/Wash™ Buffer containing 1x Halt™ Phosphatase Inhibitor Cocktail and anti-rabbit Alexa Fluor Plus 488 secondary antibody (1:500 dilution) for 20 minutes on ice. After washing off the secondary antibody, samples were again resuspended in BD Perm/Wash™ Buffer containing 1x Halt™ Phosphatase Inhibitor Cocktail and then stained with PE-conjugated anti-IL2

(1:100 dilution), AF647-conjugated p-FOS (1:50 dilution), and SYTO40 (0.01mM) for 30 minutes on ice. Samples were washed a final time and then resuspended in FACS buffer and quantified using a BD FACSymphony™ A5. AF647-conjugated rabbit isotype, unconjugated rabbit isotype, and PE-conjugated rat isotype controls were included (**Supp. Fig. 10**).

#### Oligonucleotide-Antibody Conjugation for Intracellular CITE-Sequencing

Custom oligonucleotide-antibody conjugates were fabricated for intracellular CITE-seq (inCITE-seq). Custom 80 base-pair (bp) oligonucleotides, based on a modified TotalSeq™-B design, were produced by IDT. The oligos had a 34bp 5' PCR handle followed by 7 random spacer nucleotides (designated with N's): 5'- GTGACTGGAGTTCAGACGTGTGCTCTTCCGATCTNNNNNNN -3'. A 22bp 3' capture sequence was used, which was preceded by 6 random spacer nucleotides: 5'-NNNNNNGCTTTAAGGCCGGTCCTAGCAA -3'. The final 2 adenine bases at the 3' end of the capture sequence used phosphorothioated bonds. Finally, between the 2 sets of random spacer nucleotides was an 11bp unique barcode. 3 of these 80bp custom oligonucleotides were obtained with their own unique 11bp barcodes (barcode1: GTCACTACGAG, barcode2: TGGCTACAAGT, and barcode3: CGACATTGACA) and were conjugated at the 5' end to a proprietary Protein G derivative with inducible cross-linking moieties upon UV light activation (oYo-Link®; AlphaThera Inc., Philadelphia, PA). This product results in high-efficiency and site-specific conjugation to the Fc region of a broad diversity of IgG isotypes from many different source species; 2 oYo-links bind per antibody molecule.

The procedure for conjugation of inCITE-seq antibodies was performed as described below. The following antibodies (Ab) were conjugated: p-STAT3 Y705 (3µg Ab; clone 4/P-STAT3; barcode1), p-ERK1/2 T202/Y204 (1.5µg Ab; clone 197G2; barcode1), p-p65 S536 (1µg Ab; clone 93H1; barcode2), p-FOS S32 (0.5µg Ab; clone D82C12; barcode3), and rabbit isotype (3µg Ab; clone DA1E; barcode3). The oligo-conjugated oYo-links were reconstituted to 33µM and added to the respective antibody aliquots at a molar ratio of 1 Ab : 5 oYo in clear 0.6mL microcentrifuge tubes. The Ab/oYo solution was gently pipetted up and down to mix and then incubated at room temperature for 10 minutes. After this, the Ab/oYo mixtures were placed on ice and exposed to UV light for 2 hours using the LED PX2 Photo-Crosslinking Device (AlphaThera Inc.).

After cross-linking, excess unbound oYo-link was removed using streptavidin-conjugated magnetic beads that were preincubated with biotinylated human IgG Fc fragments. Despite the small size of oYo-link (~8kDa), we found that even 100kDa ultracentrifugation filters were unable to remove unbound oligonucleotide-conjugated oYo-link. Therefore, we alternatively introduced excess Fc to bind unbound oYo-link. First, 560µL of beads (~62.5µL of beads per µg of antibody) were washed 3x with 1mL of 1x PBS containing 0.01% Tween-20 (PBST). The beads were then resuspended in 560µL of PBST and 67µL of IgG Fc (resuspended in nuclease-free water at 8.35mg/mL) was added to give a final concentration of 1µg Fc per 1µL beads. The 1.5mL microcentrifuge tube containing the mixtures were then incubated on a vortexer at low-speed for 30 minutes at room temperature to allow binding. After incubation, the bead mixture was then washed 3x with PBST and then 2x with 1x PBS, before being resuspended in 40µL of 1x PBS. For each 1µg of Ab used in the oYo-link conjugation, 4uL of bead mixture was added to the tube. The Ab/oYo-link/bead mixture was then incubated on a vortexer at low-speed for 1 hour at room temperature. 3 sequential magnetic separations were then performed by incubating the mixture on a magnetic rack for 5 minutes at room temperature and then transferring the supernatant to a new tube and repeating.

Lastly, the cleaned-up Ab-oligonucleotide conjugates were then incubated with E. coli ssDNA binding protein (EcoSSB) to prevent off-target oligonucleotide interactions, as recently reported (Chen *et al.*, 2022). To do this, we first calculated the number of moles of oYo-link added per antibody; this equated to ~$3.3 \times 10^{-11}$ mole per µL of oYo-link used (applicable when oYo-link is resuspended to 33µM, per the manufacturer's instructions). After calculating the number of moles of oYo-link used for each antibody, we then multiplied this number by 12 for the number of moles of EcoSSB to be used per Ab reaction, which was based on the notion that each tetramer of EcoSSB binds to ~35bp of ssDNA (Chen *et al.*, 2022). Based on the molecular weight of EcoSSB, this ultimately equated to adding ~7.5µg of EcoSSB per 1 µL of oYo-link used. This calculation purposely allowed excess EcoSSB, ensuring complete saturation of oligonucleotides. For each oligonucleotide-Ab conjugate mixture, 5µL of 10x NEBuffer4, the required volume of EcoSSB, and nuclease-free water were combined to give a final total volume of 50µL. The mixtures were then incubated at 37°C for 30 minutes in a thermocycler to allow EcoSSB binding. No further purification was used.

**Th17 T Cell Single-Cell RNA-Sequencing**

The same stimulated and unstimulated cultures (n=3) from the phospho-flow cytometry experiment described earlier were used for scRNA-seq. scRNA-seq was performed on 2 occasions using these samples: once on the day of fixation (designated: d0) and again after the samples had been stored at -80°C for 2 months (designated: d60). We will describe the d60 protocol first, which is where the inCITE-seq data in Figures 6 and 7 are derived from. Samples were allowed to warm to room temperature for ~15 minutes after which aliquots of each were moved to FACS tubes. Samples were then treated with 10% AcOH for 5 minutes at room temperature and then 4mL of ice-cold 3x SSC containing 8mM EDTA and 1x Halt™ Phosphatase Inhibitor Cocktail was added. Samples were filtered and centrifuged in a prechilled centrifuge at 300G for 5 minutes. The supernatant was decanted, and the samples were resuspended in 500µL of ice-cold staining buffer (1x SSC in nuclease-free water containing 1% BSA, 100U/mL heparin sodium salt, 4mM EDTA, 5% True-Stain Monocyte Blocker™, 5% TruStain FcX™ PLUS, 1x RNasin® Plus, 2x Halt™ Phosphatase Inhibitor Cocktail, and 1mg/mL of blocking oligonucleotide). Samples were allowed to block on ice for ~30 minutes during which the cell concentration of each sample was manually counted. The samples were then diluted to an equal concentration of cells/µL (1.5 million cells per mL) using the same staining buffer as above. Each sample was then split into even volumes and inCITE-seq was performed using two separate panels of antibodies, thereby generating 12 total unique samples (n=3 unstimulated cultures + n=3 stimulated cultures x 2 panels of inCITE-seq antibodies). All samples were hashed into the same well for 10X Genomics 3' V3 to control for any potential well-to-well variations in capture efficiency, sample handling, or sequencing. The following hashing scheme was used:

> All <u>unstimulated</u> culture samples were stained with TotalSeq™-B0301 anti-mouse Hashtag1 antibody (designated: Hashtag1) and all <u>stimulated</u> culture samples were stained with Hashtag2. In addition to these hashes, one of six different hashes was used to distinguish each culture and panel combination: cultures A, B, and C for panel1 were stained with Hashtags 3, 4, and 5, respectively while cultures A, B, and C for panel2 were stained with Hashtags 6, 7, and 8, respectively.

Panel1 for inCITE-seq was stained with p-STAT3 Y705 (barcode1), p-p65 S536 (barcode2), and rabbit isotype (barcode3). Panel2 was stained with p-ERK1/2 T202/Y204 (barcode1) and p-FOS S32 (barcode 3). After EcoSBB incubation, all antibodies were chilled on ice and antibodies for the aforementioned panels were combined. Staining buffer, as described above, was added to each panel to a final volume of 175µL and 0.02mM SYTO40 was added. Each panel was then split evenly into 2 parts (87.5µL each) and then 0.875µg (or 1.75µL) of either Hashtag1 or Hashtag 2 was added. Each panel/hashtag combination was then evenly divided 3 ways (28µL each) into clean, empty FACS tubes, for a total of 12. For each culture/panel combination, 0.6µL of the respective Hashtag was added to each tube. Finally, 175µL of each of the respective cultures was added to the designated tubes and the samples were stained on ice for 30 minutes. After 30 minutes, samples were washed 2x with 4mL of ice-cold wash buffer (1% BSA, 4mM EDTA in 1x SSC). Each sample was resuspended in 250µL of 1x SSC containing 0.04% BSA and 1x RNasin® Plus. 10,000 DNA+ cells from each of the 12 tubes were then FACS-sorted into the same tube. The final cell concentration was counted, and cells were loaded into one 10X Genomics 3' V3 well.

The d0 scRNA-seq using these same samples was from an earlier inCITE-seq optimization experiment. Samples from this experiment were treated identically to those of the d60 samples, with the following exceptions: EcoSBB was not used, magnetic removal of unbound oYo-link was not performed, samples were not blocked with 1mg/mL of blocking oligo, and separate 10X Genomics 3' V3 wells were used for unstimulated and stimulated cells. Culture A was stained with either Hashtag 1, 2, or 3. Culture B was stained with either Hashtag 4, 5, or 6. Culture C was stained with either Hashtag 7, 8, or 9.

### QUANTIFICATION AND STATISTICAL ANALYSES

#### Bone Marrow Single-Cell RNA-Sequencing

FASTQ files from bone marrow scRNA-seq samples were demultiplexed and aligned using CellRanger Count (Version 6.1.1). The "include-introns" flag was used. For alignment, we used the pre-built mouse reference genome from 10X Genomics, mm10 (GENCODE vM23/Ensembl 98). After initial CellRanger Count completion, the "metrics_summary" output file from each sample was used to calculate unique reads per cell. This number was normalized per sample for the proportion of mapped reads with valid barcodes (termed "Fraction of Reads in Cells" by 10X Genomics) and sequencing saturation, defined as "the fraction of reads originating from an already-observed UMI". The unique reads per cell calculation is defined by the following equation:

$$\text{reads per cell} = \frac{\left(\frac{100 - x}{100}\right)\left(\frac{y}{100}\right) z}{c}$$

where $x$ = percent sequencing saturation, $y$ = percent of reads in cells, $z$ = total number of reads, and $c$ = estimated number of cells.

Using this metric, 3 of the 4 samples were then downsampled to match the number of unique reads per cell in the lowest sample. Downsampling was performed using the Seqtk command line package (version 1.3) on raw FASTQ files. Downsampled FASTQs were then used to rerun CellRanger Count. Values from "metrics_summary" output files were again used to recalculate the above equation to verify correct downsampling.

After verification of downsampling, filtered and unfiltered CellRanger Count output files were then loaded into Scanpy (Wolf et al., 2018), and the scAR package (Sheng *et al.*, 2022) from scvi-tools (Gayoso et al., 2022a) was used to remove ambient RNA from each sample individually. For scAR, 0.995 was used for the "probability" parameter. Using the denoised matrix, we performed quality control filtering using a threshold of 500 for the minimum number of transcripts (i.e. UMIs) and the minimum number of genes per cell. Likewise, we used a cutoff of 0.1 for the proportion of mitochondrial reads per cell. 36,744 total cells were recovered after quality control filtering. The Solo package (Bernstein et al., 2020) from scvi-tools was then used to identify and remove doublets in each sample individually. The "soft" parameter for the predict() function was set to False, thereby returning discrete labels for predicted doublets.

The 4 samples were concatenated, and the single-cell Variational Inference (scVI) model from scvi-tools was used to compute a latent space representation of the combined samples. In constructing this particular model, we did not filter on highly variable genes, and we included 2 categorical covariates: sample-wise labels, for each of the 4 samples, and a group label for DES or viable. In addition, we included 5 continuous covariates: mitochondrial proportion, natural log of total counts per cell, natural log of total genes per cell, and "G2M" and "S" scores. The cell cycle scores were computed with the Scanpy function score_genes_cell_cycle() using the cell cycle gene list from (Tirosh et al., 2016). The resulting latent space was then used to compute nearest neighbors, followed by dimensionality reduction using the UMAP algorithm in Scanpy. In addition, we used the scVI function get_normalized_expression() with the "library_size" parameter set to "latent" to extract normalized expression values. Using the Leiden algorithm in Scanpy, we computed initial clustering (resolution=0.1). This initial clustering and normalized expression were then used to remove contaminating erythrocytes, which were relatively frequent in both DES samples. To do this, we removed the main erythrocyte cluster as well as any cells that had expression values above roughly the 87$^{th}$ percentile for the following 4 hemoglobin genes: Hbb-bs, Hbb-bt, Hba-a1, and Hba-a2. After removing the erythrocyte cluster, the predicted doublets via Solo, and the cells with hemoglobin expression above the 87$^{th}$ percentile, 27,555 cells remained.

After the removal of doublets and contaminating erythrocytes, we recomputed the scVI model to arrive at our final clustering. For this model, we subsetted the dataset on the top 1,000 highly variable genes using the Scanpy function highly_variable_genes() with the "flavor" parameter set to "Seurat_v3" and the "batch_key" parameter set to the sample identifier. For constructing the model we used the denoised raw count matrix, 1 categorical covariate (the sample identifier), and 4 continuous covariates (natural log of total counts per cell, natural log of total genes per cell, and "G2M" and "S" scores). The resulting latent space was then transferred to the non-subsetted dataset, which was subsequently used to compute nearest neighbors, followed by dimensionality reduction using the UMAP algorithm in Scanpy. Initial clustering was performed using the Leiden algorithm (resolution=0.06), which returned 7 clusters. Using well-defined bone marrow marker genes to guide subclustering, we then subclustered twice to arrive at our final clustering, which returned 12 unique cell populations. We then used the R package SCRAN (Lun et al., 2016) for cluster-wise size factor normalization of the denoised raw counts expression data. After dividing each expression value by the respective size factor, we then used natural log+1 transformation to complete normalization.

**Pearson Correlation Analyses**

For the Pearson correlation analyses in Figure 2, we downloaded raw FASTQ sequencing data from Katzenelenbogen *et al* 2020 (GSE150877). The data was from paired methanol-fixed and viable human PBMCs from 3 separate donors (6 samples in total), and the original experiment was performed using the 10X Genomics 5' v2 platform. The preprocessing of this dataset was essentially identical to that of the aforementioned mouse bone marrow scRNA-seq but with some dataset specific parameters. Briefly, CellRanger Count (Version 6.1.1) was used with the 10X Genomics pre-built human reference genome GRCh38 (GENCODE v32/Ensembl 98). Reads per cell, calculated with the aforementioned equation, were used in combination with Seqtk, to downsample to the highest common sequencing depth. Ambient RNA was removed using scAR. For quality control filtering, a threshold of 300 was used for minimum genes and minimum counts and a threshold of 0.2 was used for mitochondrial proportion. Solo was used to remove doublets. scVI modeling was used for computing dimensionality reduction and was performed using 4,000 highly variable genes. Categorical covariates were sample identifiers and fixation group identifiers. Continuous covariates were natural log genes per cell, natural log counts per cell, mitochondrial proportion, ribosomal proportion, and "G2M" and "S" scores. Leiden clustering identified 10 cell populations (**Supp. Fig. 4**). Finally, SCRAN was used for cluster-wise size factor normalization of the denoised raw counts expression data.

For performing gene-wise Pearson correlations, we restricted our analysis to clusters that contained at least ≥25 cells per sample. To begin, for each cluster in each sample, we calculated the mean expression value of each gene. For each cluster in the fixed or viable group, we then calculated the mean expression value using the individual sample means from the step before. For expression values at this stage, we used non-log transformed, size factor normalized counts. We then imported the "fixed" and "viable" mean expression tables into R and computed gene-wise Log2 fold-changes (Log2FC) by taking the Log2 transformation of normalized live expression (non-log transformed) divided by normalized fixed expression (non-log transformed). Genes that had 0 counts in either the live or fixed groups were eliminated. We then used the R package Correlation (Makowski et al., 2020) to compute gene-wise Pearson correlations for each cluster comparing Log2FC, transcript length, number of exons, G-C proportion, and expression magnitude. We used the Bonferroni method for p-value adjustments. For determining gene-wise characteristics in mice and humans, we used the canonical transcript from each gene. We exported G-C percent and transcript length (including UTRs and CDS) from BioMart. We used the "genes.gtf" file from each reference genome in combination with the R package GenomicFeatures to extract the number of exons per gene using the transcriptLengths() function. Expression magnitude was calculated with respect to the mean viable sample and was standardized on a 0 to 1 scale.

**Differential Gene Expression Analyses**

Differential gene expression (DGE) analyses were performed on natural log+1 transformed, size factor normalized expression data (SCRAN). The R package Seurat (Satija et al., 2015) was used for DGE computation using the MAST (Finak et al., 2015) implementation of the FindMarkers() function. The following parameters were used: "min.pct" was set to 0.01 to eliminate genes expressed in <1% of cells, "logfc.threshold" was set to 0 thereby not filtering genes based on fold change size, and "mean.fxn" was set to "rowMeans" thereby using group mean expression for Log fold-change calculations. In addition, for the bone marrow scRNA-seq "latent.vars" was set to natural log genes per cell and the sample identifier. For the Th17 inCITE-seq, "latent.vars" was set to natural log genes per cell.

**Intersection Analyses of Differentially Expressed Genes**

Output files from FindMarkers() were used for analyses of shared differentially expressed genes (DGs) between groups (clusters in Figure 3; phospho-targets in Figure 7). Gene names and direction of change for each group were combined for both intersection analyses. The R package ggupset was used for visualization.

**Pathway Enrichment Analyses**

For pathway analyses, the python program GSEApy (Fang et al., 2022) was used with the output data from FindMarkers(). Two different types of analyses were used. [1] Over-representation analyses (**Figure 3B**) were performed using the enrich() function with "background" genes set to all genes from the FindMarkers() output. Upregulated genes and downregulated genes were run separately. [2] Gene Set Enrichment Analyses (GSEA; **Figure 4J**) was done with the prerank() function using LogFC from FindMarkers() output to rank genes. "perumutation num" was set to 1000, "min_size" was set to 5, and "max_size" was set to 1000. The following gene sets were downloaded from gsea-msigdb and used for both analyses: Reactome, Hallmark, Wiki, Biocarta, Go-Biological, Go-Cellular, and Go-Molecular.

**Transcription Factor Activity Analyses**

The python package pySCENIC (Aibar *et al.*, 2017) was used for inferring transcription factor activity. Ranking databases and Motif to TF annotations were downloaded from the cistargetDBs website. Two ranking databases, both from reference genome mm10, were used: TSS±10kb and TSS±500bp/±100bp. The following 3 command-line functions were used in succession on the raw denoised matrix: "arboreto_with_multiprocessing.py", "pyscenic ctx", and 'pyscenic aucell". The output matrix was then loaded into Seurat, and the MAST implementation of FindMarkers() was used to compare conditions within groups; no "latent.vars" were used.

**Ligand-Receptor Analyses**

The R package NicheNet (Browaeys *et al.*, 2020) was used for ligand-receptor analyses. We performed a differential ligand-receptor analysis between viable and DES clusters, using script adapted from the NicheNet GitHub tutorial titled "Differential NicheNet analysis between conditions of interest". Databases for ligand-receptor priors were downloaded from the NicheNet GitHub page. Natural log+1 transformed, size factor normalized expression data (SCRAN) were used. The following prioritization weights were used: "scaled_ligand_score" = 5, "scaled_ligand_expression_scaled" = 1, "ligand_fraction" = 1, "scaled_ligand_score_spatial" = 0, "scaled_receptor_score" = 0.5, "scaled_receptor_expression_scaled" = 0.5, "receptor_fraction" = 1, "ligand_scaled_receptor_expression_fraction" = 1, "scaled_receptor_score_spatial" = 0, "scaled_activity" = 0, "scaled_activity_normalized" = 1, "bona_fide" = 1, "bona_fide" = 1. “expression_pct” was 0.1, “lfc_cutoff” was 0.1, “specificity_score_targets” was ‘min_lfc’, and “specificity_score_LR_pairs” was ‘min_lfc’. NicheNet was run twice, once for viable and once for DES, on each of the two clusters of interest (monocytes or neutrophils). The cluster of interest was set as the receiver and all other clusters were used as potential senders.

**RNA Velocity and Pseudotime Analyses**

The command line package velocyto (La Manno et al., 2018) was used for determining splicing counts in the bone marrow scRNA-seq dataset. RNA velocity analyses were then performed using the python packages scVelo (Bergen et al., 2020) and velovi (Gayoso et al., 2022b). We used scAR to denoise the spliced and unspliced expression matrices from velocyto. scAR requires a large number of empty droplets for accurate modeling, but the velocyto “run10x” function uses cell barcodes from the “filtered” CellRanger output. Therefore, we first calculated the 200,000 barcodes with the highest number of UMIs per sample and replaced the “barcodes.tsv.gz” file in the CellRanger filtered outputs directory with this list. We chose the top 200,000 because this was sufficient to run scAR using a “probability” of 0.995 and because velocyto was unable to scale up to the full barcode whitelist from 10X Genomics 3’ V3 (1,683,776 barcodes). We then performed scAR ambient RNA removal using the full 200,000 cell velocyto matrices as well as the filtered velocyto matrices, which were subsetted to include only the barcodes that CellRanger used in its original “filtered” list. scAR was performed twice for each sample, once with the unspliced matrices and once with the spliced matrices.

We performed RNA velocity on the LEPR stroma cluster. This included LEPR stroma from the 4 samples that made up the main bone marrow scRNA-seq dataset as well as LEPR stroma from an earlier validation experiment in which 2 femoral bone marrow scRNA-seq samples from 1 mouse were used (1 fixed with DES and 1 paired viable sample that was isolated using the mortar and pipette mechanical dissociation method). The 2 samples from the earlier validation experiment were preprocessed identical to that of the larger bone marrow scRNA-seq dataset described previously. The 6 samples were combined and scVI modeling was used for initial dimensionality reduction to isolate the combined LEPR stroma cluster. For this model, we subsetted the dataset on the top 1,000 highly variable genes using the Scanpy function highly_variable_genes() with the “flavor” parameter set to “Seurat_v3” and the “batch_key” parameter set to the sample identifier. For constructing the scVI model we used the denoised raw count matrix, 1 categorical covariate (the sample identifier), and 4 continuous covariates (natural log of total counts per cell, natural log of total genes per cell, and “G2M” and “S” scores). After UMAP computation and Leiden clustering, we then subsetted on the LEPR stroma and recalculated SCRAN normalized gene expression using the combined 6 samples with all cells assigned to the same cluster. We then performed scVI modeling again on the subsetted LEPR stroma dataset. This final model used 1,500 highly variable genes, 1 categorical covariate (the sample identifier), and 2 continuous covariates (“G2M” and “S” scores). We then recomputed UMAP dimensionality reduction and Leiden clustering for the final time. The Scanpy function ‘embedding_density()’ was then used to compute cell densities in UMAP space for viable and DES LEPR stroma.

velovi, a deep generative modeling framework for estimating RNA velocity (Gayoso et al., 2022b), was used for RNA velocity modeling. Briefly, denoised unspliced and spliced counts were normalized. To do this, we took the natural log+1 transformation of the denoised counts divided by the SCRAN size factor (previously calculated for each cell using the total expression matrix). Using the scVI latent space with the scVelo function ‘moments()’ using ‘n_neighbors’ set to 80, we then computed 1$^{st}$ and 2$^{nd}$ order moments for each cell across its nearest neighbors. We then subsetted the dataframe on the top 5,000 highly variable genes and used the velovi function preprocess_data() with ‘min_max_scale’ set to True, which scales the splicing expression matrices from 0 to 1, and with ‘filter_on_r2’ set to True, which filters out genes according to linear regression fit. From the velovi model, we extracted ‘velocity’, ‘fit_alpha’, ‘fit_beta’, and ‘fit_gamma’, and used the scVelo functions ‘velocity_graph()’ and ‘velocity_embedding_stream()’ to fit the smoothened velocity measurements onto UMAP space. We used the scVelo function ‘velocity_pseudotime()’ to calculate RNA velocity pseudotime and the scVelo function ‘paga()’ for trajectory inference.

**Analyses of Th17 T Cell intracellular CITE-Sequencing**

CellRanger count was used for demultiplexing, genome alignment, and antibody UMI counting. scAR was used to remove ambient RNA. Hashing cutoffs were determined visually using histograms. Doublets were defined as cells that were above the positive staining cutoff for 2 or more culture/panel-specific hashes or above the positive staining cutoff for both stim/unstim-specific hashes. Doublets were removed, as were cells that did not have any positive staining for a culture/panel-specific hash and/or a stim/unstim-specific hash. Quality control filtering was performed using a cutoff of 500 for total genes or total counts per cell and a cutoff of 0.1 was used for mitochondrial transcript proportion per cell. Preliminary UMAP dimensionality reduction and Leiden clustering revealed a small population of contaminating B cells (~7% of the total dataset), which were subsequently removed.

We calculated the median value of rabbit isotype counts per cell for each culture and stim/unstim combination (i.e. cultureA_stimulated, cultureA_unstimulated, etc; hereafter referred to as “samples”). For each sample, we then subtracted the median isotype value from the raw counts of the 2 phospho inCITE-seq antibodies per cell. If the subtraction resulted in a negative count for an inCITE-seq antibody in a given cell (i.e. if the median isotype value was greater than the counts for an inCITE-seq antibody), then we clipped the resulting value at 0. Median isotype counts showed little variation across samples, ranging from 3 to 5. The isotype corrected inCITE-seq counts were then natural log+1 transformed to give the final values shown in Figure 6. We annotated cells as positive for a given inCITE-seq phospho-target if a cell had >0 counts after isotype correction and natural log+1 transformation.

Lastly, we used scVI modeling to compute a latent space representation for UMAP dimensionality reduction. This model used 2,000 highly variable genes, 1 categorical covariate (the combined identifier for culture, panel, and condition [i.e. stim/unstim]), and 4 continuous covariates (natural log of total counts per cell, natural log of total genes per cell, and “G2M” and “S” scores). The resulting latent space was then transferred to the non-subsetted dataset, which was subsequently used to compute nearest neighbors followed by dimensionality reduction using the UMAP algorithm in Scanpy. Finally, SCRAN was used, with all cells assigned to the same cluster, to compute size factors for the normalization of gene expression.

**Th17 Single-Cell RNA-Sequencing: d0 versus d60**

Th17 scRNA-seq from paired DES-fixed cultures at d0 and d60 (with respect to storage in DES) were analyzed and preprocessed using the following methods. CellRanger count and Velocyto were used to compute total expression matrices and spliced/unspliced expression matrices, respectively. No ambient RNA removal was used for this analysis. Hashing cutoffs and doublet removal were performed as described previously. Quality control filtering was performed using a cutoff of 500 for total genes or total counts per cell and a cutoff of 0.1 was used for mitochondrial transcript proportion per cell. Contaminating B cells were removed. We then computed the number of genes and number of counts per cell using the 3 different gene expression matrices: total expression, unspliced expression, and spliced expression. Lastly, we used scVI modeling to compute a latent space representation for UMAP dimensionality reduction. This model used 2,000 highly variable genes, 1 categorical covariate (the combined identifier for culture, panel, and condition [i.e. stim/unstim]), and 4 continuous covariates (natural log of total counts per cell, natural log of total genes per cell, and “G2M” and “S” scores). The resulting latent space was then transferred to the non-subsetted dataset, which was subsequently used to compute nearest neighbors, followed by dimensionality reduction using the UMAP algorithm in Scanpy.

**Analyses of Flow Cytometry Data**

All flow cytometry data were analyzed using FloJo (Version 10.8).

**Statistics and Data Visualization**

Statistical analyses were performed using the R package rstatix or GraphPad Prism (version 6.01). Sample sizes and statistical tests are listed in the figure legends. All graphs with error bars report mean ± SEM values. Data were visualized using the R package ggplot2 (Wickham, 2011), with the exception of UMAP plots, which were generated using Scanpy. Correction for multiple hypothesis testing was applied where indicated.

### SUPPLEMENTAL FIGURES

**Supplemental Figure 1**. Paired Th17 T cell single-cell RNA-sequencing from day 0 and day 60 of DES fixation.

**Supplemental Figure 2**. Flow Cytometry gating scheme for viable and DES-fixed bone marrow.

**Supplemental Figure 3**. Bone marrow single-cell RNA-sequencing.

**Supplemental Figure 4**. Single-cell RNA-sequencing analysis of paired methanol-fixed and viable humans PBMCs from Katzenelenbogen et al. 2020.

**Supplemental Figure 5**. Cluster-wise linear regressions of paired viable and fixed samples.

**Supplemental Figure 6**. Integration of single-cell RNA-sequencing datasets for LEPR stroma analyses.

**Supplemental Figure 7**. Flow cytometry gating scheme and isotype controls for PFA/methanol and DES-fixed Th17 T cells.

**Supplemental Figure 8**. Antibody-oligonucleotide conjugation for intracellular CITE-seq.

**Supplemental Figure 9**. Extended data from Th17 intracellular CITE-seq.

**Supplemental Figure 10**. Extended data and isotype controls for IL2 flow cytometry in Th17 T cells.

**Supplemental Figure 11**. Comparison of three dissociation methods for viability and RNA recovery in CD45+ cells.

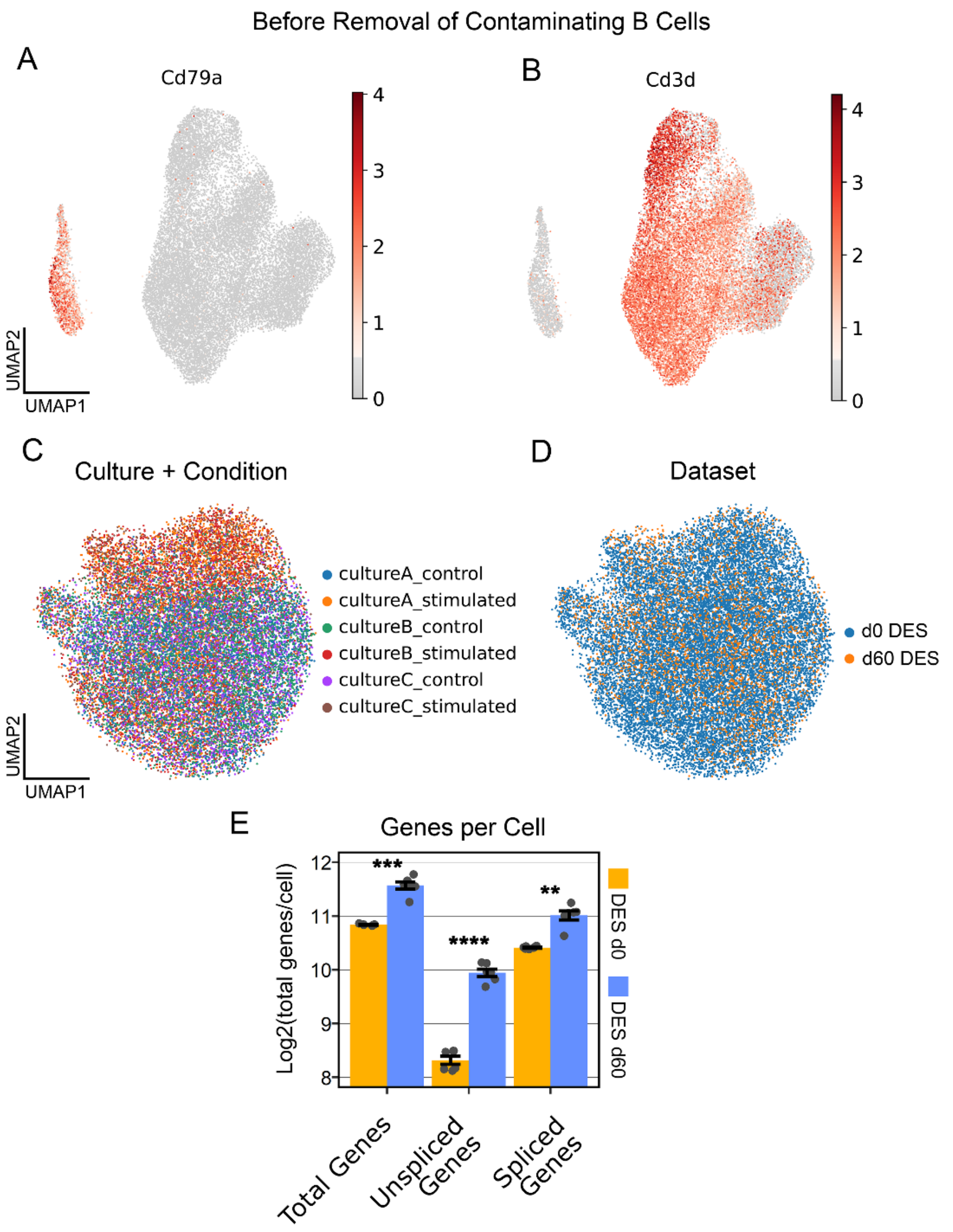


**Supplemental Figure 1. Paired Th17 T cell single-cell RNA-sequencing from day 0 and day 60 of DES fixation.** (**A-B**) Normalized expression of (**A**) Cd79a and (**B**) Cd3d in UMAP space showing contaminating B cells. (**C-D**) Integration of d0 and d60 datasets visualized in UMAP space showing labels for (**C**) combination of culture and condition and (**D**) dataset (i.e. d0 versus d60). (**E**) Mean genes per cell for each culture/condition combination (n=6) showing total genes, unspliced genes, and spliced genes by dataset (d0 versus d60). Two-way ANOVA with Bonferroni correction. ** $p \leq 0.01$; *** $p \leq 0.001$; **** $p \leq 0.0001$.

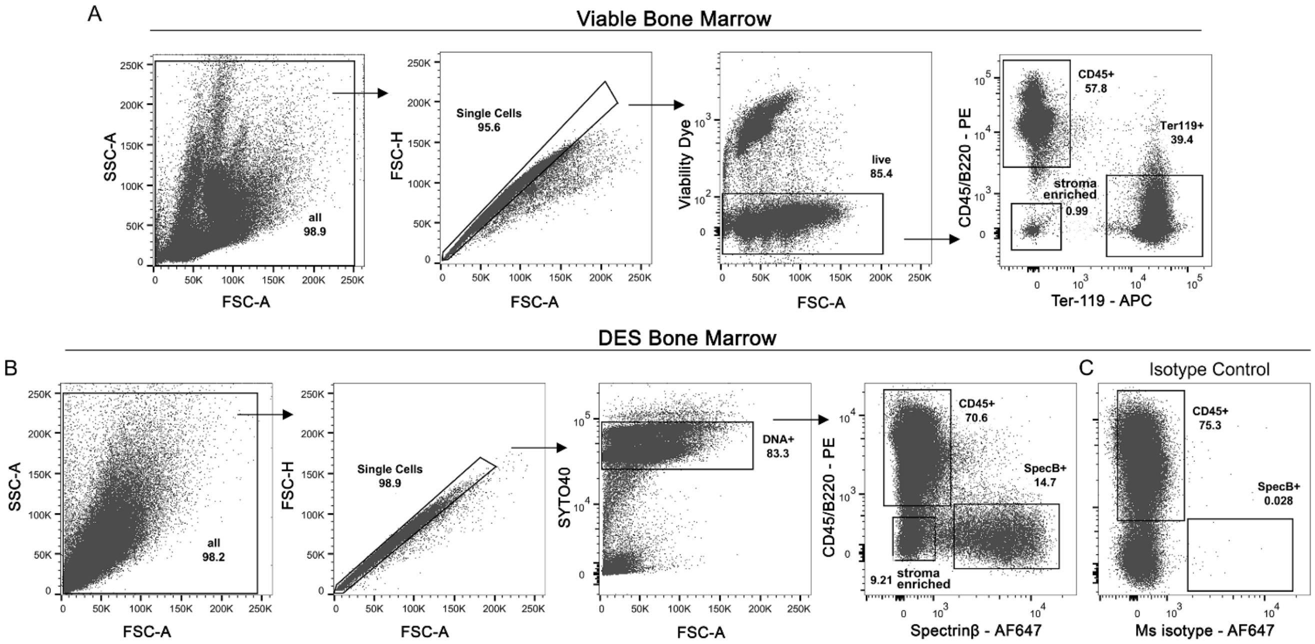


**Supplemental Figure 2. Flow Cytometry gating scheme for viable and DES-fixed bone marrow.** (**A**) Gating scheme for viable bone marrow. (**B**) Gating scheme for DES-fixed bone marrow. (**C**) Isotype control staining for Spectrinβ in DES bone marrow.

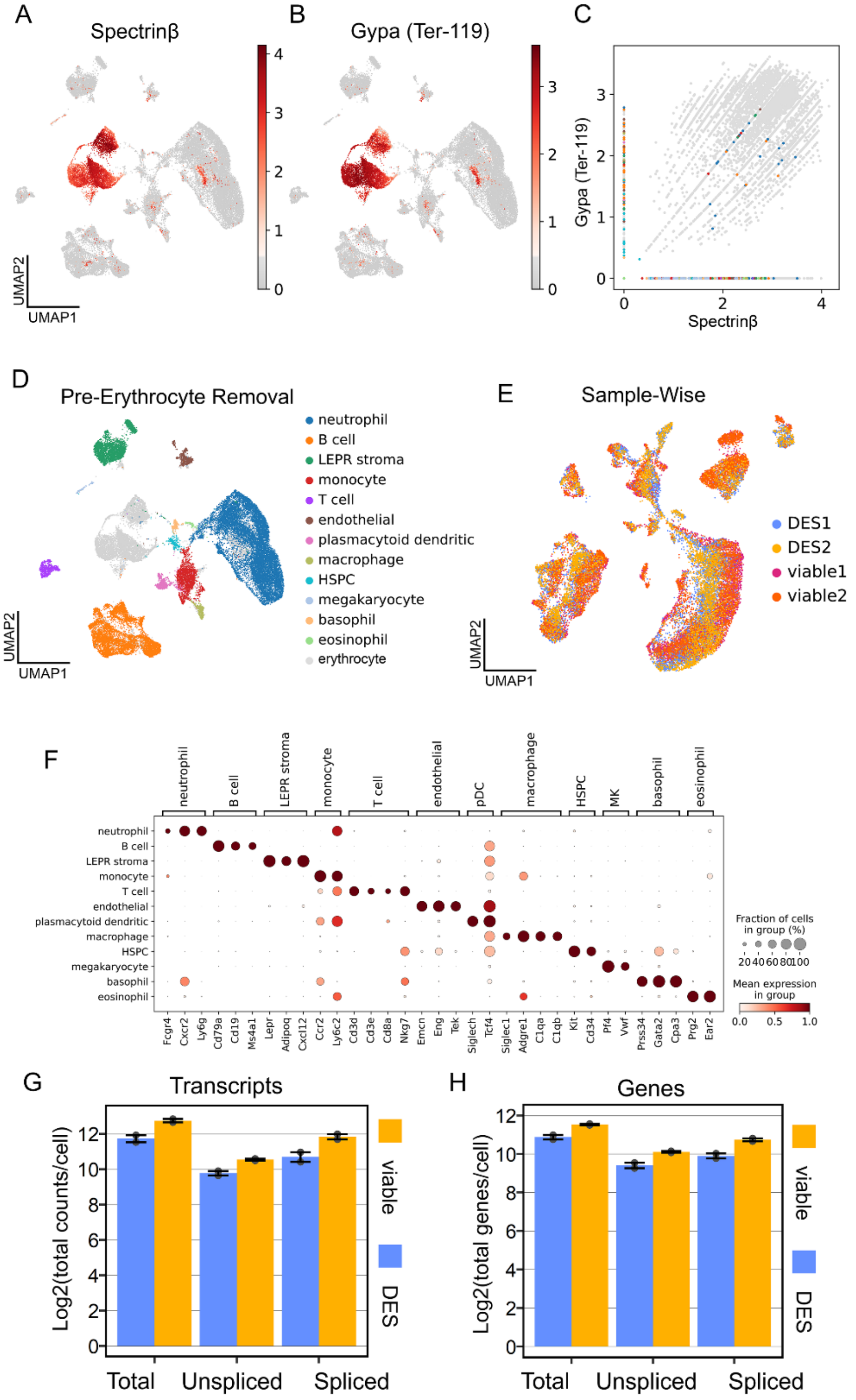


**Supplemental Figure 3. Bone marrow single-cell RNA-sequencing.** (**A-C**) Overlap in staining of (**A**) Gypa (Ter-119 antigen) and (**B**) Spectrinβ. (**C**) Scatter plot showing per cell correlations between Gypa and Spectrinβ. Color represents the cluster specific colors from (**D**). (**D**) UMAP representation before removal of contaminating erythrocytes. (**E**) Sample-wise labels of bone marrow scRNA-seq showing dataset integration. (**F**) Dot plot showing marker genes for each cluster. (**G-H**) Bar plots comparing mean total, unsliced, and spliced values per cell between DES and viable bone marrow for (**G**) transcripts per cell and (**H**) genes per cell.

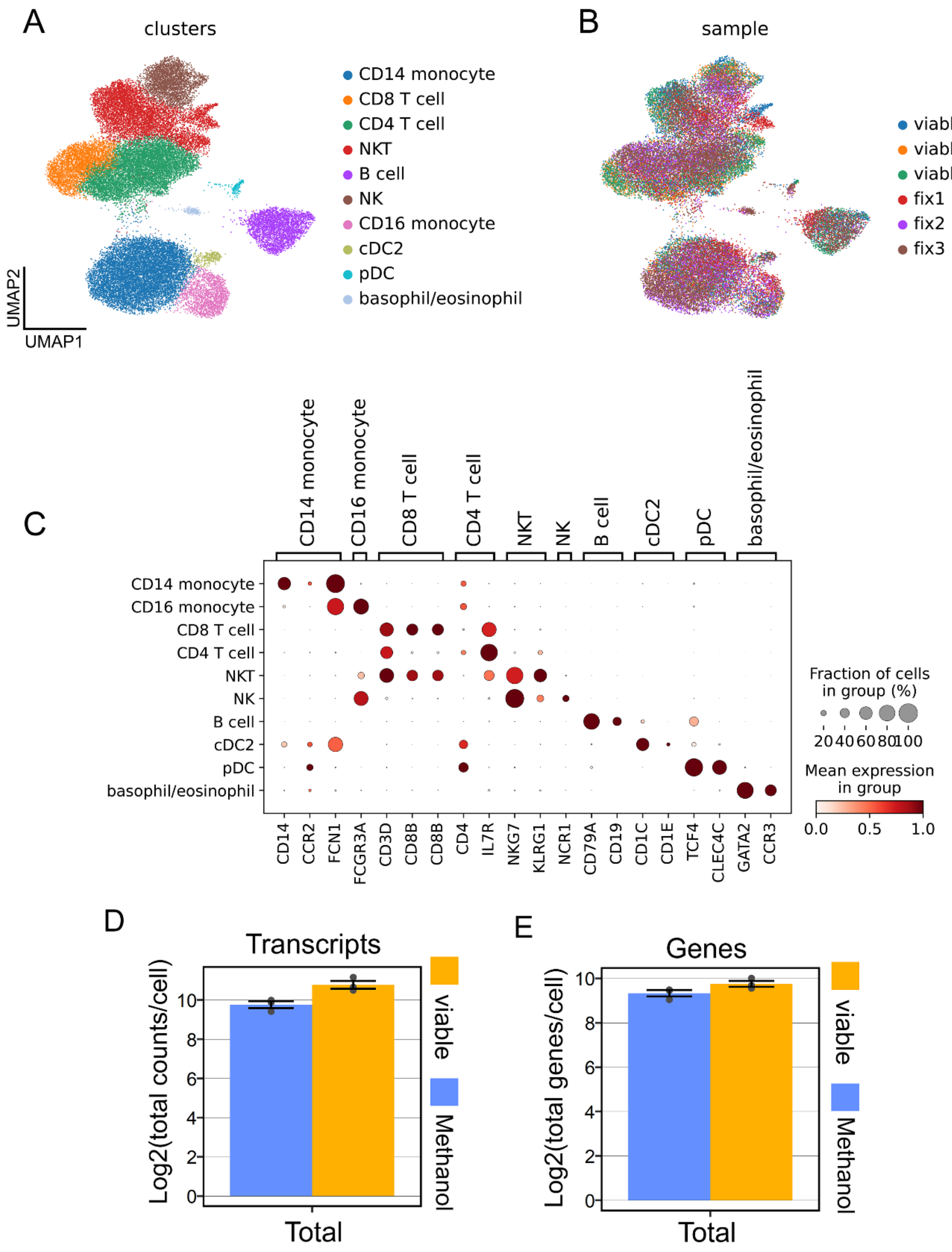


**Supplemental Figure 4. Single-cell RNA-sequencing analysis of paired methanol-fixed and viable humans PBMCs from Katzenelenbogen *et al.*, 2020.** (**A**) UMAP representation of 10 distinct cell populations. (**B**) Integration of methanol-fixed and viable PBMCs from 3 donors showing sample-wise labels in UMAP space. (**C**) Dot plot showing marker gene expression in the 10 clusters. (**D-E**) Bar plots showing mean (**D**) transcripts per cell and (**E**) genes per cell.

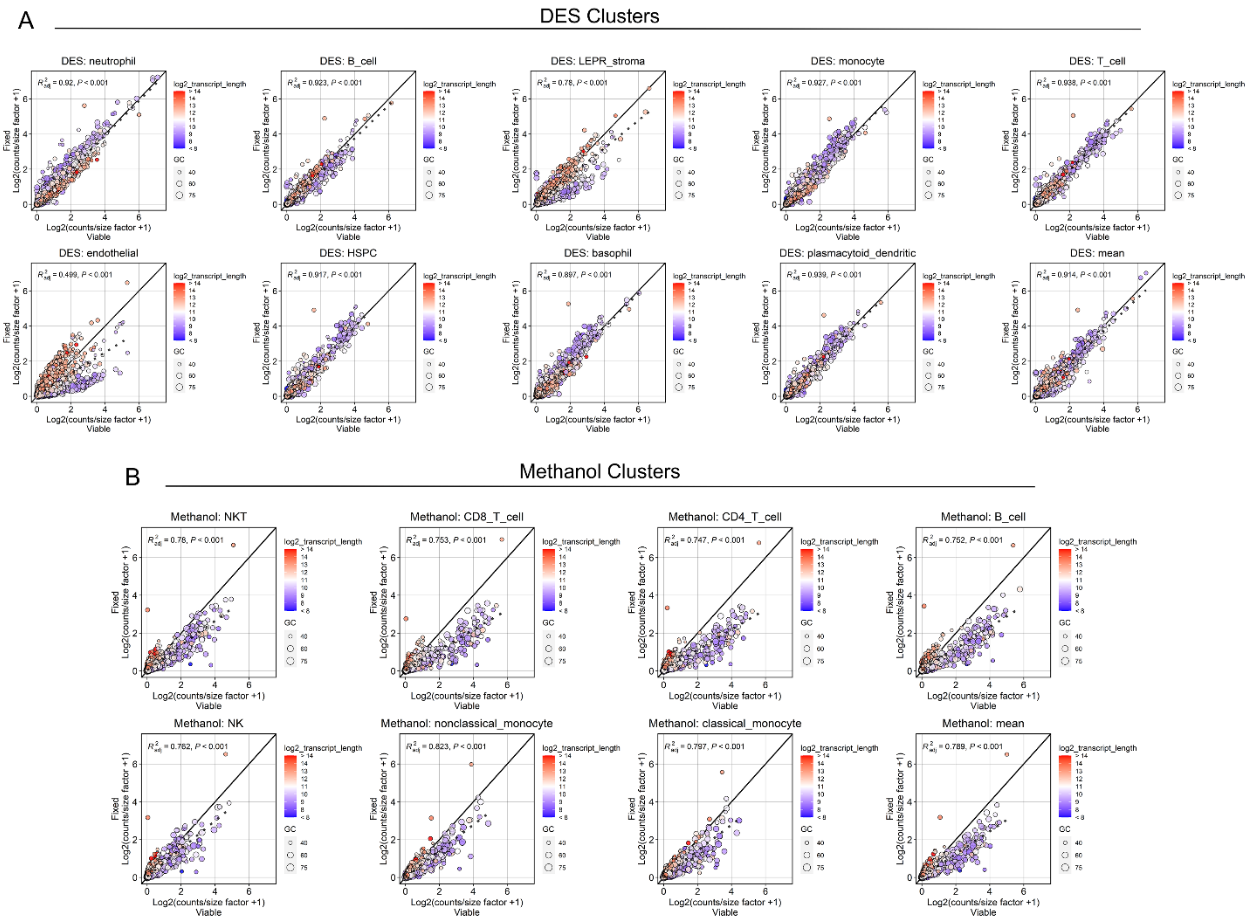


**Supplemental Figure 5. Cluster-wise linear regressions of paired viable and fixed samples.** (**A**) DES fixation in mouse bone marrow scRNA-seq. (**B**) Methanol fixation in human PBMC scRNA-seq.

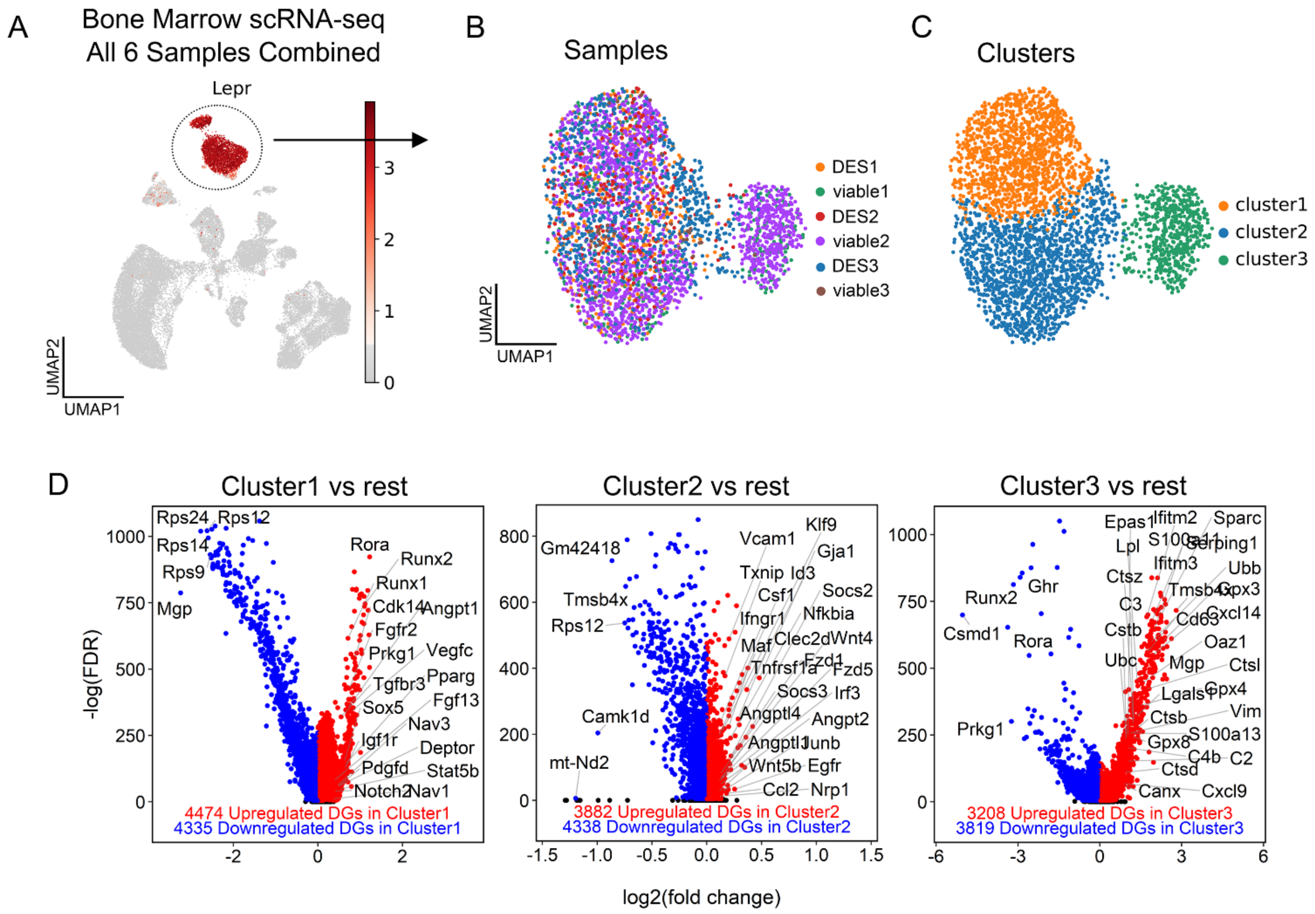


**Supplemental Figure 6. Integration of single-cell RNA-sequencing datasets for LEPR stroma analyses.** (**A**) UMAP representation of the combined dataset for n=6 samples (3 mice, paired DES-fixed and viable bone marrow). Lepr gene expression is shown and the circle indicates the cell population that was used for further analysis of LEPR stroma. (**B**) Sample-wise labels in UMAP space showing integration of all 6 samples for LEPR stroma. DES3 and viable3 are from the earlier validation dataset from one mouse. (**C**) Leiden clustering of the LEPR stroma showing 3 clusters. (**D**) Volcano plots showing differential gene expression in each of the 3 clusters versus the rest. Only viable cells were used for this analysis in order to understand LEPR stroma activation in dissociated viable stroma.

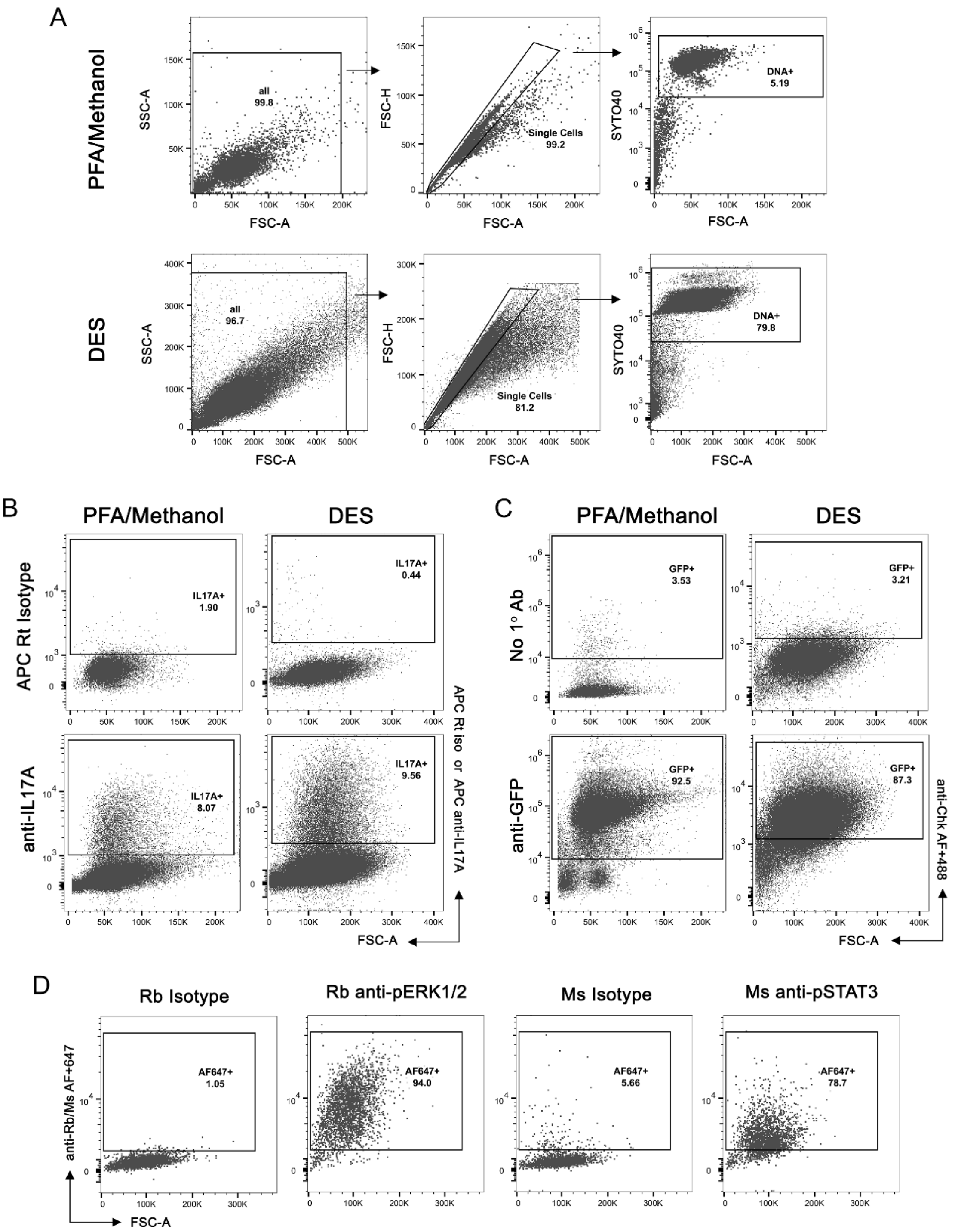


**Supplemental Figure 7. Flow cytometry gating scheme and isotype controls for PFA/methanol and DES-fixed Th17 T cells.** (**A**) Gating scheme for Th17 T cells fixed with either PFA/methanol (top) or DES (bottom). (**B**) IL-17A isotype controls for PFA/methanol and DES fixed cells. (**C**) anti-GFP negative controls for PFA/methanol and DES fixed cells. (**D**) Rabbit and mouse isotype controls for phospho-flow cytometry in DES fixed cells.

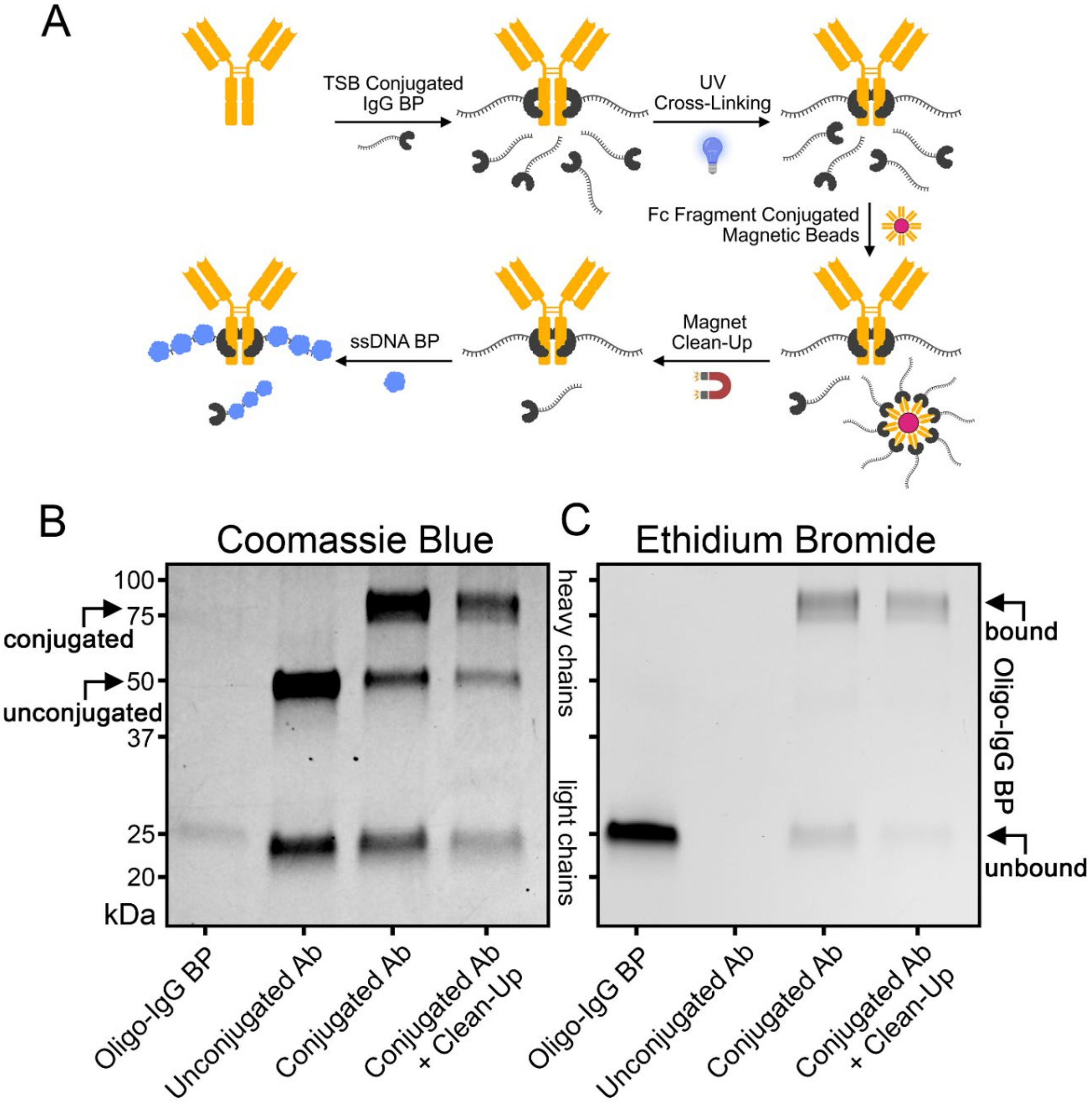


**Supplemental Figure 8. Antibody-oligonucleotide conjugation for intracellular CITE-seq.** (**A**) Schematic overview of antibody-oligonucleotide conjugation and preprocessing for inCITE-seq. (**B**) Polyacrylamide gel electrophoresis under denaturing conditions stained with Coomassie blue showing antibody conjugation. Each lane = 1µg of antibody. (**C**) The same gel from (**B**) stained with ethidium bromide showing bound and unbound oligonucleotide before and after clean-up with human IgG Fc magnetic beads.

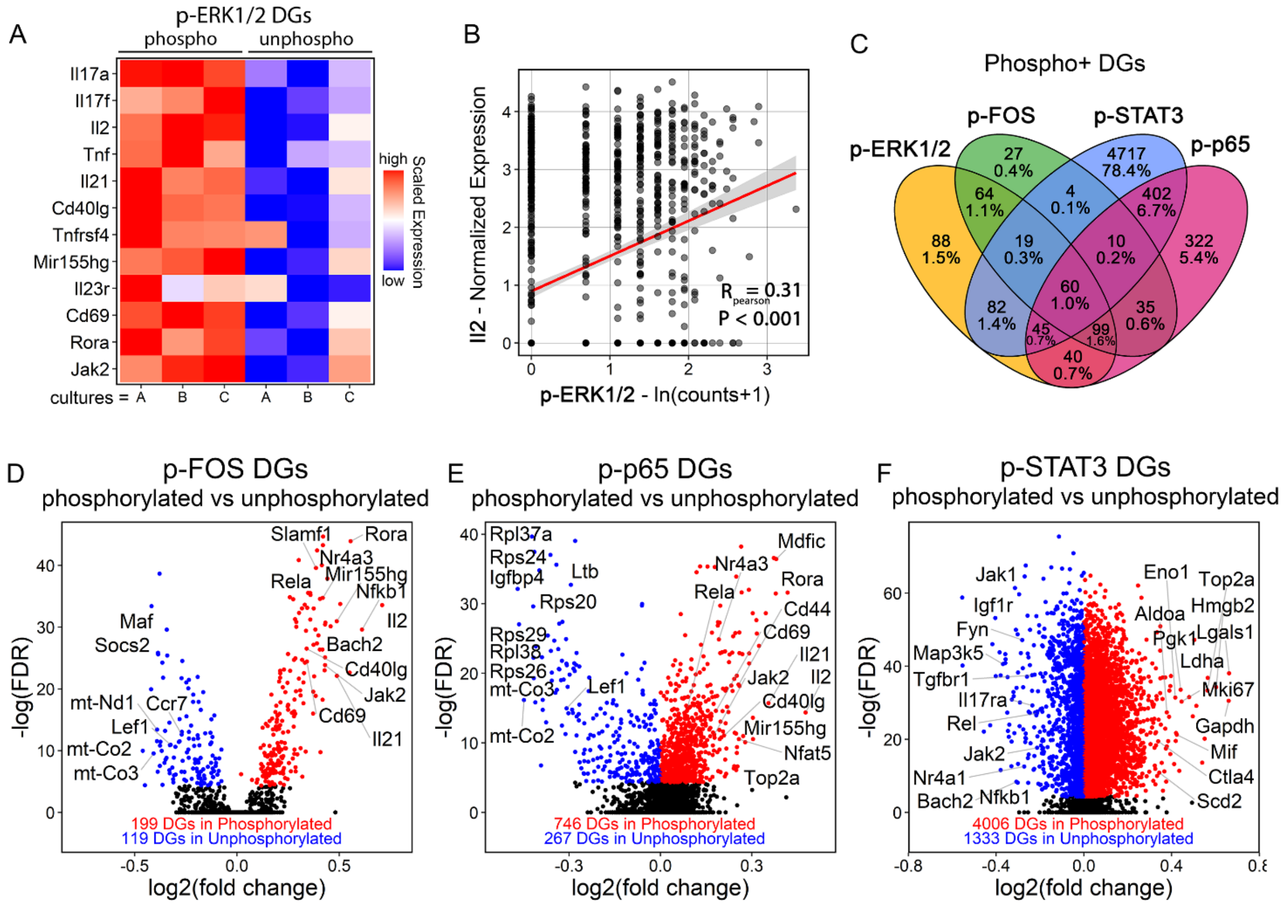


**Supplemental Figure 9. Extended data from Th17 intracellular CITE-seq.** (**A**) Heatmap showing scaled expression of selected p-ERK1/2 differentially expressed genes (DGs) by culture/condition. (**B**) Scatter plot showing Pearson correlation between IL2 expression and p-ERK1/2 magnitude. (**C**) 4-way Venn diagram showing overlap in DGs for p-ERK1/2, p-FOS, p-STAT3, and p-p65. (**D-F**) Volcano plots showing selected DGs for phosphorylated versus unphosphorylated (**D**) p-FOS, (**E**) p-p65, and (**F**) p-STAT3.

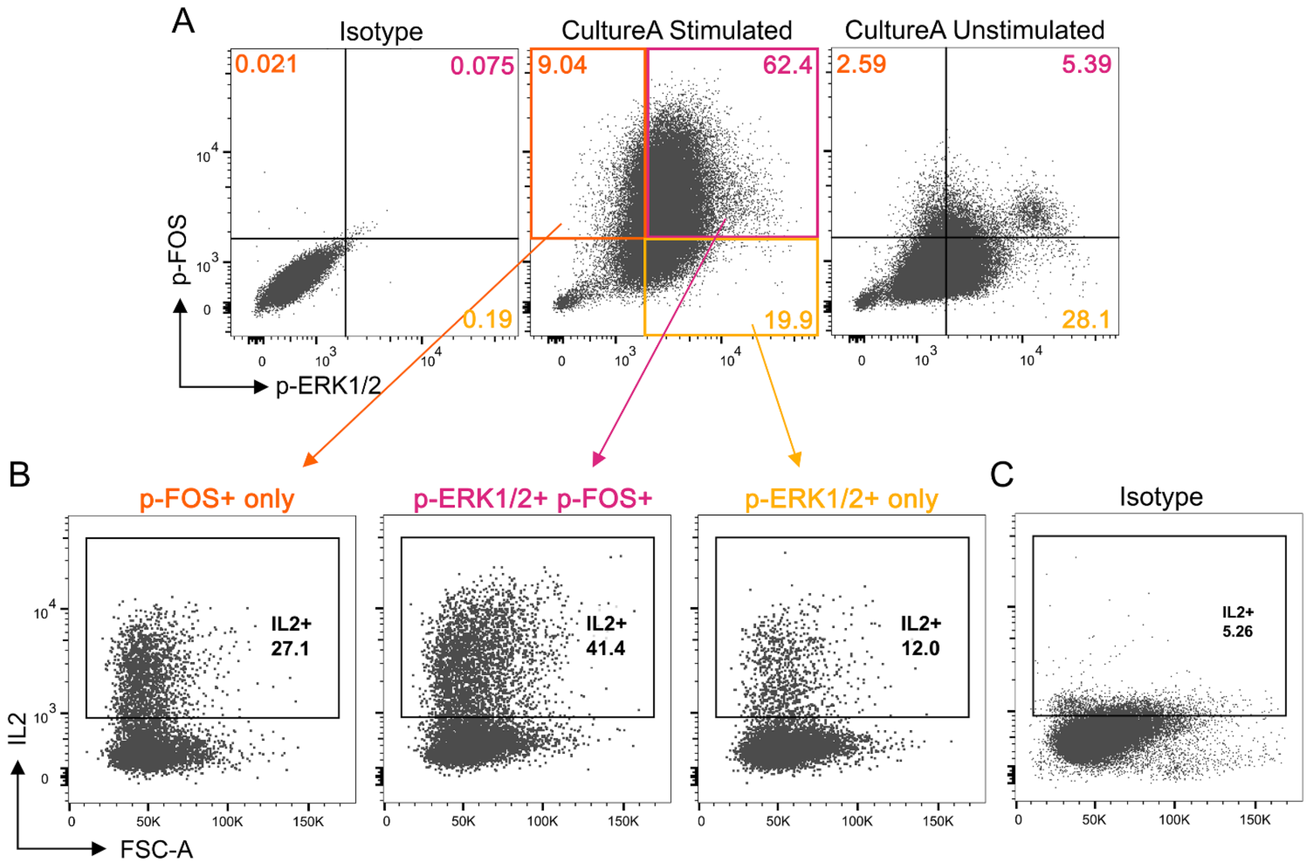


**Supplemental Figure 10. Extended data and isotype controls for IL2 flow cytometry in Th17 T cells.** (**A**) p-FOS and p-ERK1/2 gating scheme with representative examples. (Left) Alexa Fluor 647 isotype for p-FOS and rabbit isotype with anti-rabbit Alexa Fluor Plus 488 secondary for p-ERK1/2. (Middle) Representative stain for p-FOS and p-ERK1/2 after 3 hours of stimulation with PMA/ionomycin. (Right) Representative stain for p-FOS and p-ERK1/2 in the same culture as (middle) but without PMA/ionomycin stimulation. (**B**) IL2 staining in the three populations from (**A**, middle). The same number of events are shown for all three populations. (**C**) PE-conjugated rat isotype for IL2 gating.

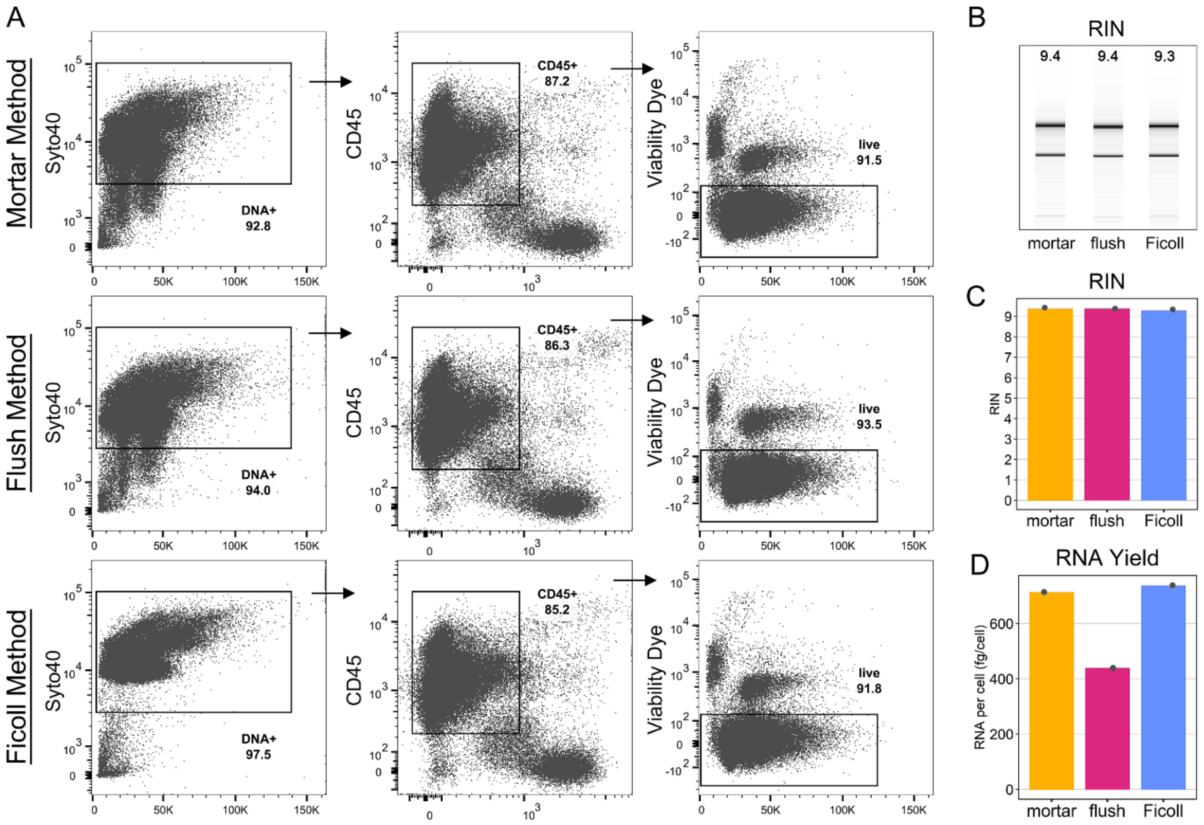


**Supplemental Figure 11. Comparison of three dissociation methods for viability and RNA recovery in CD45+ cells.** (**A**) Flow cytometry gating scheme showing viability staining in CD45+ cells using either the mortar-dissociation method (top), the bone marrow flushing method (middle), or centrifugation into Ficoll followed by dissociation via pipetting (bottom). (**B-D**) RNA analyses from 100,000 FACS sorted CD45+ bone marrow cells from the three dissociation methods. (**B-C**) Analysis of RNA integrity number (RIN) showing electrophoresis results (**B**) and RIN quantifications (**C**). (**D**) RNA yield per cell from Agilent 2100 estimates. n=2 mice; 1 femur per dissociation method.
